# Comprehensive benchmarking of somatic mutation detection by the SMaHT Network

**DOI:** 10.1101/2025.10.09.678885

**Authors:** The Somatic Mosaicism across Human Tissues Network, Alexej Abyzov

**Author notes:** A full list of SMaHT Network authors and their affiliations is available at the end of the article.

## Abstract

Somatic mosaicism is increasingly recognized as a fundamental feature of human biology, yet the detection of somatic mutations remains challenging. The SMaHT Network conducted four large-scale benchmarking experiments involving cell-lines and donor tissues, to evaluate sequencing technologies, experimental approaches, and computational methods for detecting different types of somatic mutations, generating community resource with >1,000× short-read and 100-400× long-read data for each of the nine analyzed samples. We determined effective strategies for utilizing short- and long-reads sequencing for mutation detection and demonstrated that using donor-specific assemblies and human pangenome improved calling, extending mutation catalogs to challenging genomic regions. We benchmarked six duplex technologies and showed that single-cell sequencing resolves cell type-specific mutational patterns and heterogeneity. Our results indicate that bulk, single-cell, and duplex analyses are complementary – and leveraging all three provides comprehensive characterization of mosaicism within tissues. Together, these findings provide a roadmap for accurate, genome-wide somatic mutation discovery and analysis.

## Introduction

Somatic mosaic mutations, i.e., post-zygotic genetic alterations, are increasingly recognized as important contributors to human health and disease beyond the context of cancer^1,2^. These mutations can arise during early development or in differentiated tissues, leading to cellular and genetic diversity that occurs naturally, but also has been shown to play important roles in neurodevelopmental disorders, cardiovascular diseases, autoimmune conditions, and age-related health decline^3–5^. Unlike cancer, mutations arising from normal biological processes are relatively rare, thus detecting low variant allele fraction (VAF) somatic mutations remains a major technical and analytical challenge, particularly due to low mutation burden and, consequently, low signal-to-noise ratios in the data.

Different sequencing approaches present distinct strengths and limitations for mosaic mutation detection. In bulk sample sequencing, the sensitivity to detect mutations is directly linked to sequencing depth, and, at standard genome coverage levels (e.g., 30-50×), low-VAF mutations are challenging to distinguish from sequencing errors or artifacts. Single-cell whole genome amplification (scWGA) followed by sequencing can detect private mutations in individual cells, but is limited by low throughput and often suffers from amplification noise, allelic dropout, and amplification biases, making it difficult to reliably detect mutations of all types genome-wide^6,7^. High-fidelity single molecule (also called “duplex”) sequencing has emerged as a powerful strategy for reducing sequencing noise and enabling accurate mutation detection but is currently constrained to mostly quantifying average mutation burden and spectrum across cells in a sample or to deep profiling of specific targeted loci^8^, rather than genome-wide mutation discovery. Compounding these technical limitations unique to each sequencing modality is the absence of standardized analytical approaches for calling low VAF mutations or annotating/interpreting their functional impact. Only a handful of methods have been developed specifically to detect low VAF somatic mosaic mutations, and these methods face trade-offs between sensitivity, specificity, runtime, and the ability to detect more complex mutation types, such as structural variants (SV) or tandem repeat changes. Importantly, robust benchmark and control human data to evaluate and improve computational tools or experimental assays are lacking in the field, further hampering progress in our understanding of how somatic mosaicism contributes to development, disease, and aging. As such, establishing rigorous experimental and analytical standards, and generating gold-standard datasets are essential to enable accurate and reproducible detection of somatic mutations that may be clinically relevant.

Cancer consortia have developed powerful resources for somatic mutation evaluation^9,10^, but these efforts were primarily focused on detecting mutations with higher VAF (>5%), often relying on paired analysis of tumor and matched normal tissues, limiting the applicability for detecting low VAF mutations in normal tissues without a “matched normal” comparator sample. Other benchmark efforts by the Brain Somatic Mosaicism Network (BSMN)^11,12^, the Genome in a Bottle (GIAB) consortium^13^, and Ha et al.^14^ developed important insights and datasets for evaluating somatic mutation detection. But these benchmark studies were focused on evaluating limited mutation types, i.e., SNVs and mobile element insertions (MEIs), with relatively high VAF (typically over 3%) or mutations in exonic regions. Furthermore, to the best of our knowledge, rigorous benchmarking of duplex technologies and scWGA has not been reported so far.

The Somatic Mosaicism across Human Tissues (SMaHT) Network aims to map somatic genetic diversity across different tissues and cells within individuals. In the initial phase of the SMaHT, benchmarking efforts were made to address aforementioned gaps, expand and compare assessment capabilities, enable comprehensive detection of somatic mutations from using short- and long-read bulk whole genome sequencing (WGS), inform the project’s production phase, and support efforts from the broader scientific communities in development and validation of tools that identify low VAF (<5%) SNVs, indels, MEIs and SVs. Here we describe key results of benchmark studies presented in a collection of papers from the Network^15–23,24–31^, outline the datasets and resources generated from the benchmark experiments in the first phase of the SMaHT Network which are publicly available for research communities, and provide conceptual and practical recommendations for the discovery of somatic mutations of various types across different experimental and sequencing contexts. To establish a common framework for consistency and clarity, we define terminology used in our studies and list key organizational entities of the SMaHT Network (**Box 1, Table S1**).

### Box 1.

**Terminology utilized and key organizational entities of the SMaHT**

**Somatic mutation:** a mutation that is acquired in a person’s lifetime, rather than inherited from their parents. Also known as post-zygotic mutations (PZM) or mosaic mutations.

**SNV:** Single nucleotide variant, also sometimes referred to as single base substitution (SBS), are the mutations of one base into another, such as C>T.

**Indel:** Short insertions or deletions of DNA sequence, by convention fewer than 50 base pairs in length.

**SV:** Structural variant, mutations that involve large parts (over 50 bases) of the genome and may include large deletions, tandem duplications, inversions and translocations between chromosomes.

**MEI:** Mobile element insertion, a special class of SVs that involve the integration of retrotransposons into the genome.

**sSNV, sIndel, sSV, sMEI:** prefix ‘s’ is often used to emphasize that mutations are somatic.

**VAF:** Variant Allele Fraction, the ratio of sequencing reads supporting a specific mutation and the total number of reads covering the mutant site. It can be used to deduce the fraction of cells harboring a specific mutation.

**Mutation burden/load:** the total number of mutations (of a given class) in a tissue sample or cell.

**DSA:** Donor-Specific Assembly, complete or near telomere-to-telomere haplotype assembly for a tissue donor.

**Benchmarking or reference mutation set:** a set of mutations that exists in a tissue, as supported by orthogonal evidence.

**Single-molecule or duplex sequencing:** dual-DNA-strand, ultra-low error rate sequencing approaches that randomly sample mutations in bulk DNA to estimate average mutation burdens and signatures across cells.

**Primary template-directed amplification (PTA):** a whole genome amplification technique that in the context of this benchmarking exercise was applied to the analysis of genomes of individual cells.

**Easy genomic region:** intervals in the 1000 Genomes Project strict accessibility mask, representing confidently mappable, consistently callable loci.

**Difficult genomic region:** loci in PanMask pm151 (strict) but excluded from the 1000 Genomes Project strict mask, representing moderately mappable regions with elevated short-read error/ambiguity.

**Extreme genomic region:** the remainder of the genome outside both masks, enriched for highly repetitive or structurally complex sequence.

**<u>Key organizational entities</u>**

**GCC:** genome characterization center – a center that is primarily focused on generating sequencing data.

**TTD:** tools and technology development center – a center that is primarily focused on developing and testing experimental technologies and analytical tools for discovering somatic mutations.

**TPC:** tissue procurement center – a center that is focused on collecting, processing, and distributing samples from post-mortem donors.

**OC:** organization center – a center that organizes events and resources involving consortium participants.

**DAC:** data analysis center – a center that is focused on data aggregation, processing, analysis, and release.

### Results

#### Benchmark Design

To address the challenges of benchmarking the detection of diverse somatic mutation types and advance method development, we designed and carried out four benchmarking experiments: 1) “HapMap”, using a mixture of 6 well-established lymphoblast cell lines; 2) “COLO829”, well-characterized cancer and matched normal cell lines mixed at a known ratio; 3) a primary fibroblast sample with subclonal structure and matched clonal iPSC lines, and 4) six intact tissues or homogenate tissue samples from four phenotypically normal *post-mortem* donors (**Fig. 1**). These experiments were designed to comprehensively address the gaps in benchmark efforts that a single experiment alone could not address otherwise (**Table S2**), creating a constellation of experiments that generated deep, complementary datasets enabling a comprehensive benchmarking effort (**Table 1**).

**Figure 1.**
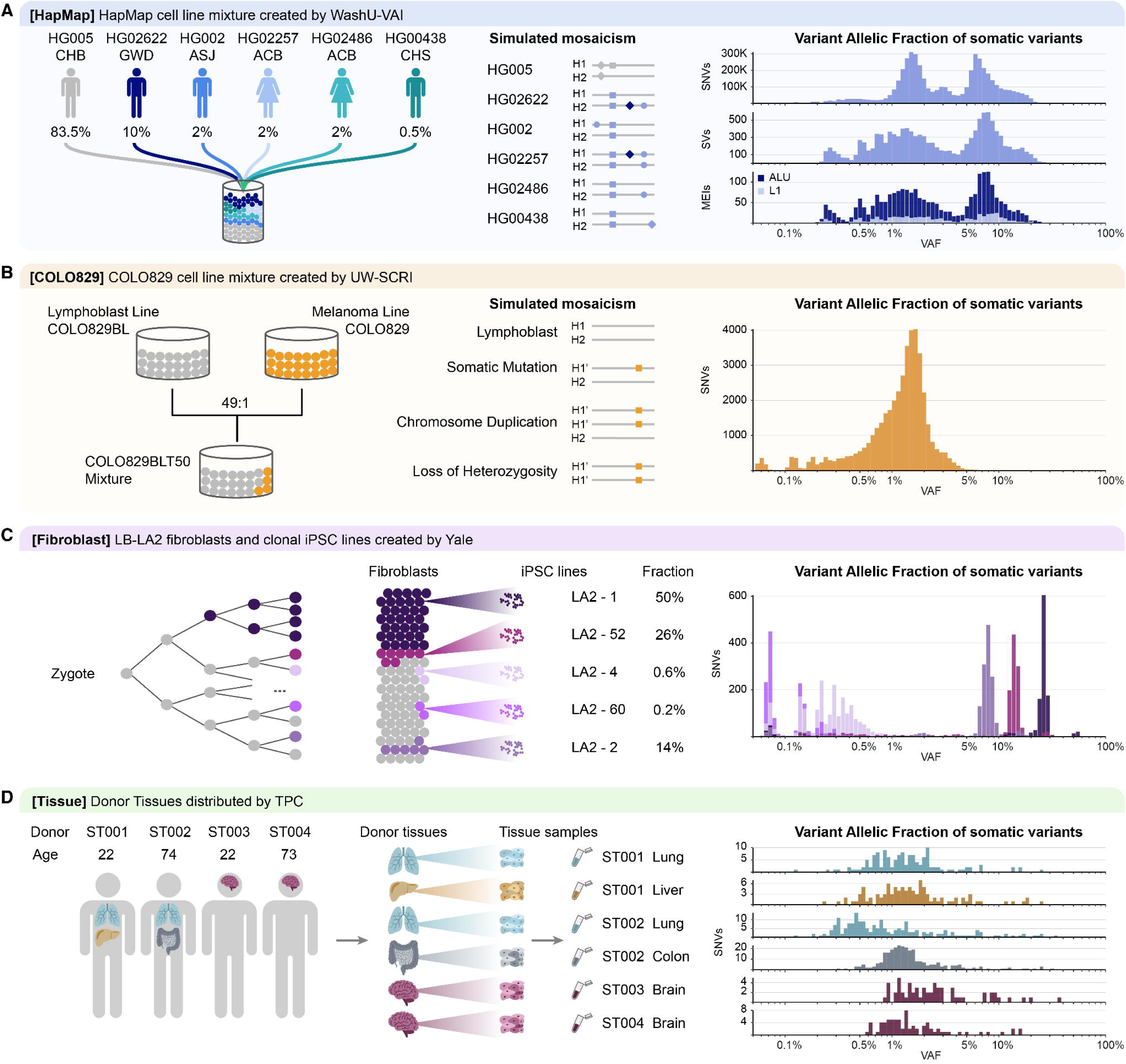
SMaHT benchmarking experiments. Four benchmarking experiments were carried out by the SMaHT Network. **A)** In the “HapMap” experiment, six well-characterized HapMap cell lines from donors with various ethnic backgrounds (CHB = Chinese; GWD = Gambian in Western Division, Mandinka; ASJ = Ashkenazim Jewish; ACB = African Caribbean in Barbados; CHS = Chinese Han in the South) were mixed at known proportions. Germline variants present in some (but not all) lines simulated somatic mutations (SNVs, indels, SVs, and MEIs) at VAFs ranging from 0.25% to 16.5%. The resulting sample has a mix of 12 haplotypes. **B)** In the “COLO829” experiment, well-characterized lymphoblastoid and melanoma lines from the same donor were mixed at a 49:1 ratio. Mutations in the melanoma line simulated SNVs and indels at VAF up to 5% in the mix. **C)** In the “Fibroblast” experiment, a sample of primary fibroblasts, which has natural mutations accumulated during the lifetime of the donor, was analyzed. Five clonal iPSC lines were derived from the fibroblasts, allowing the inference of somatic SNVs and indels at VAF from 0.1% to 25% and revealing the clonal structure in the fibroblasts. **D)** In the “Tissue” experiment, six frozen tissues obtained *post-mortem* from four phenotypically normal donors were analyzed to benchmark the detection of somatic mutations in production-like samples. Intact tissue samples as well as homogenized samples were used depending on experimental assay types.

**Table 1.** Summary and comparison of benchmarking experiments and reference mutation sets conducted by the SMaHT Network. The reference mutation sets from four donor tissues were inferred from the analysis of bulk whole genome sequencing data. MEI counts ranged depending on whether they were classified based on repeat annotation (upper bound) or mechanistic signature of target primed reverse transcription (lower bound). Certain variant types were not analyzed in a few samples as denoted by ‘-’. *For most analyses homogenized tissue samples were used to ensure uniformity across analytical groups, however, single cells were also isolated from the intact tissue. *estimated

|  | **HapMap mixture** | **COLO829BLT50** | **Fibroblasts** | **Donor tissues** |
| --- | --- | --- | --- | --- |
| **Center for sample preparation** | WashU-VAI | UW-SCRI | Yale | TPC |
| **Mutation characteristics** | Germline based | Cancer based | Natural | Natural |
| **Haplotypes in a sample** | 12 | 2 | 2 | 2 |
| **SNVs** | ~5,000,000 | 49,377 | 11,227 | 850* |
| **Indels** | ~1,800,000 | 2,255 | 2,718 | - |
| **MEIs** | 3,066-10,607 | 1 | - | 21 |
| **SVs/CNVs** | 34,140 | ~100 | a few | - |
| **Reference mutation sets from clonal line** | Yes | Yes | Yes | No |
| **Mutation frequencies** | 0.25%-16.5% controlled | 0.1%-5% controlled | 0.1%-25% uncontrolled clonal structure | variable |
| **Haplotype assemblies available** | Yes | Yes | Yes | Yes |
| **Sample condition** | *In vitro* grown | *In vitro* grown | *In vitro* grown | *Post-mortem snap frozen\** |
| **Terms for material and data distribution** | Unrestricted | Unrestricted | Protected | Protected |

A common way of simulating somatic mosaicism is physical or *in silico* mixing of samples with well-characterized genomes and with known germline variants. Such germline variants shared across a few, but not all mixed genomes will simulate somatic mutations of different frequencies in the mix (**Fig. 1A**). HapMap cell lines are particularly suitable for this purpose, as they have been extensively characterized using a wide array of genomic technologies, providing comprehensive, complementary genomic data suitable for benchmarking efforts. Additionally, donors of HapMap samples provided broad consent for future research uses, enabling extensive genotyping, sequencing, and various genetic variation investigations. The SMaHT Network has leveraged six genetically diverse diploid cell lines with near telomere-to-telomere (T2T) genomes assembled by the Human Pangenome Reference Consortia (HPRC) (**Fig. 1A**). These cell lines were combined at predefined ratios to simulate large numbers of somatic SNVs, indels, MEIs, and SVs at VAFs ranging from 0.25% to 16.5%, and the accuracy of the mixture was confirmed using droplet digital PCR (ddPCR). The availability of near T2T genome assemblies enables us to benchmark mutation detection beyond the constraints of a standard reference genome. However, as the simulated somatic mutations were derived from germline variants, their properties (distribution across genome, mutation spectra, etc.) may differ significantly from genuine somatic mutations observed in natural biological contexts. Also, the simulated mutations are present on multiple haplotypes rather than just two as in case of real biological samples and this discordance likely compromises mutation calling when using phasing information (e.g., with long reads) and limits the ability to benchmark somatic mutation calling in genomic regions where large stretches of DNA differ between individuals (e.g., satellite arrays). That led us to carry out further benchmark experiments in which all cells have the same two original mutated haplotypes.

To address this, the SMaHT Network created a publicly available benchmarking resource using an established hypermutated melanoma cancer cell line (COLO829) and a paired normal lymphoblastoid cell line (COLO829BL) derived from the same individual (**Fig. 1B**). For this benchmark sample, we generated a nearly complete T2T assembly and used deep short-read and long-read sequencing of both lines to obtain a set of somatic cancer mutations in the COLO829 line. We then created a benchmark sample (COLO829BLT50) from the COLO829BL and COLO829 cells mixed at a defined ratio (49:1), and the accuracy of the mixture was confirmed using ddPCR. Aneuploidies in the COLO829 cell line enabled this mixture resource to simulate mutations with VAFs ranging from 0.1% to 5%. However, this approach was limited as the COLO829 somatic mutations are largely derived from a single mutational process (i.e., UV damage), which lacks MEIs; and this cell mixture product is confounded by cell culture-derived clonal mutations in both the COLO829 and COLO829BL cell lines.

To provide an opportunity for benchmarking the discovery of genuine somatic mutations in non-tumor tissue and across a wide range of allele frequencies, we used a sample of primary human fibroblasts from a skin biopsy. The fibroblast sample was known to have clonal structure with five clones represented by five single fibroblast cell-derived induced pluripotent stem cell (iPSC) lines^32^. We used these iPSC lines to derive a true-positive set of mutations present in the fibroblasts at VAF from 0.1% to 25% (**Fig. 1C**). However, there were no somatic MEIs and only a few somatic SVs in the fibroblasts, making this approach suitable for only testing discovery of somatic SNVs and indels.

All aforementioned benchmark experiments used cell lines, which, being grown *in vitro*, are not likely to fully reflect all challenges when dealing with real human tissues. Therefore, the Network conducted benchmarking experiments using *post-mortem* human tissue from four donors (**Fig. 1D**) to test feasibility of mutation discovery in DNA and nuclei isolated from frozen tissue samples that are equivalent to samples in the production phase of the SMaHT Network. Equally important, mutation discovery in real human tissues provided the opportunity of assessing to what extent mutation heterogeneity across tissues could be captured using various sequencing approaches including bulk WGS^15–21^, duplex sequencing^22^, and single cell analyses by primary template-directed amplification (PTA)^23^. As the genuine set of somatic mutations in the human tissues is not known, we used orthogonal evidence from different analytical approaches and sequencing platforms to assess the accuracy of mutation detection.

For each experiment, a single laboratory prepared homogenized tissues (or suspended cells) and distributed samples to five Genome Characterization Centers (GCCs) and other groups in the Network (**Table S1**). Tissue homogenization and cell suspension ensured that samples with the same genomic content were analyzed across the Network for consistency and robustness of data generation. For a few assays (including PTA benchmarking) intact tissue samples processed at different sites were also used to generate the data.

#### SMaHT Benchmarking Data Generation

To comprehensively benchmark somatic mutation detection, we established a set of ‘core’ and ‘extended’ assays that were applied to the benchmark samples (**Fig. 2A**). The ‘core’ assays consisted of deep short- and long-read bulk WGS and RNA sequencing (RNA-Seq) across multiple sequencing platforms, aimed to identify moderate to high VAF mutations that arise during early development or during clonal expansions later in life. In contrast, the ‘extended’ assays aimed to discover ultra-low VAF somatic mutations (e.g., duplex-sequencing and single cell sequencing), infer cellular lineages with single-cell WGS, and profile hard-to-map genomic loci using long-read-based variant call validation and haplotype-resolved diploid genome assemblies (e.g., Hi-C, ultra-long ONT sequencing). Five GCCs and multiple TTDs (**Table S1**) applied these assays in the benchmarking experiments, enabling us to evaluate the impact of center-to-center variation in sequencing data on the robustness of somatic mutation detection, and to assess the utility and complementarity of the different core and extended assays for detecting the full spectrum of somatic mutation types (**Tables S3, S4**).

**Figure 2.**
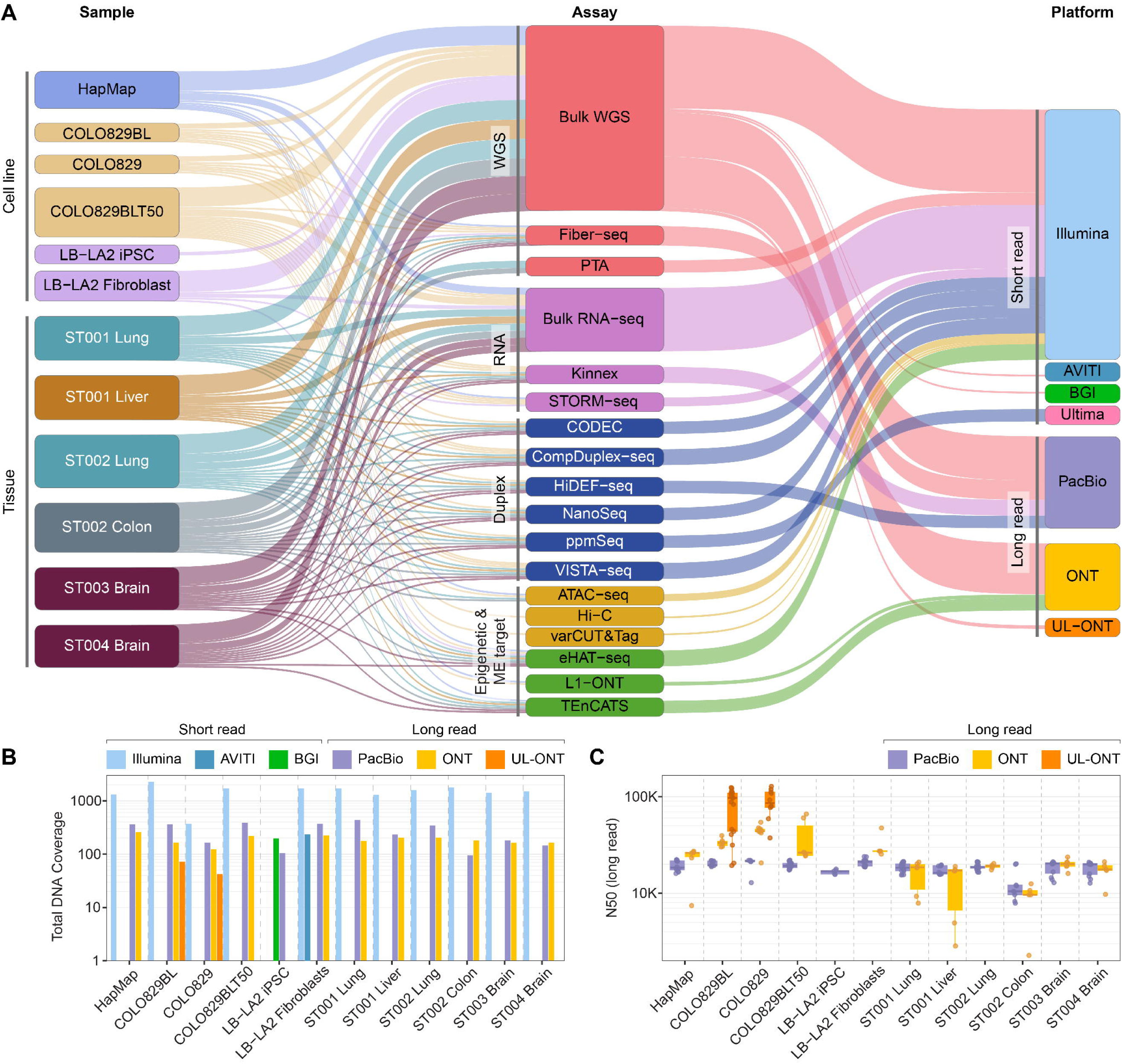
Summary of the SMaHT benchmarking data. **A)** The Sankey plot describes the cell line and tissue samples, experimental assays, and sequencing technologies applied in the benchmarking experiments. The width of connecting lines between columns reflects the number of samples from which data were generated from the corresponding experimental assays and sequencing technologies across multiple centers (**Table S1)**. Total sequencing coverages of WGS datasets using all short-read and long-read sequencing technologies applied in the benchmarking experiments are shown in **(B)**, with N50 as a representative data quality metric for PacBio and ONT long-read sequencing in **(C)**. Details of data generation are provided in Tables S3 and S4.

All sequencing data and metadata were submitted and systematically processed in a uniform manner by the Data Analysis Center (DAC) to ensure consistency and quality (**Methods; Fig. S1**). This included the removal of platform-specific sequencing artifacts (e.g., poly-Gs), alignment to a specific version of the GRCh38 reference genome, and reassignment of read group identifiers that follow the standard nomenclature to maintain consistency of BAM files across multiple sequencing centers and assay types. Each sample was checked for human and microbial contamination, sample mislabeling or swaps, as well as library qualities (e.g., DNA insert sizes for short paired-end sequencing or read lengths for PacBio/ONT long-read sequencing), sequencing and alignment qualities (e.g., aligned base mismatch, mapping rates) at the DAC where the data submitted from GCCs are uniformly processed and analyzed (**Table S5**). In addition, individual standard sequencing QA and QC were performed at GCCs where the data were generated. For RNA-Seq, standard metrics, such as 5’ and 3’ coverage ratios and library complexity checks, were performed. In addition, the quality, contiguity and completeness of the haplotype-resolved diploid genome assemblies were also checked using multiple approaches. Critical QC check results, such as human and microbial contamination, are tracked as part of metadata of the files accessible from the SMaHT Data Portal and public data repository.

In total, the benchmarking experiments produced over 90 Tb of raw sequencing data across the core and extended assays (**Table S4**). For each benchmarking sample, we generated deep WGS data with >1,200x short-read sequencing genome coverage (1,712× for COLO829BLT50, 1,323× for HapMap mixture, 1,706× for fibroblasts, and 1,297-1,773× for tissue samples), >300× long-read PacBio HiFi sequencing genome coverage (390× for COLO829BLT50, 363× for HapMap mixture, 370× for fibroblasts, and 95-345× for tissue samples), >200× long-read ONT sequencing genome coverage (218× for COLO829BLT50, 259× for HapMap mixture, 224× for fibroblasts, and 163-204× for tissue samples; **Fig. 2B,C**), and 72-299 million read pairs from polyA-enriched and total RNA-Seq. We also generated short- and long-read sequencing data from parental (COLO829 and COLO829BL) or derived (iPSCs) samples to aid in the discovery of benchmarking mutation catalogs. In addition, more than a dozen extended assays were applied to the benchmarking material, including various duplex sequencing approaches, single-cell sequencing methods, targeted sequencing of mobile elements, and chromatin profiling techniques (**Fig. 2A**). These assays encompassed duplex sequencing of donor tissues and umbilical cord blood, as well as whole-genome sequencing of 102 single cells at coverage depths ranging from 25× to 60×. For easy access of the benchmarking data to the scientific community, the Data Analysis Center built the SMaHT Data Portal (https://data.smaht.org) and the benchmarking data are registered under dbGaP (**Table S6**; **KEY RESOURCE TABLE**).

#### Benchmarking experimental and computational methods for somatic SNV and indel discovery

We first sought to benchmark the performance of somatic SNV and indel detection using the COLO829BLT50 benchmarking material^15^ (**Fig. 1B**; **Table 1**). Across five Illumina WGS datasets generated from different sites (170×–470×), 49% and 16.7% of the VAF <1% and 1-3% COLO829-specific SNVs were not observed in at least one dataset, highlighting the experimental challenges associated with low-VAF (<3%) somatic mutation detection even under well-controlled conditions (**Fig. 3A**). For robust detection of mutations with VAF less than 2%, we observed that bulk WGS with coverage greater than 300× was required^15^.

**Figure 3.**
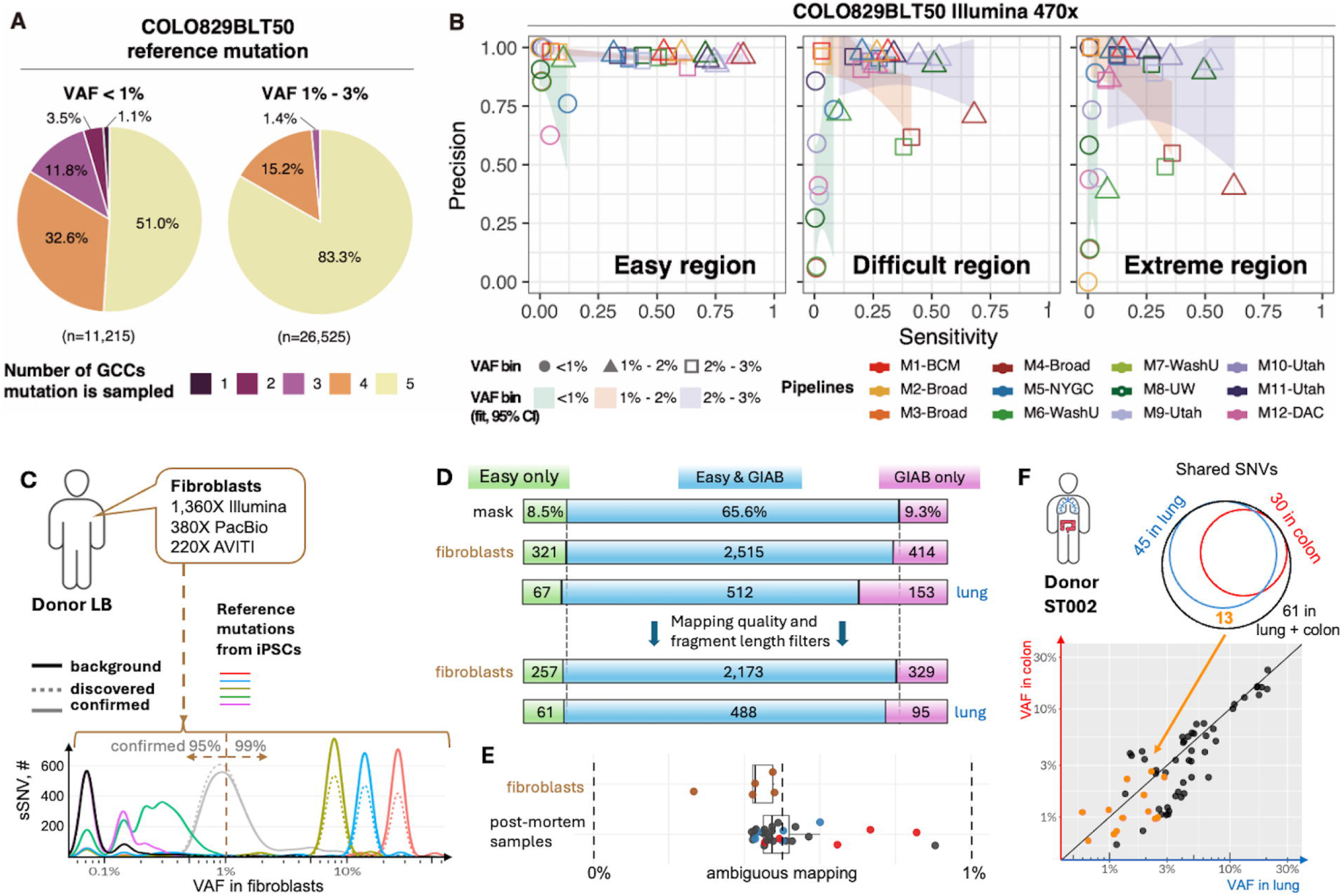
Multi-platform discovery and benchmarking of mosaic SNVs across the whole genome. **A)** Distribution of COLO829BLT50 reference mutations by the number of GCCs in which each mutation was sampled across all datasets. VAFs were defined using the two highest depth datasets (470×). The number of reference mutations in each VAF group is denoted with n. **B)** Precision–sensitivity (recall) performance of each calling pipeline in Easy, Difficult, and Extreme genomic regions, aggregated into three VAF bins (<1%, 1–2%, and 2–3%). Points correspond to pipelines; point outline color denotes the pipeline, and point shape denotes the VAF bin. Shaded bands represent VAF-bin-specific ordinary least squares regression of precision on sensitivity, fitted separately within each region panel; shading indicates the 95% confidence interval for the fitted mean (no confidence band for individual predictions). **C)** Utilizing fibroblasts and iPSCs from the LA2 skin biopsy of LB donor allowed deriving best practices for discovery somatic mutations by bulk WGS at high (>5%) and low (<1%) VAFs using 1,360X coverage Illumina data. Mutations discovered in bulk fibroblasts (grey line) but not present in the iPSC reference mutation set had high confirmation rate by long read sequencing. **D)** Excess of mutation calls outside of easy regions (e.g., in GIAB strict only) in samples from post-mortem donors (as compared to genomic mask and fibroblasts) was mostly mitigated by applying fragment length and mapping quality filters. **E)** Ambiguous mapping in GIAB strict only regions (i.e., outside easy regions) was indicative of datasets with higher false positives (red for colon sample and blue for lung sample from donor ST002; black dots for samples from other donors). **F)** Applying best practices using data from production-like samples from ST002 donor demonstrated that combining coverage from tissues enables discovery of mutations that are shared by but present at low cell fractions in the tissues.

To assess the performance of various computational pipelines on somatic mutation detection, we conducted a consortium-wide low-VAF somatic SNV/indel detection challenge, involving 7 institutions and 12 somatic variant detection workflows using five replicates of COLO829BLT50 WGS data with depths ranging from 170×-470× and five computationally generated COLO829BLT50 WGS at 100× to 500×. An often-overlooked confounder in cell line–based benchmarking is the presence of mutations acquired in cell culture, via repeated expansion and harvesting of the cell line. We identified 5,568 of such cell-line mutations with high confidence (5,372 SNVs and 196 INDELS). By considering these true variants in the COLO829BLT50 mix, in addition to the variants present in the COLO829 tumor sample, we were able to more accurately evaluate the detection performance of the mutation calling methods. While the majority of the workflows performed well in detecting mutations above 2% VAF, all had a marked decline in performance for mutations at lower frequencies^15^. Furthermore, we observed that accuracy of mutation discovery strongly depended on the genomic regions (defined in **Box 1**) with significantly reduced sensitivity and increased false positive rate in difficult and extreme regions as compared to easy regions (**Fig. 3B**). We also found that beyond 300× sequencing depth additional coverage yields diminishing returns in detecting additional mutations at VAF 2% or higher^15^.

To evaluate solutions to the challenges associated with the discovery of low-VAF mutations, we leveraged the fibroblast benchmarking material using ultra-high coverage sequencing depth data, focusing on easy genomic regions^16^. Using the clonal iPSC lines derived from this fibroblast culture, we identified subclones with VAFs as low as 0.1% and 0.3%, enabling the establishment of low-VAF reference sets (**Fig. 3C**). Our analyses revealed that ∼1,360× Illumina WGS enables accurate detection of ∼60% of SNVs at VAFs 0.5-1%. In contrast, indels with VAFs <3% could be reliably detected only when located outside of homopolymeric regions.

##### Box 2.

**Key recommendations for mutation calling.**

**SNVs and indels:**

- Best practices should be depth-aware, method-specific, and region stratified.
- Optimization and reporting are to be done separately by regions; most confident discovery of mutations at low VAF is in regions with higher confidence of read mapping, e.g., easy regions.
- At coverage <300× MosaicForecast, machine learning based tool, provides the most balanced performance; higher coverage benefits probabilistic callers like Mutect2.
- Fraction of reads with zero mapping quality in GIAB strict mask outside easy regions highlight data that is prone to yield false positive calls (particularly outside easy regions).
- Filter calls by median read mapping quality (e.g., over 56), mean fragment length (e.g., over 300 bps) and mean position in read (i.e., over 20) to arrive to a high confidence call set.
- Use large panel of normal genomes to remove recurrent artifacts.
- Use assembly-based approaches (e.g., Lancet or Mutect2) for indel discovery, but performance is limited in complex sequence context.

**SVs:**

- **Read length matters more than depth alone; use long-read sequencing when possible**: PacBio HiFi / ONT substantially outperform short reads for insertions, repeats, and complex SVs.
- **Expect strong VAF limits**: sensitivity drops sharply below ∼1–4% VAF; even with deep coverage over 100X, low VAF SVs remain hard to recover reliably.
- **Stratify performance by genomic context**: repeats, insertions, small SVs, and clustered breakpoints are consistent failure modes—report performance by these strata, not just global F1. Currently no single tool provides the best performance across VAF range, sequencing modality, and SV difficulty stratification.
- **Tune callers for mosaic detection**: default SV caller settings are often optimized for germline or cancer; mosaic-aware modes (e.g., Sniffles mosaic) and lower support thresholds improve recall but require careful control of false positives.
- **Validate candidates with orthogonal evidence:** prioritize SVs supported by multiple reads/technologies (e.g., split-read consistency, breakpoint phasing) and manual review for low-VAF calls.

**MEIs:**

- For single-platform analyses with expected VAF >3%, ∼200× short-read WGS or ∼60× long-read WGS (with higher depth yielding additional performance) as a baseline offers an optimal cost-performance balance.
- For expected VAF <3%, long-read WGS is recommended for higher sensitivity and better performance in complex regions; targeted MEI assays (e.g., eHAT-seq, TEnCATS) provide cost-effective alternatives to WGS.
- For multi-platform analyses, performance improves by integrating evidence at the raw read/alignment level; precision can be further improved with 151-mer mappability filtering (short reads) and haplotype phasing (long reads).
- xTea_mosaic (short reads) and PALMER (long reads) show consistently strong performance; phasing- and/or DSA-based MEI detection is a promising upgrade for improved detection, especially in segmental duplications.

**Assembly:**

- Evaluation of somatic variation within the most sequence and structurally complex genomic loci largely necessitates use of a donor-specific reference assembly.
- Pangenome graph references enable the appropriate mapping of reads in low mappability regions and improve the precision and recall of somatic variant identification.
- Donor-specific and pangenome graph references improve somatic variation discovery in clinically relevant genes within complex loci.

**Single-molecule/duplex:**

- All techniques result in comparable results for SNVs
- ppmSeq stood out as the most cost-efficient technique making it amenable for deep genome-wide sequencing
- HiDEF-seq is unique in accessing single stranded DNA damage
- VISTA-seq and CompDuplex-seq can be applied to single cells
- CompDuplex-seq can be integrated with total RNA-based single-cell transcriptome assay for single-nucleus dual-omics
- Nano-seq, CODEC-seq, VISTA-seq, and CompDuplex-seq are amenable for detecting indels
- Nano-seq, CODEC-seq, and CompDuplex-seq are amenable to target-capture

**Single cell:**

- Extracting high quality nuclei is key for successful amplification
- Conduct 4-loci PCR to flag cells with uneven amplification
- Once data is generated, conduct different QC for SNV/indel and for CNV analyses
- To increase precision of the call set use specialized callers that take into account allelic imbalance across genome
- Coverage over 15X per cell improves sensitivity of mutation discovery but does not improve accuracy
- Use highest stringency calls to estimate mutation burden and spectra in each cell
- Use more permissive calling for reconstructing developmental cell phylogenies
- High confidence CNVs and SVs are supported by coverage-based (i.e., read depth or b-allele frequency) and fragment-based (split read or discordant read pairs) evidence

To assess the effects of various sampling approaches and sample conditions in real tissues on genome-wide somatic SNV discovery, we leveraged the post-mortem donor tissue benchmarking material to detect somatic mutations^16^. As compared to live fibroblasts, in post-mortem tissues we observed decreased accuracy of mutation discovery that was particularly prominent in non-easy genomic regions (e.g., in GIAB strict mask outside easy regions), where confidence of mapping short reads is reduced. This likely reflects overall lower quality of and more damage to DNA due to post-mortem interval before sample collection^16^. We discovered that filtering calls by read mapping quality and fragment length supporting mutations largely eliminates excessive false positives with little reduction in the true calls (**Fig. 3D**), while read mapping in specific genomic regions highlighted data that is prone to higher rate of false positive calls (**Fig. 3E**). Finally, we demonstrated that combining 200-300× short-read WGS datasets from multiple tissues from the same donor to reach higher coverage is a viable strategy to enable the discovery of low VAF (<3%) mutations shared across tissues, which would otherwise be undetectable when each tissue is analyzed individually (**Fig. 3F**). Key recommendations for calling somatic SNVs are summarized in **Box 2**.

#### Benchmarking alternative assembly approaches for somatic mutation discovery

Current human somatic mutation catalogs are largely generated by mapping sequencing reads to either the GRCh37 or GRCh38 reference genomes. However, the actual diploid 6 Gbp genome of an individual can differ substantially from these haploid reference genomes, causing inaccurate read alignments and large missing genomic segments^19^, both of which can contribute to false positive and false negative somatic mutation calls (**Fig. 4A**). We sought to evaluate whether human pangenome references^33,34^, or diploid donor-specific assemblies (DSA)^35,36^ would impact the performance of somatic variant detection. We first developed a somatic benchmarking mutation call set for the HapMap mixture using a graph-based framework built from HapMap individual assemblies^17^. We then assessed the precision and recall of somatic mutation discovery using the HapMap mixture short-read sequencing data mapped to either GRCh38, a reference pangenome, or a pangenome-inferred diploid assembly (**Fig. 4B**). This revealed that pangenome-guided alignments improve mutation calling accuracy compared with GRCh38, especially in extremely challenging genomic regions. In 500× short-read data, precision increased from 0.79 to 0.89 and recall from 0.48 to 0.59 for SNVs, while precision increased from 0.46 to 0.52 and recall from 0.09 to 0.13 for indels (**Fig. 4C**)^18^. For easy loci, accurate detection of ∼70% of SNVs was possible for VAFs >0.5-1%, but for extreme loci, this was limited to VAF >3-4%. In 100× long-read data, precision increased from 0.53 to 0.55 and recall from 0.16 to 0.28 for SVs. While improving mutation detection at the expense of significantly more intensive computations (**Table S7**), the pangenome approach still does not benchmark somatic mutation calling in some of the most challenging genomic regions, such as satellite arrays, which contain large stretches of DNA that differ between individual haplotypes and are consequently often clipped from pangenome graphs owing to their complexity.

**Figure 4.**
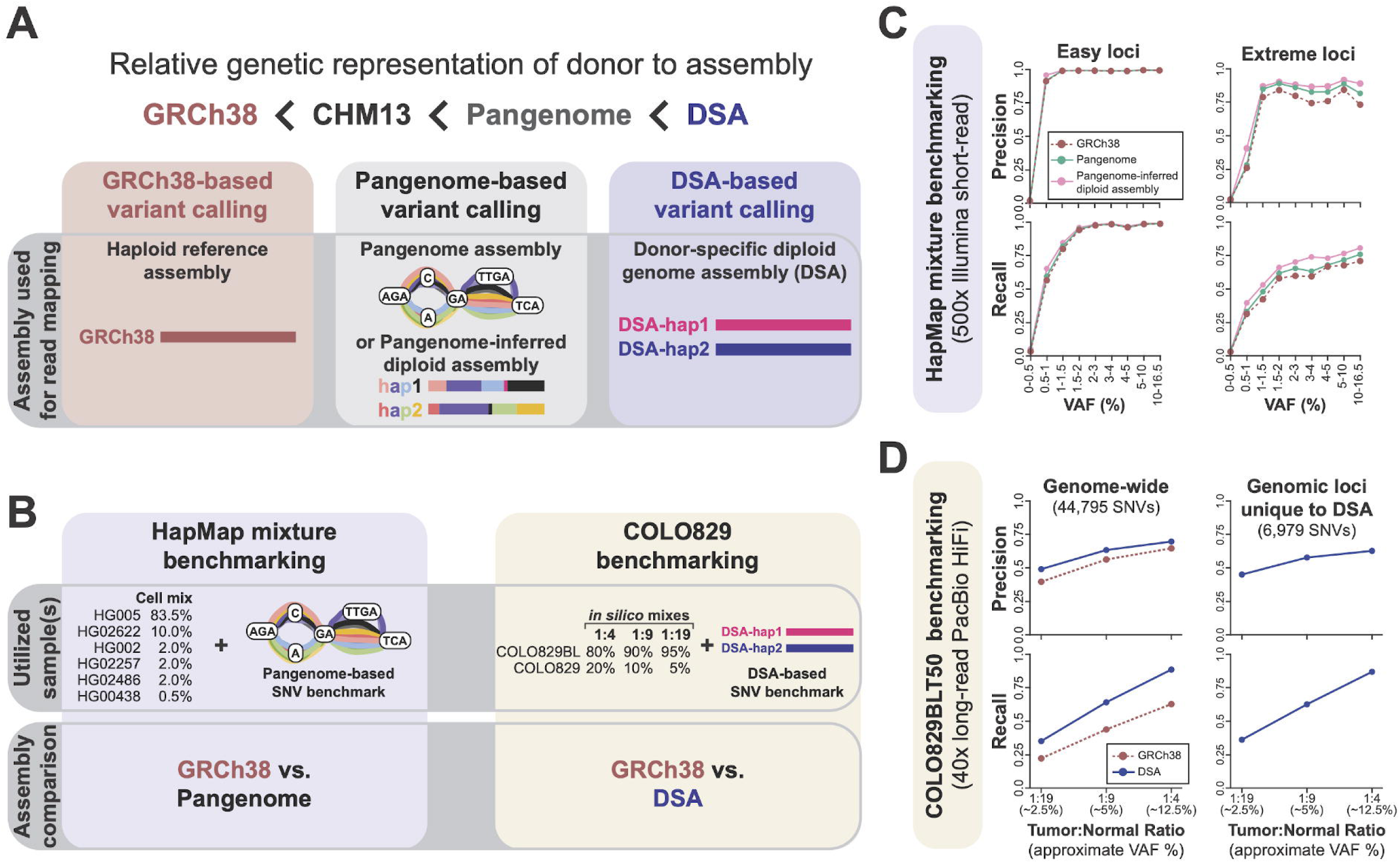
Benchmarking alternative assembly approaches for somatic mutation discovery. **A)** Conceptual diagram showing the similarity of different genome references to the genome of the studied individual. By definition, DSA is the most similar, as it is generated directly from the genome of the individual being studied. A pangenome is a reference that includes a population of individuals, but not the individual being studied, and thus is less similar. **B)** A framework for evaluating the utility of a pangenome- and DSA-based approaches for improving somatic variant detection. **C)** Precision and recall for somatic variant discovery using 500x Illumina short-read sequencing of the HapMap mixture using GRCh38- and pangenome-based approaches. Findings are separated based on the genomic mappability of the locus in which that somatic variant exists. **D)** Precision and recall for somatic variant discovery using 40x PacBio HiFi long-read sequencing data of *in silico* mixture of the COLO829BL and COLO829 lines for GRCh38- and DSA-based approaches. Findings are separated into genome-wide, as well as genomic loci that are unique to the COLO829BL DSA.

To more comprehensively benchmark somatic mutation discovery genome-wide, including within regions that lack synteny with GRCh38, we leveraged the COLO829 benchmark, for which we constructed a paired DSA for COLO829BL^19^. We then assessed the performance of somatic mutation discovery using *in silico* mixtures of the COLO829 and COLO829BL PacBio HiFi long-read sequencing data mapped to either GRCh38, CHM13, or the COLO829BL DSA (**Fig. 4B**). This revealed that using the DSA as the reference improved both the precision and recall of somatic mutation calling accuracy compared with either GRCh38 or CHM13 and enabled the accurate discovery of somatic mutations within genomic regions absent from reference genomes (**Fig. 4D**). This includes mutations within the kinetochore-binding region of the centromere, for which the paired single-molecule CpG methylation data can be used to assess associated functional alterations in centromere structure^19^. Notably, we found that CHM13 performed worse than GRCh38 for mutation detection, which is driven by unique and complex regions in CHM13 that are hypervariable between individuals. These regions are not yet well accounted for in current catalogues of germline variants resulting in an excess of false positive somatic calls (**Fig. S1**). Together, these findings demonstrate the utility of pangenome- and DSA-based approaches for improving somatic mutation discovery, especially within traditionally challenging genomic regions that are known to exhibit some of the highest rates of somatic variation^19^. Furthermore, these findings highlight the importance of using an appropriate panel of normal germline variation when performing somatic mutation discovery on a reference genome other than the donor‘s. We anticipate that development of additional computational tools that use diploid genome assemblies and graph genomes will further improve the ability of these emerging reference approaches for accurately identifying the full catalog of somatic mutations within an individual. Key takeaways for assembly-based mutation calling are summarized in Box 2.

#### Benchmarking experimental and computational methods for somatic SV and MEI discovery

We next sought to benchmark the performance of somatic SV detection using the HapMap benchmark data^20^. First, we developed a high-confidence somatic SV reference set by harmonizing SVs across the six HapMap diploid assemblies (**Figs. 1A** and **5A**), resulting in ∼34k insertions and deletions, which we validated using ddPCR and other orthogonal methods. Using this reference set, we benchmarked 12 somatic SV detection methods across Illumina, PacBio HiFi, and ONT sequencing data. Overall recall and precision of each strategy varied substantially across platforms, prompting a difficulty-stratified benchmark to pinpoint the causes of underperformance (**Fig. 5B**). This difficulty-stratified analysis revealed limitations with short-read sequencing for detecting insertions, small SVs, and tandem repeats, and limitations with both short- and long-read sequencing for detecting low-VAF SVs and adjacent SVs. Under matched sequencing depths (60-180×), we observed distinct platform-specific ceilings: short-read sequencing (i.e., Illumina) plateaued at ∼30% recall for SVs with a 5-20% VAF, even at 180× coverage, whereas long-read sequencing (both PacBio HiFi and ONT) achieved ∼40% recall for 1-5% VAF and ∼80% for 5-20% VAF at only 60× coverage (**Fig. 5C**).

**Figure 5.**
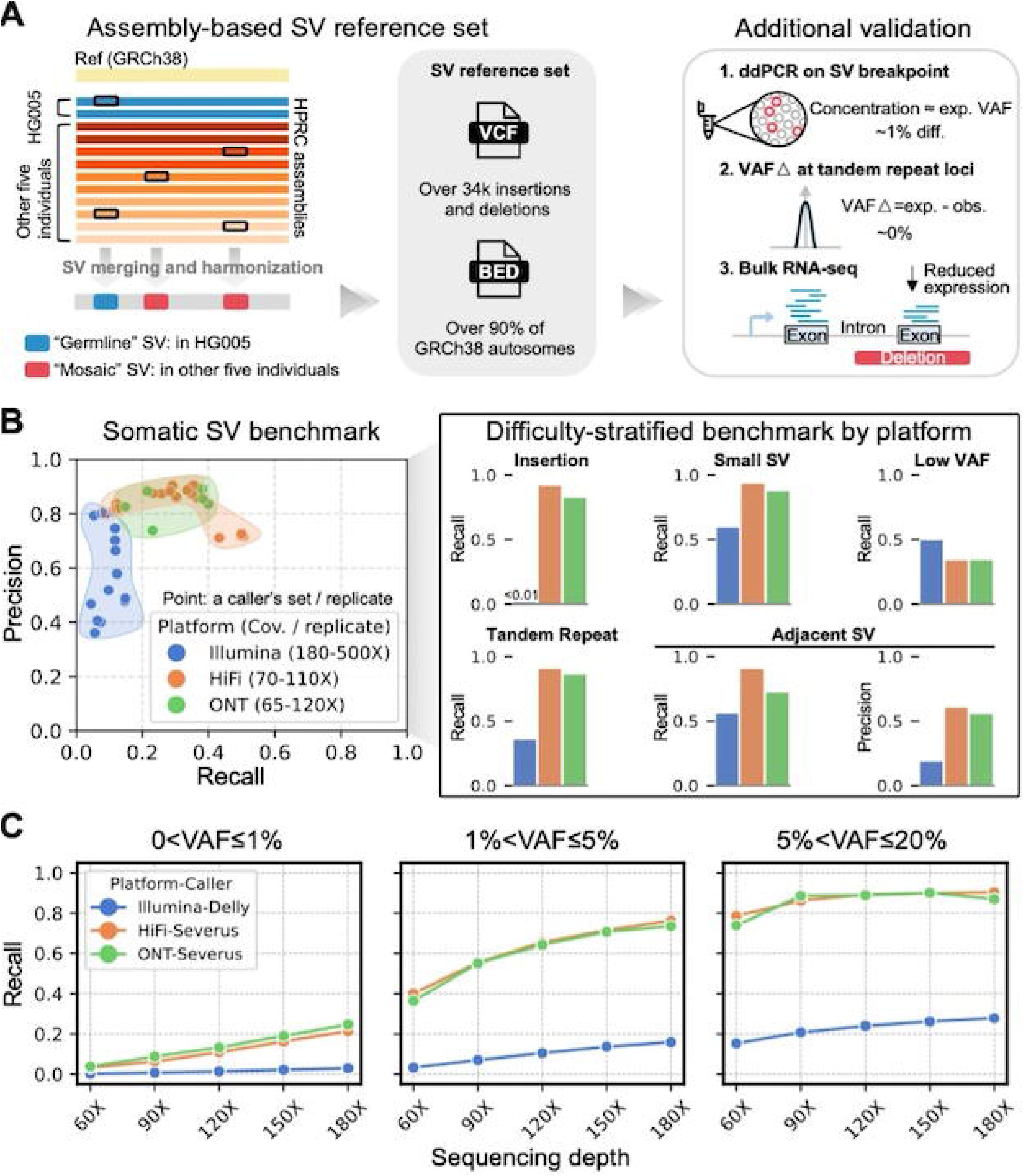
Assembly-based SV reference set for the HapMap mixture benchmark and cross-platform benchmarking of somatic SV calling. **A)** The SV reference set was constructed by harmonizing diploid assembly-based SV calls from six individuals, validated via ddPCR, VAF checks at tandem repeat loci, and bulk RNA-seq. **B)** Cross-platform performance of somatic SV callers on each replicate, plus a difficulty-stratified benchmark covering insertions, small SVs (50-250 bp), low-VAF (<10%), tandem repeat-associated SVs, and adjacent SVs (breakpoints ≤ 500 bp apart). Illumina, HiFi, and ONT are shown in blue, orange, and green (legend on the left). **C)** Platform-specific detection limits at matched sequencing coverages, stratified by VAF bins: ≤1%, 1-5%, 5-20%.

MEIs represent a clinically relevant subclass of SVs that are known to cause somatic variation^37–39^. As such, we sought to benchmark the performance of somatic MEI detection using a combination of computational methods and data from ‘core’ and ‘extended’ experimental assays^21^. First, using the HapMap mixture, we established a somatic benchmarking MEI reference set annotated with target-primed reverse transcription (TPRT) features critical for distinguishing RNA-mediated retrotransposition from DNA-level genomic rearrangements (**Table 1**, **Fig. 6A**). Our evaluation across sequencing platforms, depths, VAFs, and genomic regions highlighted the advantages of long-read WGS and ME-targeted sequencing for detecting MEIs at low VAFs (< 3%) and in challenging genomic contexts (**Fig. 6B**). Short-read WGS performance saturated at ∼200X, whereas 60X long-read WGS achieved comparable performance. Notably, long-read methods demonstrated 1.6-fold higher recall than short-read methods for MEIs at VAF <1%, while also resolving complete insertion structures and providing CpG methylation profiles, a key indicator of epigenetic ME suppression. ME-targeted assays achieved 72% recall for MEIs even with VAF <1%, offering cost-effective alternative to WGS-based approaches for detecting very low VAF events. Given the distinct error profiles between short- and long-read call sets, cross-platform integration combined with rescuing raw read-level signals further improved detection, yielding a 1.6-fold higher F1 score of 0.75, compared to simple intersection from 200× Illumina and 60× PacBio data, sequencing depths achievable for most SMaHT production samples (**Fig. 6B**).

**Figure 6.**
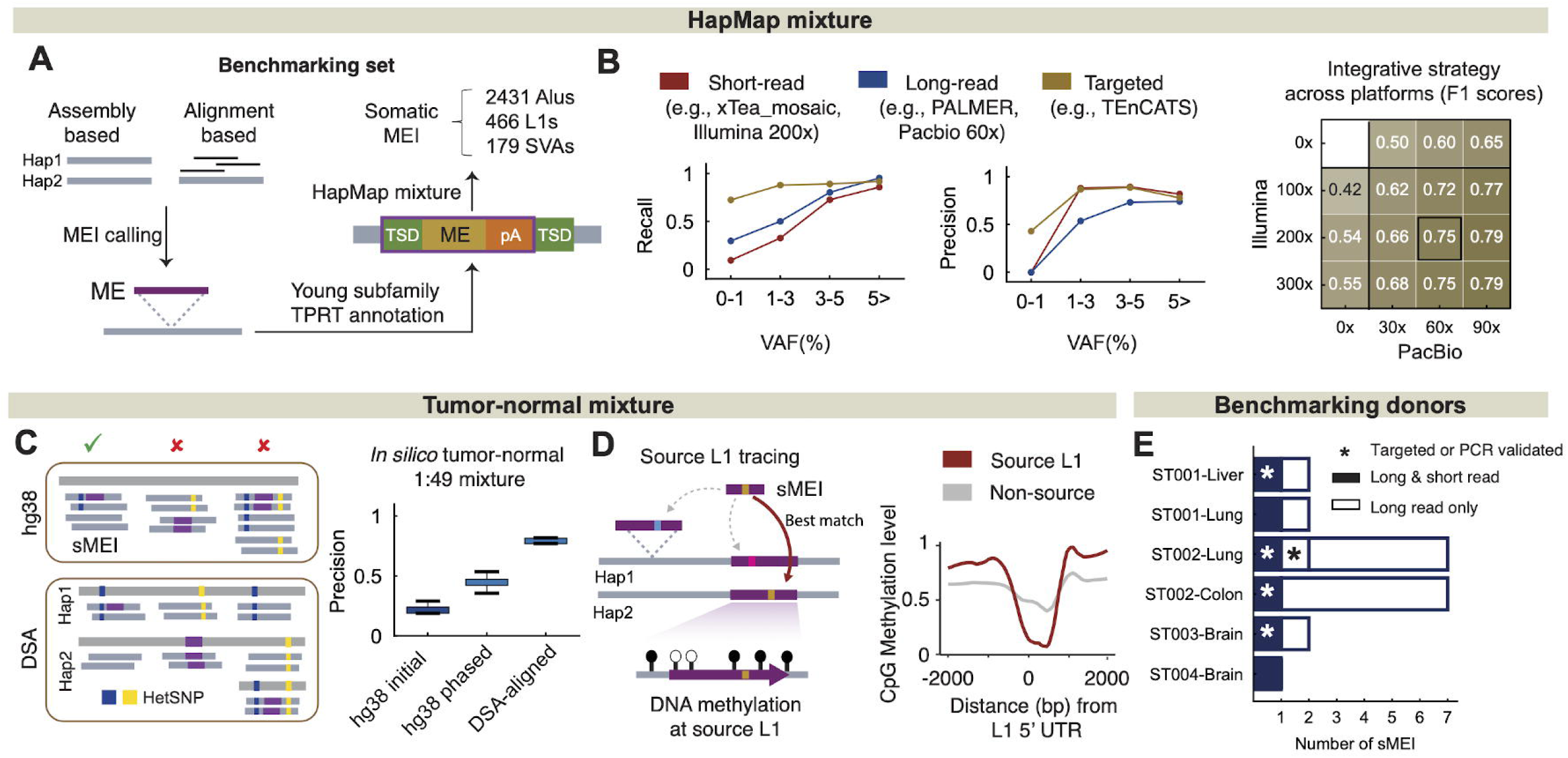
Benchmarking and Optimized Discovery of Somatic MEIs. **A)** Establishment of benchmarking set in HapMap Mixture. An sMEI truth set was generated as the foundation for evaluating performance, leveraging multiple technologies and incorporating information from repeat element subfamilies as well as target-primed reverse transcription (TPRT) features. **B)** Benchmarking sMEI detection in the HapMap Mixture. Performance (recall and precision) was assessed across multiple sequencing platforms, including Illumina, PacBio, ONT, and transposon-targeted sequencing, as well as across different ranges of variant allele frequencies (VAFs). The improvement of a multi-platform integration strategy on detection performance was evaluated based on different sequencing depths. **C)** Enhanced sMEI detection through haplotype phasing and DSA in an In-silico Tumor-Normal Mixture using the CASTLE project data. Precision improvements are demonstrated using two strategies at 2% mixture or 1% VAF level. **D)** Long-read data facilitates precise mapping of source L1 elements by leveraging internal sequence variation and assessment of DNA methylation profiling of the 5′ UTR region. **E)** Barplot of the number of somatic L1 insertion candidates identified in the benchmarking donors after implementing the optimized strategies from A, B, and C.

To further assess the performance of mutation detection using haplotype phasing and direct read alignment to DSA, we generated an *in-silico* tumor-normal mixture from the CASTLE project^40^, where 376 somatic L1 insertions were identified in the tumor sample. Even at a VAF of 1% (i.e., 2% mixture), precision improved 2.1-fold with haplotype phasing and 3.7-fold with the DSA method compared to long-read-based detection alone (**Fig. 6C**). We also developed a novel source-tracing pipeline leveraging L1 internal sequence variation, which successfully identified haplotype-resolved source elements for three times more somatic L1 insertions (67% vs 22% of insertions > 1Kbp) compared to conventional transduction-based tracing. These source L1s showed hypo-CpG methylation at promoter (5’ UTR) regions, consistent with these representing active L1 elements (**Fig. 6D**). Finally, applying these optimized strategies to benchmark donor tissues, we identified 21 somatic L1 insertions (all VAFs < 1%) across all tissues profiled—liver, lung, colon, and brain. All events were fully resolved at the sequence and structural level, including the first discovery of somatic U5 snRNA-L1 chimeric retrotransposition in normal human tissues (**Fig. 6E**). Key recommendations for calling somatic SVs and MEIs are summarized in **Box 2**.

#### Benchmarking duplex and single-cell sequencing methods

The aforementioned benchmarks outline the challenges associated with identifying somatic mutations present in <1% of cells using bulk short- or long-read sequencing methods, indicating that bulk sequencing approaches are best powered to find mutations that occur early in development or that are present in expanded clonal lineages (**Fig. 7A**). For instance, the fibroblast and the tissue donor benchmarking experiments showed that even at very high sequencing depths (∼1,400x) only mutations above 0.5% VAF can be detected with sufficient sensitivity; such VAF would normally correspond in a human to the pre-gastrulation embryonic period. In contrast, duplex and single-cell sequencing approaches have the potential to probe the somatic mutation landscape across the entire lifespan irrespective of clonality (**Fig. 7B**). Duplex sequencing tracks both strands of each original DNA molecules and calls a mutation only when complementary changes are seen on both strands, suppressing sequencing and amplification errors. This technology detects somatic mutations with high cost-efficacy and provides unbiased single molecule resolution, enabling simultaneous profiling of clonal and rare somatic SNVs and indels and their average burdens and spectra. In contrast, single cell sequencing involves whole genome amplification and sequencing of individual cells from a sample, enabling the assessment of the compendium of somatic mutations present within single cells, which can be used to uncover a cell’s mutational burden and spectra and ancestral relationships between cells.

**Figure 7.**
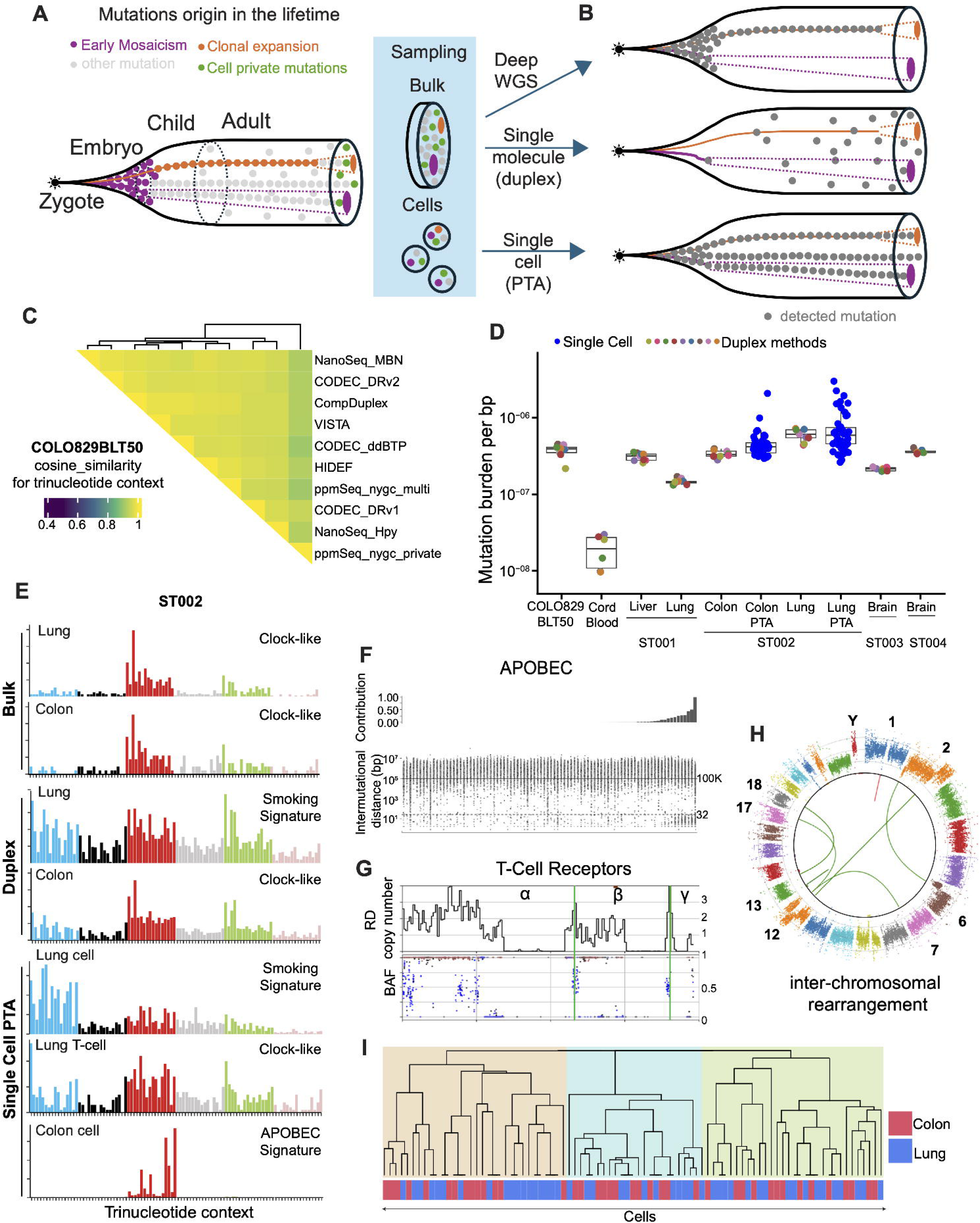
Benchmarking mutation discoveries using duplex and single cell analyses. **A)** Mutations in human cells occur across the life span. Early developmental and clonally expanded mutations will be frequent in sampled bulk tissues (orange and purple circles). They can also be sampled in single cells along with cell private and other mutations occurred during lifetime. **B)** Bulk WGS is limited to assessing mutations occurring early in life or from clonal expansions. Duplex sequencing approaches discover mutations independent of their clonal prevalence. Single cell analyses discover mutations in the analyzed cell regardless of their occurrence in time and clonal prevalence. **C)** Different duplex approaches are concordant in mutation spectra in COLO829BLT50 sample. **D)** All duplex sequencing approaches yield consistent mutation burdens estimates revealing heterogeneity between tissues^22^. PTA analyses results in concordant estimates of average mutation burdens while revealing heterogeneity between cells. **E)** Mutation spectra of two tissues profiled by deep bulk WGS, duplex sequencing, and PTA analyses. Bulk profiling is dominated by early developmental mutations and spectra in two tissues are therefore indistinguishable. Duplex-sequencing, given its unbiased profiling capability, captures tissue specific mutational patterns. PTA further resolves mutation heterogeneity at single cell level including developmental signature, APOBEC signatures, and exposure to tobacco smoking. **F)** High contribution of APOBEC mutagenesis was pronounced in a number of cells. Such cells also exhibited increase in closely spaced (within 32 bps) mutations previously termed didyma^41^. **G)** Five T-cells were identified from discovered rearrangement and deletions at the known T-cell receptor loci (heterozygous SNPs are in blue; homozygous SNPs are in brown). **H)** Some cells had large CNVs that were related to inter-chromosomal rearrangements (rearrangements in one cell are shown). **I)** Sharing of mutations between cells allowed reconstructing cell ancestry lineages. The root of the tree likely represents a zygote, while three highlighted developmental clades are apparent^23,25^.

Aligned with the SMaHT objectives of discovering mutations across the entire genome, we conducted a comprehensive benchmarking of six different whole genome duplex sequencing technologies (and their variants): CODEC, CompDuplex, HiDEF-seq, NanoSeq, ppmSeq, and VISTA-seq – applying them to the engineered COLO829BLT50, six tissues from post-mortem donors (**Fig. 1D**), and umbilical cord blood^22^. While the technologies differed in their library construction, sequencing platforms, depth of sequencing, cost-efficiency, genomic footprint, sensitivity, and ability to detect indels, the comparison revealed that the inference of mutation burden and signatures of both SNVs and indels was highly concordant across all methods (**Fig. 7C,D**). Specifically, in the donor ST002, we observed a higher mutation burden in lung than in colon, which likely can be attributed to smoking of the donor and that is reflected in the substitution mutation spectrum and smoking associated signature (**Fig. 7E**; donor information is available at dbGAP collection). Moreover, duplex sequencing yields a largely distinct set and profile of mutations from deep bulk WGS. Mutational signature analysis demonstrated that duplex sequencing uncovers mutational processes at low variant allele fractions (0.01–1%) that are largely undetectable by deep bulk WGS, consistent with their later emergence during development and/or lifetime. Overall, duplex sequencing reveals a much more extensive and unique repertoire of <1% VAF clonal mutations, although deep duplex sequencing will be required to estimate their VAFs directly from data. Currently, the choice of duplex technology will depend upon specific desired applications including targeting specific regions or conducting genome-wide assessment, ability to detect indels or single-strand damages, input material requirements, and cost (**Box 2**).

For donor ST002, we also performed primary template-directed amplification (PTA) followed by short read whole-genome sequencing (∼30× coverage) of 102 single nuclei from the lung and colon^23^. The analysis of the PTA data reproduced the differences in mutation burden between lung and colon found by duplex sequencing (**Fig. 7D**) and revealed that a fraction of cells from the lung are strongly impacted by signatures of tobacco exposure, while several cells from lung and colon revealed the signature of APOBEC mutagenesis. Such cells also exhibited an increase in closely spaced (within 32 bps) mutations previously termed didyma^41^ (**Fig. 7F**). The observed mutation heterogeneity between cells can be partially explained by the mixture of different cell types in a tissue, as, for example, in both lung and colon samples we found the presence of T cells (five in total) as evidenced by genomic rearrangement at T-cell receptor loci (**Fig. 7G**). Moreover, several cells harbored large CNVs, chromosomal aneuploidies, inter-chromosomal rearrangements (resembling that found in cancers), and, consistent with expectations for an aged male, frequent losses of chromosome Y (F**ig. 7H**). Finally, despite regional dropouts and amplification noise, PTA allowed detection of mutations shared between single cells, enabling for the first time the reconstruction from amplified cells of a phylogenetic tree of cellular lineages rooted at the zygote and validated by bulk sequencing (**Fig. 7I**). Together, duplex and single cell analyses revealed a rich landscape of somatic mutations that was otherwise obscured by bulk sequencing, while PTA positioned itself as a universal approach for single cell genomics with unbiased recovery of mutations across cells, tissues, and developmental timeline. Recommendations for conducting single cell analyses with PTA are provided in **Box 2**.

### Discussion

We present a comprehensive roadmap for the detection of somatic mosaic mutations. This effort assessed the detection of various types of somatic mutations—including SNVs, indels, MEIs, and SVs/CNVs—across a broad range of VAFs, utilizing multiple technologies such as bulk deep whole-genome, single-cell, duplex, and targeted sequencing. The benchmark experiments and the analyses highlighted here from multiple companion publications revealed several key takeaways: (i) while bulk sequencing at 200-300x has limited sensitivity to detect mutations with VAF <2%, combining coverage across several tissues to reach a sequencing depth of ∼1,400×, allows accurate discovery of SNVs at VAFs of 0.5-1%, making possible the discovery of low frequency mutations shared across tissues (**Fig. 3F**); (ii) the use of personal genome assemblies and pangenome assemblies enabled more accurate (both in terms of sensitivity and specificity) detection of somatic mutations, especially in repetitive genomic regions, than standard reference linear genomes (**Fig. 4**); (iii) read length matters more than depth alone for identifying somatic mutations (including SNVs, SVs, and MEIs) in repetitive regions (**Figs. 3D,5,6**); (iv) various whole genome duplex techniques, while having advantages and disadvantages, uncover concordant tissue-specific mutation burden and spectra that are obscured by bulk sequencing; (v) PTA is a comprehensive single-cell genomics approach that yields an expansive view of diverse somatic mutation types from development through aging across diverse tissues (**Fig. 7**). Our analyses and comparisons across technologies, cells and tissues also allowed us to make practical recommendations for detecting somatic mutations of different types (**Box 2**).

We observed that bulk-, duplex-, and PTA-based single-cell WGS provide complementary and orthogonal means of validating findings from one another, where each approach offers distinct variant discovery niches with unique error profiles that could be corrected. We note that, although duplex sequencing and PTA-based single-cell analyses offer unique insights into tissue mosaicism, they are not yet complete substitutes for the bulk WGS approach, which delivers uniform genome coverage with <1% VAF detection sensitivity at sufficient coverage. Duplex sequencing is currently best suited for genome-wide sampling of low-VAF SNVs and indels, enabling assessment of average mutational burdens and profiles. As the duplex methods continue to advance, they can be predicted to have better utility for routine detection of SVs or MEIs. PTA, on the other hand, enables high-resolution analysis of SNVs, indels, and CNVs at the single-cell level, but is a medium throughput assay and comes with significant cost when scaling to hundreds of cells per sample. This drawback can also be anticipated to decrease with the advance of robotic technologies^42^.

The choice of an experimental approach depends on the specific scientific or clinical question being addressed, as well as budgetary considerations. Based on our analyses, we identified several common scenarios that researchers and clinicians are likely to encounter and provide recommendations for the most suitable strategies (**Table 2**). A common case is the detection of somatic mutations of interest—whether pathogenic or functionally relevant—including SNVs, SVs, or MEIs with VAFs greater than 1%. This scenario is best addressed using bulk whole-genome sequencing (WGS), with long-read sequencing particularly advantageous for detecting SVs and MEIs. Duplex sequencing would require substantially higher coverage to achieve comparable sensitivity and is not applicable to SVs and only limited in its applicability to MEIs. Similarly, detecting mutations at these VAFs using single-cell sequencing would require amplification and analysis of dozens of cells, making it cost-inefficient. For these reasons, duplex and single-cell approaches are generally less favorable for this application.

**Table 2.** Recommendation of experimental approaches and analyses to address specific research and clinical challenges.

| **Challenge/Analysis approach** |  | **Bulk** | **Duplex** | **scWGA** |
| --- | --- | --- | --- | --- |
| **Detecting mutations of interest at >1% VAF (e.g., pathogenic or cancer driver)** | **SNVs and indels** | Recommended WGS or targeted sequencing | Recommended targeted sequencing. Not efficient for shallow whole-genome duplex, and success can depend upon choice of bulk control | Not cost-efficient |
|  | **SVs and complex rearrangements** | Recommended with long reads | Unresolved | Not cost-efficient |
|  | **MEIs** | Recommended with long reads or multi-platform integration | Unresolved | Recommended targeted sequencing |
| **Detecting mutations of interest at <1% VAF (e.g., pathogenic or cancer driver)** | **SNVs and indels** | Recommended WGS or targeted sequencing | Recommended if targeted sequencing. Feasible with shallow duplex, but VAF estimation not possible. | Not cost-efficient |
|  | **SVs and complex rearrangements** | Limited sensitivity and precision | Unresolved | More reliable option for large CNVs and SVs; not cost-efficient |
|  | **MEIs** | Recommended if targeted | Unresolved | Recommended targeted sequencing |
| **Detecting and monitoring mutational (SNV and indel) exposures: burden and spectra** |  | Limited to developmental early and clonally expanded mutations | Recommended | Recommended but more costly than duplex |
| **Detecting and monitoring clonal expansion by SNVs and Indels** |  | Recommended but not definitive | Limited to deep targeted sequencing | Definitive but not cost-efficient |
| **Analyzing mutations in repetitive genomic loci (satellites, VNTR, SegDups)** |  | Recommended with long reads and DSA/Pangenome | Unresolved | Unresolved |
| **Lineage tracing** |  | Not conclusive | Not conclusive | Recommended |
| **Functional consequence of mutation(s) of interest** |  | Applicable for homogeneous cell types with expression or epigenome readout | Unresolved | Applicable when conducting with expression or epigenome readout from the same cell |

For detecting mutations at lower VAFs, targeted sequencing is typically the most cost-efficient solution, while single-cell sequencing remains the only viable approach for detecting low-VAF SVs. When the goal is not to identify specific mutations but rather to characterize the overall mutational landscape of a sample, duplex sequencing is the most favorable approach. Single-cell sequencing can provide an even more detailed view of mutational heterogeneity, but at a substantially higher cost.

Detecting and monitoring clonal expansions can be achieved using both bulk and single-cell sequencing. Single-cell sequencing provides a definitive solution, as it unambiguously resolves whether multiple mutations co-occur within the same clone or across distinct clones. Bulk sequencing, while less definitive in this regard—since mutations from different clones may appear at similar VAFs—is more cost-efficient because it does not require sequencing large numbers of individual cells.

Based on our experience, the analysis of mutations in highly repetitive genomic regions (such as VNTRs, satellite repeats, and segmental duplications) is feasible with long-read sequencing, particularly when diploid synthetic assemblies (DSAs) are available. While the application of duplex and single-cell sequencing to these regions is theoretically possible, it remains an open technical challenge. For reconstructing cellular ancestry and building phylogenetic trees, single-cell sequencing is currently the only available approach.

Finally, elucidating the functional consequences of somatic mutations requires a functional readout, such as RNA sequencing, CpG methylation profiling, or an assessment of chromatin configuration or histone biochemical marks. Several such analyses have been benchmarked as part of SMaHT companion papers. For example, it was demonstrated that single-molecule chromatin fiber sequencing (Fiber-seq) can be used to accurately identify the functional impact of genetic mutations with single-molecule precision, and can also quantify somatic epimutations genome-wide, including in primary human tissue^43^. In addition, application of Fiber-seq to the COLO829 benchmarking material uncovered somatic epimutations altering the regulation of critical oncogenes^19^. Furthermore, it was demonstrated that Deaminase-Assisted Fiber-seq (DAF-seq) is an accurate and sensitive approach for quantifying the functional impact of low-VAF somatic mutations, as well as resolving the complete genomic and chromatin epigenomic map of single-cells^26^, providing a path for future studies into the functional consequence of somatic variation. Concurrent profiling of genome sequence and functional output from the same cell provide a promising path forward, and approaches extending PTA to enable joint genome, transcriptome, and functional assays have recently emerged and show considerable promise^25,26^. For instance, combined genome-transcriptome amplification from the same cell permits the identification of cell types carrying specific mutations and assess potential functional consequences on the transcriptome. Duplex sequencing coupled with functional readouts could, in principle, serve a similar role^27^.

The conclusions and described datasets represent a valuable community resource for benchmarking the current and emerging sequencing technologies, studying somatic mosaicism, and advancing the development of analytical methods for somatic mutation discovery and analysis. Ultimately, the benchmark results described in this study have helped define the experimental and analytical approaches to apply for the production phase of the SMaHT project to enable the comprehensive detection of somatic mosaicism across tissues. Specifically, the SMaHT Network will adopt a tiered approach where selected donors with high quality tissue samples derived from the three germ layers will be profiled more extensively in addition to the core assays applied for the other donors. To study mosaicism more extensively in the select donors, we will integrate bulk, duplex, and PTA approaches, produce higher coverage short- and long-read WGS, as well as construct and utilize DSA to achieve the most comprehensive and accurate characterization of somatic mosaicism across the human body.

#### Limitations of the Study

The findings from our study also illumined two key considerations when designing benchmarking experiments and choosing benchmarking materials used to detect somatic mutations. First, extensive culturing of cell lines may lead to culture acquired mutations at high VAF, complicating the definition of a comprehensive true-positive mutation set in the test sample consisting of a mixture of cell lines. Second, specificity estimates derived from experiments with *in vitro* samples, although useful for method development, do not necessarily extrapolate to post-mortem tissues. Hence, when analyzing such samples, it is essential to provide orthogonal evidence to support the validity of mutation calls even when utilizing the approach validated on *in vitro* samples. To address the second challenge at least in part, we used multiple sequencing methodologies and compared the data across different GCCs. Furthermore, we identified several challenges in mutation discovery that haven’t completely been solved, which are often related to repetitive regions of the human genome and to analyses of genomic rearrangements, which are also enriched in repetitive regions (**Table S8**).

Our benchmarking efforts further enables us to propose a more streamlined and rational design based on creating controlled mixtures of multiple iPSC lines. It is feasible to derive several iPSC lines from a single donor, using different sample sources (e.g., fibroblasts, blood, and urine-derived cells), with each line carrying distinct somatic mutations. Mixing these lines would allow precise control of mutation frequencies while preserving the original diploid haplotype structure, closely mirroring real biological samples. “Negative control” mutations—those absent from the mixture—could be defined using iPSC lines not in the mix, as demonstrated in our fibroblast-based benchmark. iPSC lines are more genomically stable than cancer cell lines, making them less prone to culture-induced artifacts and providing an inexhaustible source of material for validation and follow-up studies. These lines, along with primary fibroblasts, could also serve as the foundation for generating telomere-to-telomere DSAs, enabling the development and benchmarking of methods that leverage personalized reference genomes. One limitation of this approach is the relatively low abundance of SVs, CNVs, and MEIs in iPSC lines, compared to SNVs, reflecting relatively low frequencies of these variant types in normal human tissues. However, this could potentially be mitigated by inducing structural variation and MEI activation in primary cells prior to reprogramming.

## Resource availability

This study was conducted as part of the NIH Common Fund consortium initiative, Somatic Mosaicism across Human Tissues (SMaHT). Samples, data, and relevant software availability is described in the KEY RESOURCES TABLE. More information about the SMaHT Network and data is available online at https://smaht.org and at https://data.smaht.org.

## Acknowledgements

We are extremely grateful to the SMaHT donors, and donor families, who have generously provided such precious gifts to support this important work. This research is supported by the NIH Common Fund, through the Office of Strategic Coordination/Office of the NIH Director under awards U24 MH133204, U24 NS132103, UG3 NS132024, UG3 NS132061, UG3 NS132084, UG3 NS132105, UG3 NS132127, UG3 NS132128, UG3 NS132132, UG3 NS132134, UG3 NS132135, UG3 NS132136, UG3 NS132138, UG3 NS132139, UG3 NS132144, UG3 NS132146, UM1 DA058219, UM1 DA058220, UM1 DA058229, UM1 DA058230, UM1 DA058235, and UM1 DA058236.

## Author contributions

### Writing group

Alexej Abyzov, Elizabeth Chun, Tim Coorens, Harsha Doddapaneni, Sheng Chih Jin, M. Kathryn Leonard, Julia Markowski, Alexi Runnels, Fritz Sedlazeck, Andrew B. Stergachis, Flora M. Vaccarino, Shadi Zaheri, Weichen Zhou.

**Figure 1** is contributed by Julia Markowski and Wenxuan Hong.

**Figure 2** is contributed by Julia Markowski and Elizabeth Chun.

**Figure 3** is contributed by Yoo-Jin Jiny Ha and Yeongjun Jang.

**Figure 4** is contributed by Qichen Fu, Sheng Chih Jin, Nahyun Kong, Youngjun Kwon, Juan F. Macias-Velasco, Benpeng Miao, Anna Minkina, Min-Hwan Sohn, Andrew B. Stergachis, Zitian Tang, Zilan Xin.

**Figure 5** is contributed by Yuwei Zhang and Fritz Sedlazeck.

**Figure 6** is contributed by Seunghyun Wang, Mingyun Bae, Jinhao Wang, Boxun Zhao, Ryan Mills, Weichen Zhou, Eunjung Alice Lee.

**Figure 7** is contributed by Tim Coorens, Joe Luquette, Abhiram Natu, Milovan Suvakov, and Yang Zhang.

### Method and supplement writing contributors

Thomas J Bell, Michele Berselli, Thomas Blanchard, Hsu Chao, Rui Chen, Elizabeth Chun, Harsha Doddapaneni, Gilad Evrony, Christian D. Frazar, Robert Fulton, Stephanie Gardiner, Stephanie Georges, Chris Grochowski, Yoo-Jin Jiny Ha, Eric Ho, Shen Hui, Caitlin N. Jacques, Yeongjun Jang, Sheng Chih Jin, Ben Johnson, Ziad Khan, Youngjun Kwon, Eunjung Alice Lee, M. Kathryn Leonard, Mary Majewski, Heer Mehta, Benpeng Miao, Donna Muzny, Abhiram Natu, Muchun Niu, Jeffrey Ou, Nancy Parmalee, Alexi Runnels, Min-Hwan Sohn, Andrew B. Stergachis, Milovan Suvakov, Livia Tomasini, Melissa VonDran, Jeffrey M. Weiss, Shadi Zaheri, Yang Zhang, Chenghang Zong, Weichen Zhou.

### Declaration of generative AI and AI-assisted technologies

Portions of the text were refined using ChatGPT to improve clarity and flow of the English. All revised passages were subsequently reviewed to ensure that the original meaning and intent were preserved.

## Supplemental information includes

**Figure S1.** Benchmarking different assemblies (DSA, GRCh38 and T2T-CHM13) for somatic SNV discovery.

**Table S1.** Benchmark Data Generators.

**Table S2.** Comparison of biological and technical advantages (+) and disadvantages (-) of reference cell systems used for mutation benchmarking.

**Table S3.** Overview SMaHT Benchmarking Data generated by Centers (additional file).

**Table S4.** Benchmarking WGS and RNA-Seq Data Overview (additional file).

**Table S5.** Various QC checks performed on whole genome sequencing data at the SMaHT Data Analysis Center.

**Table S6.** Data access levels of the SMaHT benchmarking sequence data and derived data, including germline and somatic mutations, gene expression, and epigenetic profiles.

**Table S7.** Advantages and disadvantages of DSA and pangenome-based somatic mutation detection.

**Table S8.** Remaining challenges in discovering somatic mutations.

**Table S9**. Details of the variants confirmed in the COLO829BLT50.

**Table S10**. Details of the variants confirmed in the HapMap sample mixture.

**Table S11**. Details of the primers and probe designed to validate the variants in the HapMap sample mix.

## STAR★METHODS

### STUDY PARTICIPANT DETAILS

LB is a living 29-year-old male donor. ST001 is 22-year-old male. ST002 is a 74-year-old male donor. ST003 is a 22-year-old male. ST004 is a 73-year-old male.

Informed consent was obtained from LB participant according to the regulations of the Institutional Review Board (IRB protocol # 1104008337) and Yale Center for Clinical Investigation at Yale University.

Authorization for tissue donation for donors ST001 and ST002 was provided by the donor next-of-kin at a TPC-partnering organ procurement organization (OPO). Ethical approval for the tissue collection was obtained by National Disease Research Interchange (NDRI) through the University of Pennsylvania (IRB#5 FWA00004028). Frozen dorsolateral prefrontal cortex tissue from deceased donors ST003 and ST004 was obtained from the NIH NeuroBioBank repository at UMBTB. Collection of brain tissue from donors was performed as approved by the IRB of the University of Maryland, Baltimore and the University of Miami School of Medicine where the respective cases were recovered. Due to the authorization structure, only the age and sex of the benchmarking post-mortem donors can be made public.

## METHOD DETAILS

### Sample acquisition, preparation and distribution

#### Preparing HapMap samples and mixture

*contributed by Sheng Chih Jin and Benpeng Miao*

Each cell line was individually thawed, recovered, and expanded before pooling. Cells were cultured in T25 flasks containing 10 ml culture medium (84% RPMI-1640, 15% FBS, and 1% Glutamax), maintained within the optimal viable concentration range (2×10⁵ – 5×10⁵ cells/ml), and split by dilution every 3–5 days based on cell counts measured using a Vi-Cell Blu cell counter (Beckman Coulter) with trypan blue viability assessment. On the day of harvest, cells from each line were pooled into a 250 ml conical tube, thoroughly mixed, and approximately 10 ml was transferred to a T25 flask. Aliquots were taken for cell counting and sterility assessment; the latter was performed by streaking samples onto two sheep’s blood agar plates and incubating them for two weeks at 30°C and 37°C. Based on cell counts, volumes of individual cell suspensions were calculated to achieve the target mix ratio: 83.5% HG005, 10% HG02622, 2% each of HG002, HG02257, and HG02486, and 0.5% HG00438. The calculated volumes were combined in a secondary tube, centrifuged at 228 RCF for 10 minutes, resuspended in 800 ml cryopreservation medium (65% RPMI-1640, 30% FBS, and 5% DMSO), and dispensed into cryovials at 5×10⁶ cells/vial. Cryovials were then cryopreserved using a controlled-rate freezer and stored in liquid nitrogen vapor.

#### Preparing COLO829 samples

*contributed by Nancy Parmalee*

##### COLO829BLT50 Cell Culture and Mixing Methods

Pure populations of COLO829BL cells (cat # CRL-1980, lot # 70022927) and COLO829 cells (cat # CRL-1974, lot # 70024393) were obtained from ATCC (6/21/2023) and expanded. All cells were grown at 37°C with 5% CO_2_. COLO829BL lymphoblastic suspension cells were thawed in recovery media which consisted of 1 part RPMI-1640 (Fisher cat# 11875093) 15% fetal bovine serum (FBS, Fisher cat# 10082147) to 1 part 100% FBS. Cells were centrifuged in recovery media at 300 rcf for 5 minutes to remove DMSO. The pellet was resuspended in recovery media and plated in a T-25 flask with 5 mL total media for a volume of 0.2 mL media per cm^2^ to allow for maximum gas exchange. After approximately 2 days in culture cells were split by dilution into RPMI-1640 15% FBS and split by dilution daily thereafter. Flask sized was increased with expansion of the culture maintaining a volume of 0.2 ml media per cm^2^. When the culture reached a volume of 100mL it was transferred to a 500 mL Erlenmeyer cell culture flask and maintained with shaking at 90 rpm. The maximum volume of the culture was 200 mL per 500 mL flask. COLO829BL cells were harvested on a continuous basis. Cells were harvested by centrifugation in 50 mL conical tubes at 300 rcf for 5 minutes. The pellet was resuspended in 500 mL RPMI-1640 15% FBS. Days in culture ranged from 9-18 days. Cells were counted using the Countess cell counter (Invitrogen Thermo Fisher) with trypan blue (Fisher cat#15250061) and cryopreserved in freezing media consisting of RPMI-1640 15% FBS with 10% DMSO (Sigma Aldrich cat# D2650-100ML) at a cell density of 3 million viable cells per mL in 2 mL cryovials and stored in liquid nitrogen vapor phase.

COLO829 adherent melanoma cells were thawed and plated directly into a T-75 culture flask in 15 mL RPMI-1640 10% FBS. Media was changed approximately every 4 days. Cells were grown to near confluence and split by removing media, washing with PBS pH 7.2 (Fisher cat# 20012027), and trypsinizing for five minutes at 37° C using 0.05% trypsin EDTA (Fisher cat# 25300062). Volumes of both PBS and trypsin were 5 mL for T-75 flasks and 15 mL for T-175 flasks. COLO829 cells were harvested at confluence from 90 T-175 flasks on a single day at passage 4. Cells from all 90 flasks were combined and mixed to equilibrate flask-specific somatic mutations across the mixture. Cells were counted using the Countess cell counter with trypan blue and cryopreserved in freezing media consisting of RPMI-1640 10% FBS with 10% DMSO at a cell density of 3 million viable cells per mL in 2 mL cryovials and stored in liquid nitrogen vapor phase. 433 vials of 2 mL per cryovial were produced.

On a single day cryopreserved COLO829BL and COLO829 cells were thawed from the previously cryopreserved vials, kept on ice, and counted using the Countess cell counter with trypan blue. Based on the Countess readings the COLO829 and COLO829BL cells were mixed at a 1:49 ratio using viable cell counts. Viability of the COLO829 cells was 93% and viability of the COLO829BL cells was 91%. Cells were aliquoted into cryovials with constant swirling to maintain the homogeneity of the mix. A total of 380 vials were cryopreserved at a cell density of 2.55 million viable cells per mL for a total of 5.1 cells per 2 mL vial and stored in liquid nitrogen vapor phase. To assure homogeneity of the mixture DNA was isolated from the first vial aliquoted and the last vial aliquoted to test for differential settling of cells during the mixing process. Droplet digital PCR assays of the final mix showed no significant difference between the first and the last vial.

##### COLO829BLT50 Bio-Rad Droplet Digital PCR (ddPCR) Quality Control Methods

Bio-Rad–designed droplet digital PCR (ddPCR) assays were used to target 5 mutations present in COLO829 tumor cells detected: *MADD* p.S1620F, *MAP4K1* p.P298P, *RELN* p.E331K, *SCN11A* p.T340T, *ZNF217* p.P651S. Each assay was validated by a temperature gradient experiment to determine the annealing temperature for ideal cluster separation. All PCR reactions were set up in a UV-treated hood with positive airflow, and samples were run in quadruplicate. Each 22-µL PCR mixture contained 10 µL of ddPCR SuperMix, 1 µL of assay, 1 µL (10 ng) of DNA, and 10 µL of water.

Twenty microliters of the PCR mix were used to generate droplets using the Bio-Rad Droplet Generator. PCR was performed using the following parameters: 10 minutes at 95°C, followed by 40 cycles of 30 seconds at 94°C, 60 seconds at the annealing temperature of the assay (52.7°C for *MAP4K1* p.P298P, *SCN11A* p.T340T, *ZNF217* p.P651S and 55°C for *MADD* p.S1620F, *RELN* p.E331K), and 10 minutes at 98°C, and held at 12°C. We used the Bio-Rad QX200 Droplet Reader within 24 hours and analyzed the data with Bio-Rad Quantasoft software. Each run included a no-template control, WT control, and positive mutant control.

Samples were positive if the variant fluorescence was significantly different from the fluorescence of the WT control using 95% confidence intervals for total error. The total error is displayed by the Quantasoft software and defined as the greater of either the technical error (Poisson error) or the empirical error (standard error of the mean). Variant allele fractions (VAFs) were calculated as the concentration of variant droplets out of the total concentration of droplets containing at least one copy of variant or WT DNA.

#### Preparing iPSC and fibroblast samples

*contributed by Livia Tomasini*

##### Fibroblasts sample collection and iPSC generation

The skin biopsy was collected from the inner side of the left upper arm and fibroblast primary cultures were selectively expanded as previously described^32,84^ using the explant method and fibroblast culture media (DMEM high-glucose-based medium supplemented with 10% FBS, 1:100 L-glutamine, 1:100 N.E. amino acids, 1:100 Pen/Strep, and 10 ng/mL FGF2, all reagents from Gibco Life Technologies). Cells were passed twice using TrypLE Express (Gibco Life Technologies). One million cells per vial were cryopreserved using Recovery Cell Culture Freezing Medium (Thermo Fisher Scientific). iPSCs were generated using the Epi5 Episomal iPSC Reprogramming Kit (Thermo Fisher Scientific) following manufacturer’s recommendations. The iPSC lines were propagated using mTeSR1 media (Stem Cell Technologies) and passed using Dispase (Stem Cell Technologies). After expansion, iPSC lines were cryopreserved in Cryostor CS10 (Stem Cell Technologies).

##### LB-LA2 fibroblasts culture and expansion

One cryovial of the fibroblast cell line from individual LB, left arm location (line LB-LA2) was thawed in fibroblast culture media (DMEM high-glucose-based medium supplemented with 10% FBS, 1:100 L-glutamine, 1:100 N.E. amino acids, 1:100 Pen/Strep, and 10 ng/mL FGF2, all reagents from Gibco Life Technologies). After centrifugation at 1000 rpm for 4 minutes, cells were plated in a tissue culture treated 75 cm^2^ flask (Falcon). Cells were cultured at 37°C with 5% CO2 in room air and the medium was changed every three days. Cells were passed and expanded when reaching about 70% confluency using TrypLE Express (Gibco Life Technologies), following manufacturer’s recommendations. The initial 75 cm^2^ flask was passed after two days in culture and plated in three 75 cm^2^ flasks. Three days later, cells were passed to five 175 cm^2^ flasks using the same procedure. Four days later, the cells were passed to twenty 175 cm^2^ flasks. After five days, the cells were passed again to forty 175 cm^2^ flasks. After five days, the cells were trypsinized as above, the pellets were resuspended in fresh fibroblast culture media and counted using a manual hemocytometer. Cells were resuspended in Recovery Freezing media (Gibco Life Technologies) at a density of 3 million cells per mL. A total of 52 cryovials with 2 mL each (6 million cells per vial) were produced at passage 6 to minimize culture-acquired mutations. This procedure ensured that each site received the identical fibroblast cell suspension for sequencing.

##### LB-LA2 iPSC clones culture and expansion

iPSC clones derived from the fibroblast line LB-LA2 (LB-LA2 clone #1; clone #2; clone #4; clone #52 and clone #60) were thawed in mTeSR1 media (Stem Cell Technologies), centrifuged at 800 RPM for 4 minutes and each plated in a well of a 6-well tissue culture treated plate (Corning), pre-coated with a 1% solution of Geltrex LDEV-Free hESC-qualified Reduced Growth Factor Basement Membrane Matrix (Gibco Life Technologies). iPSC lines were cultured at 37°C, 5% CO2 and 5% O2. The media was changed every two to three days. After one week, the cells were passed using Gentle Cell Dissociation Reagent (Stem Cell Technologies) following manufacturer ‘s instructions. Each clone was plated on one 10cm diameter tissue culture dish (Falcon) pre-coated with Matrigel as above. Between 5 to 8 days later depending on the clone, cells were passed again following the same procedure, and each plated on four 10cm Matrigel pre-coated dishes. iPSCs were harvested between 7 to 9 days later depending on the clones. At the time of harvesting, iPSC colonies were dissociated using TrypLE Express (Gibco Life Technologies), resuspended in fresh mTeSR1 media and counted using a manual hemocytometer. The single cell suspensions were resuspended in Cryostor CS10 (Stem Cell Technologies) at a density of 3 million cells per mL and 2 mL were aliquoted per cryovial (6 million cells per vial). A total of four cryovials were produced for clone #4 at passage 6; a total of five cryovials for clone #1 and for clone #2, both at passage 6; a total of two cryovials for clone #60 at passage 5; and a total of three cryovials for clone #52 at passage 4.

##### Sample distribution

The shipping of the fibroblast sample to all GCCs was done at once. Cells for each of the 5 iPSC lines were shipped to Broad GCC for genomic characterization.

#### Human Tissue Collection, Preparation, and Distribution

*contributed by Kathryn Leonard, Melissa VonDran, Eric Ho, Thomas Blanchard, Thomas J. Bell*

Non-brain tissues for the SMaHT benchmarking study were recovered by the TPC from two post-mortem donors. Donors over the age of 18 years were considered eligible for inclusion in the study if tissues could be collected within 24 hours of death. Samples of liver, lung, and colon were recovered from each donor using SMaHT-specific recovery, processing, and preservation protocols at the OPO. Briefly, for each tissue, five 10 cm x 1 cm x 1 cm samples from defined anatomical locations in each organ were rinsed with saline, cut into several smaller pieces, placed in individual 50 ml conical tubes and snap frozen. A companion 1 cm3 sample from each tissue was preserved in 10% formalin, praffin-embedded, and stained with Hematoxylin and Eosin (H&E) for subsequent histological analysis. Each tissue was reviewed by a pathologist at the University of Maryland Brain and Tissue Bank (UMBTB) to evaluate tissue composition and general pathological features such as inflammation, and organ-specific pathological conditions. Frozen dorsolateral prefrontal cortex tissue from deceased donors ST003 and ST004 was obtained from the NIH NeuroBioBank repository at UMBTB.

##### Homogenized Tissue Samples

To minimize regional variability and ensure consistency across sequencing centers, frozen tissues were pulverized into a powder using a pre-chilled mortar and pestle with liquid nitrogen. The homogenized material was pooled, mixed for uniformity, and aliquoted into pre-weighed cryovials using pre-chilled spatulas, then stored at -80°C at the TPC for distribution to GCCs or TTDs.

##### Intact, Non-homogenate Tissue Samples

For experimental procedures that required intact tissue, pre-chilled forceps were utilized to breakdown intact frozen tissue samples (∼1 gram) into approximately 150mg aliquots. The intact tissues were aliquoted into pre-weighed cryovials using pre-chilled spatulas, then stored at -80°C at the TPC. Sample aliquots were distributed by TPC on dry ice to GCCs or TTDs.

##### Molecular Quality Control

RNA and DNA quality from snap-frozen tissue samples was examined by UMBTB prior to distribution of samples to sequencing centers. Tissues were lysed with ZrOBZO beads in a Bullet Blender (Next Advance) followed by isolation of RNA and DNA via QIACube using the RNeasy Lipid Tissue and DNeasy Blood and Tissue Kits, respectively, (Qiagen). RNA quality was assessed using the Agilent 2100 Bioanalyzer to generate RNA Integrity Numbers (RIN). Samples with RIN scores of 4.0 or higher were included for distribution and further analysis. DNA quality was assessed by measuring concentration and purity using a Nanodrop spectrophotometer. In addition, each GGC or TTD also performed quality control assays to confirm sample suitability for their experimental procedures.

##### Donor Metadata

The clinical data collected for each SMaHT donor belong to one of two categories: donor-level data or tissue sample-level data. Donor-level data encompass all clinical measures of the donor, including donor demographics (age, sex, height, weight, and body mass index (BMI)), medical history (cause of death and past medical history), social history (alcohol, tobacco, and illicit drug use), and serological testing results. Details on access to donor level data are contained within the official SMaHT data use policy available through the SMaHT web portal at (https://data.smaht.org).

##### Tissue Sample Metadata

Tissue sample-level data are attributes belonging to each sample collected and include the tissue type, ischemic time, comments from the prosector and pathology reviewer, and process metadata such as batch ID.

### Data generation

#### Sequencing at WashU-VAI

*contributed by Sheng Chih Jin, Robert Fulton, Ben Johnson, and Mary Majewski*

##### Long-Read Sequencing (PacBio HiFi SPRQ Chemistry)

PacBio HiFi SMRTbell libraries was prepared following PacBio protocol ‘Procedure & Checklist – Preparing Whole Genome and Metagenome Libraries Using SMRTbell Prep Kit 3.0’. Genomic DNA was fragmented with a mode of ∼20 kb using the Diagenode Megaruptor 3 instrument. Genomic DNA is initially processed with DNAFluid+ (P/N E07020001) using speed of 40 to dissociate aggregates and homogenize the DNA. The homogenized DNA was then sheared twice using Shearing kit (P/N E07010003) with speeds of 28 and 30. Sheared sample was assessed via fluorometry (Qubit High Senstivity DNA Kit) and Agilent Femto Pulse (Genomic DNA 165kb Kit). Libraries are made according to PacBio protocol utilizing barcoded adapters from the SMRTbell barcoded adapter plate 3.0 (PacBio P/N 102-009-200) to allow for multiplexing of samples during sequencing. Libraries were size selected using Sage PippinHT instrument and the 0.75% Agarose High-Pass 75E kit (P/N HPE7510) with a start size of 15000bp-17000bp. Size selected libraries were prepared for sequencing following instructions generated in PacBio SMRT Link v13.0 Sample Setup and utilizing PacBio Revio polymerase kit (P/N 102-817-600). Sequencing was performed on PacBio Revio sequencer with an ‘On Plate Concentration’ of 170pM-200pM. 3 SMRTcells were generated for a total of ∼100X/SMRTcell and 300Gb total coverage. Average Q-score values of Q33, Q33, and Q32 for the 3 SMRTcells was achieved.

##### Watchmaker Stranded Total RNA-seq

Total RNA integrity was determined using Agilent Bioanalyzer or 4200 Tapestation. Library preparation was performed with 100ng to 500ng of total RNA. Libraries were generated with Watchmaker Library Prep Kit with Polaris Depletion (Watchmaker). Briefly, ribosomal RNA was removed by an RNaseH method and purified with RNAClean beads (Beckman). mRNA was then fragmented in buffer by heating depending on RNA quality per protocol. mRNA was reverse transcribed to yield strand specific cDNA. A second strand and A Tailing reaction was performed to yield fragments with an A base added to the 3’ ends. Illumina sequencing adapters were ligated to the ends. Ligated fragments were then amplified per protocol using primers incorporating unique dual index tags. Fragments were sequenced on an Illumina NovaSeq X Plus using paired end reads extending 150 bases.

##### KAPA Hyper PCR-Free Library Prep

Genomic DNA samples were quantified using the Qubit Fluorometer. Genomic DNA (∼600-1000ng) was fragmented on the Covaris LE220 instrument targeting ∼375bp inserts. Fragmented DNA was size selected using 0.8X ratio of Ampure XP beads (Beckman Coulter) to remove fragments less than 300bp. Dual indexed libraries were constructed utilizing the KAPA Hyper PCR-free library prep kit (Roche Diagnostics, Cat # 7962371001). Full length custom adaptors were used during ligation (IDT, UDI/UMI configuration with 10bp UDIs and a 9bp UMI in the i7 position). Libraries were run with KAPA Library Quantification kit (Roche Diagnostics) to measure molar concentration. Libraries are sequenced on NovaSeq X using paired end reads extending 150bp. For this application targeting 500× coverage, 4 libraries were constructed and utilized to generate the >500× coverage.

##### Single-cell Total RNA-seq (STORM-seq)

Single cell suspensions from the benchmarking cell line mixtures were rapidly thawed in a 37C water bath and resuspended in increasing 1:1 dropwise volumes of warm (37C) flow buffer (HBSS with no divalent cations + 2% FBS + 25 mM HEPES) with gentle agitation. Resuspended cells were then pelleted at 300 x g at 4C for 5 minutes. Cells were washed in warm flow buffer and re-pelleted. Next, cells were counted and resuspended at 1 x 10^6 cells/mL in warm flow buffer containing 0.5 ug/mL DAPI for active viability surveillance during sorting. Single, live cells were index sorted into each well of a 384-well plate (Eppendorf) containing 2.17 uL Fragmentation Buffer (1.17 uL PBS pH 7.2 [Gibco 20012-027], 0.17 uL 10X Lysis Mix, 0.17 uL SMART scN6, 0.69 uL scRT buffer) using a BD FACSymphony S6 sorter running BD FACSDiva v9.1.3 and equipped with BD StepSort. The S6 was run with a 130 um nozzle at 14 psi. For cell deposition into 384-well plates, we used single cell sort mode. Single cell libraries were prepared using the STORM-seq kit (Takara Cat # 634751) and protocol (https://doi.org/10.5281/zenodo.15178455)^85^. Briefly, after sorting, plates were immediately transferred to a pre-heated thermal cycler, heated to 85C for 3 minutes and snap cooled on ice for 2 minutes. Once cooled, 1.17 uL First Strand Master Mix (0.75 uL SMART scTSO mix, 0.08 uL RNase Inhibitor, 0.34 uL SMARTscribe RT) is added to each well. Immediately prior to dispensing, ERCC RNA Mix 1 (Thermo Fisher) is added to the First Strand Master Mix to create a 1:1000000 final dilution. First Strand Synthesis is performed at 42C for 180 minutes, followed by 70C for 10 minutes and 4C hold. After 1st Strand Synthesis, PCR1 (10 cycles of amplification: 94C 1 min, [98C 15s, 55C 15s, 68C 30s]x10, 68C 2 min, 4C hold) was performed with the addition of SMARTer RNA Unique Dual Index Sets A-D (SMARTer RNA Unique Dual Index Kits 96U sets A-D, Takara Cat # 634752, 634753, 634754, 634755). 4.67 uL of PCR1 Master Mix (0.33 uL nuclease free water, 4.16 uL SeqAmp CB PCR buffer, SeqAmp DNA Polymerase) is added to each well. Each well then receives 1 uL of a Unique Dual Index (UDI). UDIs were diluted 1:4 in 10mM Tris-HCl, pH 8.0 (Teknova) prior to addition. The plate was pooled and a bead-based cleanup (Beckman Coulter AMPure XP Beads) was performed. 162 uL of rRNA Depletion Master Mix (123.12ul nuclease free water, 16.2 uL 10X ZapR buffer, 11.02ul scZapR, 11.02ul sc-R Probes) is used to elute the depleted cDNA from the dried beads. The eluate is incubated at 37C for 60 min, 72C for 10 min, 4C hold. Finally, 12 cycles of amplification (94C 1 min, [98C 15s, 55C 15s, 68C 30s]x12, 4C hold) were performed for PCR2 (PCR2 Master Mix (208 uL nuclease free water, 400 uL SeqAmp CB PCR Buffer, 16ul PCR2 primers, 16 uL SeqAmp DNA polymerase) and the final library was eluted in 20 uL 10mM Tris-HCl pH 8.0. After QC, where necessary, an additional bead clean-up was performed to remove any remaining adapter-dimer.

##### Short-Read Sequencing

For both Kapa Hyper PCR-free libraries as well as the Bulk RNA-seq libraries, the molarity of each library was accurately determined through qPCR utilizing the KAPA library Quantification Kit according to the manufacturer’s protocol (KAPA Biosystems/Roche) to produce cluster density appropriate for the Illumina NovaSeq X Plus instrument. Normalized libraries were sequenced on a NovaSeq X Plus Flow Cell using the151×10×10×151 sequencing recipe according to manufacturer protocol to generate a >500× WGS coverage, and >100M read pairs for the RNAseq library. For the STORM-seq libraries, sizing was performed using the Agilent Bioanalyzer HS kit and concentration was determined using a Qubit fluorometer and dsDNA HS kit before sequencing using an Illumina NovaSeq 6000 S2 2×150 bp flow cell. Libraries were sequenced to generate an average of 1M reads per cell.

#### Sequencing at UW

*contributed by Jeffrey Ou, Christian D. Frazar, Caitlin N. Jacques, Jeffrey M. Weiss*

##### Short-Read DNA (Illumina WGS)

Short-Read whole genome sequencing was carried out using the Illumina NovaSeqX Plus sequencer. DNA was extracted from cell lines and tissues using the Qiagen DNAEasy Blood and Tissue kit (69506). Lysis time varied by tissue type and ranged from 4 hours to overnight. DNA from 1mL blood samples were extracted using the Qiagen MagAttract HMW DNA kit (67563) following the manufacturer’s instructions. The QIAmp DNA Mini kit (51304) was used to purify DNA from buccal swabs, following the manufacturer’s instructions. The integrity of the extracted DNA was confirmed using the Agilent Femto Pulse instrument and the DNA yield was confirmed using the Invitrogen Qubit kit and Qubit 4 fluorometer. Starting with a minimum of 750ng of DNA, samples are sheared in a 96-well format using a Covaris R230 focused ultrasonicator targeting 380bp inserts. The resulting sheared DNA is cleaned with Takara NucleoMag beads to remove sample impurities prior to library construction. Shearing is followed by size selection and sample prep is performed using the KAPA Hyper Prep kit (KR0961 v1.14). End-repair, A-tailing, and ligation are performed as directed. Two final NucleoMag cleanups are performed after ligation to remove excess adapter dimers from the library. All library construction steps are automated on the Revvity Janus platform. Library yield is quantified using Invitrogen Quant-IT dsDNA High Sensitivity kit (Q33120). Libraries are validated in triplicate using the Biorad CFX384 Real-Time System and KAPA Library Quantification Kit (KK4824). Barcoded libraries are pooled using liquid handling robotics prior to loading. Massively parallel sequencing-by-synthesis with fluorescently labeled, reversibly terminating nucleotides is carried out on the NovaSeq X Plus sequencer. Base calls are generated in real-time on the instrument (RTA 4.29.2) and then demultiplexed, fastq files are produced by bcl-convert v4.2.7.

##### RNA Sequencing (PacBio bulk Kinnex)

Extracted total RNA was quality checked using UV-Vis spectroscopy (Denovix DS-11 FX) and Agilent Bioanalyzer 2100 using the Total RNA Nano 6000 kit (Agilent, G2939A & 5067-1511.) Kinnex full-length RNA libraries were generated per manufacturer’s recommendations (PacBio, 103-072-000). Samples were sequenced on the Revio platform on SMRT Cells 25M with Revio Chemistry V1 (PacBio, 102-817-900) or SPRQ (PacBio, 103-520-200) with Adaptive Loading and 30-hour movies. Data were postprocessed using SMRT Link v13.1 or 13.3 with the “Read Segmentation and Iso-Seq” pipeline to segment and classify reads.

##### Long-Read DNA (PacBio HiFi/Fiber-Seq)

Extracted DNA samples were checked for quantity using Qubit dsDNA HS (Thermo Fisher, Q32854) measured on DS-11 FX (Denovix) and size distribution using FEMTO Pulse (Agilent, M5330AA & FP-1002-0275.) Depending on initial length distribution, samples were sheared to a target peak length of ∼20 kbp. Some samples were left unsheared; others were sheared with Megaruptor 3 Hydropores (Diagenode, B06010003 and E07010003.) Moderately degraded or low-concentration samples were subjected to a light shear at setting 28 or 29, while intact DNA samples were sheared by processing twice, at settings 28 or 29 and 30 or 31. Highly intact and non-homogeneous DNA samples (e.g. derived from fibroblasts) were pre-sheared with Megaruptor 3 DNAFluid+ (Diagenode, E07020001) before final shearing. Sheared DNAs were subjected to PacBio HiFi library prep via the SMRTbell Prep Kit 3.0 (PacBio, 102-182-700) using barcoded adapters (PacBio, 102-009-200). When final library QC permitted (minimum of 500 ng of individual or pooled libraries,) size selection was performed with Pippin HT using a high-pass cutoff of 8-15 kbp (Sage Science, HTP0001 & HPE7510.) Low-mass libraries or those with short final size distributions (∼5-10 kbp) were instead treated with a mild size-selection using diluted AMPure PB beads per the manufacturer’s protocol. Libraries were sequenced on the Revio platform on SMRT Cells 25M with Revio Chemistry V1 (PacBio, 102-817-900) or SPRQ (PacBio, 103-520-200) with Adaptive Loading and 30-hour movies.

##### Long-Read DNA (ONT Standard WGS)

DNA was extracted from tissue samples using the NEB Monarch HMW DNA extraction kit for Tissues (#3060L). No more than 25mg of tissue was cut into small chunks, then placed into a 1.5mL tube and mashed with the pestle. The sample was lysed according to the protocol at a shaking speed of 1000rpm (liver at 2000rpm; colon at 2000rpm for 15min, no shaking 30min). After lysis and RNAse treatment, brain samples were put on ice for 3 min before protein precipitation and phase separation. 800uL of the upper phase is combined with isopropanol to precipitate the DNA. DNA washes and elution followed the manufacturer’s protocol. DNA was quantified by Qubit and size distribution was measured on the Femto Pulse. In order to selectively remove smaller fragments of DNA, we performed a short read eliminator (PacBio 102-208-400) and cleaned up the DNA with a bead wash. Libraries were constructed using between 2-9ug of DNA and the Ligation Sequencing Kit from ONT (SQK-LSK114) with modifications to the manufacturer’s protocol. End repair was incubated for 20 minutes, and the adapter ligation was incubated for 1 hour. The final library was eluted in 30ul of EB and quantified by Qubit. 200-400ng of library was loaded onto a FLO-PRO114M R10.4.1 flow cell for sequencing on the PromethION, with two nuclease washes and reloads after 24 and 48 hours of sequencing.

##### Long-Read DNA (ONT Ultra-long WGS)

Ultra-high molecular weight DNA was extracted from fibroblast cell lines using a phenol chloroform extraction protocol (Logsdon, protocols.io, 2020) or using the NEB Monarch HMW DNA extraction kit for Cells & Blooc (#T3050L) following the manufacturer’s protocol with the following exceptions: 6 million cells was used for the starting input with a shaking speed of 600 rpm during the lysis step. DNA was precipitated with 300uL EEB (ONT) and solubilized at 4°C for two days. Libraries were constructed using the Ultra-Long DNA Sequencing Kit V14 (SQK-ULK114) following the manufacturer’s protocol. For monarch DNA, two extractions were combined for library preparation and final elution volume was doubled from the protocol. For phenol chloroform extracted DNA, approximately 40ug of DNA was input into library prep and the final elution volume ranged from 2x-4x the protocol volume, depending on DNA visualized during the clean-up step. Final libraries were left at room temp over night to solubilize. 75 uL of library was loaded onto a primed FLO-PRO114M R10.4.1 flow cell for sequencing on the PromethION, with two nuclease washes and reloads after 24 and 48 hours of sequencing.

##### Hi-C DNA Sequencing

Cultured fibroblasts were harvested and counted. 1-2M cells were washed and frozen dry at -80°C. Omni-C library preparation was conducted using either the Dovetail Omni-C Kit (PN 21005G Dovetail® Omni-C® Kit) for human cells v2.0 or Dovetail Omni-C Kit for human cells v2.1 (updated July 2025) with minor modifications. Cell pellets were treated with 0.5-2uL of the Omni-C nuclease mix. Sample digests were reviewed and compared against the recommended range in the Omni-C protocol prior to proceeding with proximity ligation. 150-200ng of proximity-ligated sample was used for library preparation. The number of PCR cycles used for library preparation was dependent upon the amount of starting material from proximity ligation and ranged between 9-12 cycles. The final library was sequenced on an Illumina NovaSeqX platform and generated at least 1.2 billion 2×150 read pairs.

##### Data used to create DSA

To generate DSA for COLO829BL, we used a combination of PacBio HiFi sequencing data, ONT ultra-long WGS data, and Hi-C. To generate DSA for the fibroblast (LB-LA2) and benchmarking donor samples (ST001–ST004), we leveraged PacBio HiFi-only assemblies. Where short-read Illumina sequencing data were available for a given sample, these data were used to calculate the quality value (QV) of that sample’s genome assembly.

#### Sequencing at BCM

*contributed by Chris Grochowski, Hsu Chao, Muchun Niu, Ziad Khan, Heer Mehta, Donna Muzny, Rui Chen, Chenghang Zong, Harsha Doddapaneni*

##### DNA Extraction

DNA from different sample sources (Cell lines, Tissue homogenates, Blood and Tissue Cores) was extracted using the Chemagic Prime 8 robot and Chemagen’s proprietary Magnetic Bead technology, with the Chemagic Prime DNA Blood kit and the manufacturer’s 2k protocol (Revvity).

##### RNA Extraction

RNA from cell lines material was extracted using the Chemagic Prime 8 robot and Chemagen’s proprietary Magnetic Bead technology, with the Chemagic Prime Total RNA Blood 2.5k Kit (Revvity). RNA from other sample sources (Tissue homogenates, Blood and Tissue Cores)_was extracted according to the manufacturer’s Trizol protocol manually. The isopropanol/aqueous phase was then transferred to a RNeasy spin column (Qiagen), and the manufacturer’s RNeasy protocol was then followed.

##### Short read WGS sequencing

Two independent methods (Picogreen assay and 1% E-gels)) were used to determine the quantity and quality of the DNA before library construction. For library preparation, DNA (1 ug) was sheared into fragments of approximately 450-600 bp in a Covaris E220 system (Covaris, Inc. Woburn, MA) in batches of 96 samples at a time, followed by double SPRI bead clean up to select a narrow band of sheared DNA for library preparation. DNA was end-repaired, 3’-adenylated and ligated using a set of 96 8-bp adapters (Illumina TruSeq UD Indexes v2, # 20040870) for sample barcoding. The final library size estimation and quantification are completed using the Fragment Analyzer (Agilent AAT, Inc) electrophoresis system and QuantStudio™ 6 Flex Real-Time PCR System (Applied Biosystems) respectively to achieve an average final library size of ∼530bp and must be greater than 470 bp. Libraries were sequenced on The NovaSeqX instrument to generate 150 bp, dual indexed and paired-end sequence reads in a format of multiplexed pools to generate 412× - 541× coverage.

##### Mapping and QC pipeline for internal quality assessment before data submission to DAC

WGS sequence data were aligned to the hg38 reference genome, followed by variant calling using lllumina’s Dynamic Read Analysis for GENomics (DRAGEN) software, v4.3.6. Genome coverage was evaluated by calculating the mean coverage and the distribution of coverage across the genome, including the proportions of bases covered at ≥1×, ≥10×, and ≥20×. The proportions of bases meeting these thresholds and the total number of mapped bases at Q20 or higher were reported for internal QC tracking. Alignment data were assessed for contamination using VerifyBamID v1.1.3 and an orthogonal confirmation of sample identity was applied using the Error Rate In Sequencing (ERIS) software developed at the HGSC to rapidly compare sequence data to genotypes from SNP arrays via an “exact match” test.

##### Bulk RNA Sequencing

Whole transcriptome sequencing (total RNAseq) data was generated using the Illumina TruSeq Stranded Total RNA with Ribo-Zero Globin kit (20020612, Illumina Inc.) or Watchmaker RNA Library Prep Kit with Polaris® Depletion (7BK0002-096, Watchmaker Genomics). RNA quality and quantity was estimated using Agilent Bioanalyzer. To monitor sample and process consistency, 1 µl of the 1:50 diluted synthetic RNA designed by External RNA Controls Consortium (ERCC) (4456740, ThermoFisher) was added to 1 µg total RNA. In addition, as a process control, the Universal Human Reference RNA (UHR) (740000, Agilent Inc.), was processed in parallel with the RNA samples. Libraries were sequenced on the NovaSeq 6000 instrument using the S4 reagent kit (300 cycles) to generate 2×150bp paired-end reads. In order to generate a minimum of 157-300M read-pairs per sample.

##### Mapping and QC pipeline for internal quality assessment before data submission to DAC

The RNA-Seq analysis pipeline cleans and processes raw RNA sequencing data (FASTQs), providing robust QC metrics and has the flexibility to map the reads to either GRCh37 reference or GRCh38 (after excluding the alternate contigs). The pipeline aligns RNA-Seq reads, removes duplicates, and generates QC metrics. It also quantifies gene expression using RSEM, generates quality control metrics, and produces raw gene feature counts.

##### Long read WGS

WGS on ONT platform: Genomic DNA was quantified using Qubit dsDNA quantification broad range assay (Thermo Fisher Scientific). DNA size was determined using the Femto Pulse System (Agilent). A total of 3 libraries was prepared per sample to be sequenced on 3 flow cells with the aim of achieving 90× coverage. DNA was sheared using Covaris g-tubes (Covaris 520079) to achieve an average size of 20-25 kb for DNA from cell lines and 15-20 kb for DNA from tissues. Sheared DNA was size selected on the PippinHT instrument (Sage Science) using the 6-10 kb or the 15-20 kb High-Pass definition. ONT libraries were prepared using the SQK-LSK114 kit following the manufacturer’s instructions. Final libraries were eluted in 27 µL of the ONT elution buffer and quantified using the Qubit dsDNA quantification broad range assay. Libraries were loaded at 15 fmoles to be sequenced on R10.4.1 flowcells.

##### Mapping and QC pipeline

For each ONT sequencing event, the unaligned bams were consolidated into a single fastq, followed by alignment to hg38 reference genome using minimap2 v2.24. Additional sequencing is performed if the minimum mean genome coverage is not met. For sequencing data quality control, alignment data were assessed for sex concordance, contamination using VerifyBamID v1.1.3, and an orthogonal confirmation of sample identity using the HGSC’s ERIS software. For additional downstream secondary analyses, we used Sniffles2 v2.0.5 for SV calling, Clair3 v3.0.1.11 for SNV calling, and WhatsHap v2.3 for phasing. These tools were conveniently executed via Princess2, a highly configurable open workframe. Post analysis, select files were archived. Our pipeline was implemented on our local high-performance compute cluster. All ONT data contained MM and ML methylation tags, which described the location of the base modifications and the likelihood of the modification (i.e. 5hmC, 5mC, or no modification) at each given location. Deliverables included ONT consolidated unaligned bams and fastqs.

##### WGS on PacBio platform

Genomic DNA was quantified using Qubit dsDNA quantification broad range assay (Thermo Fisher Scientific). DNA size was determined using the Femto Pulse System (Agilent). A total of 2 libraries were prepared for each sample to be sequenced across 3 SMRT Cells to achieve 90× coverage. DNA was sheared using Covaris g-tubes (Covaris 520079) to achieve an average size of 18-22 kb.Sheared DNA was size-selected on the PippinHT instrument (Sage Science) using the 6-10 kb or the 15-20 kb High-Pass definition. The size selected DNA was used as input for Pacbio SMRTBell Prep Kit 3.0 for library preparation. DNA damage repair, A-tailing and adapter ligation were performed as per manufacturer’s instructions. SMRTBell Adapter Index Plate 96A was used for barcoding each library. Adapter ligated DNA was nuclease treated following the manufacturer’s guidelines and purified using 1X Pacbio SMRTBell Cleanup beads. Final libraries were eluted in 30 µL Pacbio elution buffer and quantified using the Qubit dsDNA quantification high-sensitivity assay (Thermo Fisher Scientific). Final library size was determined using the Agilent Femto Pulse.Sequencing primer annealing and polymerase binding were performed using the Revio Binding Kit 3.0. Libraries were loaded onto the PacBio Revio machine utilizing SMRTlink v13 for workflow setup and loaded at 325pM loading concentration with 30 hours of movie time.

##### Mapping and QC pipeline for internal quality assessment before data submission to DAC

For PacBio, the PacBio unaligned HiFi bams were demultiplexed and aligned to hg38 reference genome using PacBio’s SMRTLink v13 tools lima v2.9.0 and pbmm2 v1.13.1, respectively. Additional sequencing is performed if the minimum mean genome coverage is not met. For sequencing data quality control, alignment data were assessed for sex concordance, contamination using VerifyBamID v1.1.3, and an orthogonal confirmation of sample identity using the HGSC’s ERIS software. For additional downstream secondary analyses, we used Sniffles2 v2.0.5 for SV calling, Clair3 v3.0.1.11 for SNV calling, and WhatsHap v2.3 for phasing. These tools were conveniently executed via Princess2, a highly configurable open workframe. Post analysis, select files were archived. Our pipeline was implemented on our local high-performance compute cluster. All PacBio data contained MM and ML methylation tags, which described the location of the base modifications and the likelihood of the modification (i.e. 5hmC, 5mC, or no modification) at each given location. PacBio unaligned HiFi bams were delivered.

Key QC metrics and thresholds have been established on both platforms (mean coverage, % genome coverage at 10×, HIFI yields, mean read length, contamination, and Q30 or Q10 Mapped bases) and sequence data is screened for this before submission to DAC.

##### Single cell RNA sequencing

Frozen samples were minced on dry ice, lysed in a nuclei extraction buffer, and dissociated using gentleMACS. Nuclei were filtered, centrifuged, resuspended, and triturated. After magnetic labeling with anti-nucleus beads, nuclei were purified using column separation, washed, and resuspended for counting, morphology assessment, and downstream library preparations. The library preparation and sequencing of single-nuclei cDNA were carried out following the manufacturer’s protocols (https://www.10xgenomics.com). To obtain single cell GEMS (Gel Beads-In-Emulsions) for the reaction, single-nuclei suspension was loaded onto a Chromium X. The library for single nuclei RNA-seq was prepared with the Chromium GEM-X Single Cell 5’ Reagent Kits v2 (10x Genomics), while the library of single nuclei ATAC-seq was prepared with the Chromium Next GEM Single Cell ATAC Reagent Kits v2 (10x Genomics). The constructed libraries were subsequently sequenced on an Illumina Novaseq 6000 (https://www.illumina.com). The sequencing data were processed using the 10X Genomics software (Cell Ranger v7.2.0 for gene expression (GEX) and Cell Ranger ATAC v2.1.0 for ATAC-seq analysis).

##### Droplet Digital PCR (ddPCR) Validation

A competitive TaqMan probe approach was used to validate the sample concentrations for the COLO829BLT50 as well as the HapMap sample mixture using the method we recently described **(Tables S9, S10, S11)**^86^. To assess the level of COLO829BL versus COLO829 we confirmed the presence and fraction of multiple well-known somatic variants from the COSMIC database within the mixture (**Table S9**).

Five BioRad designed ddPCR Mutation Assays targeting variants identified in the COLO829 cell line were ordered from Bio-Rad Laboratories (Hercules, CA). The assays targeted the following variants: MAP4K1 p.P298P (c.894C>T; Assay ID dHsaMDS546407091), RELN p.E331K (c.991G>A; Assay ID dHsaMDS550923980), MADD p.S1620F (c.4859C>T; Assay ID dHsaMDS851278352), SCN11A p.T340T (c.1020G>A; Assay ID dHsaMDS432990804), and ZNF217 p.P651S (c.1951C>T; Assay ID dHsaMDS581392264). Each assay was run using standard ddPCR protocols and run on a Bio-Rad QX200 Droplet Digital PCR System^86^. Droplets were generated using the QX200 Droplet Generator, thermal cycled under recommended conditions, and read on the QX200 Droplet Reader. Data were subsequently analyzed using QuantaSoft software.

#### Sequencing at NYGC

*contributed by Alexi Runnels*

##### DNA/RNA Dual Extraction

DNA and RNA for all benchmarking samples were extracted using Qiagen’s AllPrep kit (80204), using manufacturer instructions. For RNA samples with low RIN output from AllPrep extractions, additional RNA was extracted using Qiagen’s RNeasy Mini kit (74106).

##### HMW DNA Extraction

HMW DNA for long read WGS was extracted using New England Biolabs’ Monarch HMW Extraction kit (T3050L and T3060L).

##### Short read WGS Library Prep

Whole genome sequencing (WGS) libraries were prepared using the NEBNext Ultra II FS DNA PCR-free Library Preparation Kit (NEB E7430L) in accordance with the manufacturer’s instructions. 500ng of DNA was sheared enzymatically and was subsequently end-repaired and adenylated. DNA fragments were ligated to Illumina sequencing adapters and the libraries underwent bead-based size selection. Final libraries were quantified using the QuantStudio5 Real-Time PCR System (Applied Biosystems) and Fragment Analyzer (Agilent).

##### Bulk RNA Library Prep

mRNA libraries were prepped using Illumina TruSeq Stranded mRNA Library prep (Illumina 20020595), in accordance with manufacturer recommendations, and using IDT for Illumina custom 10nt Indices (IDT). Briefly, 500ng of total RNA was used for purification and fragmentation of mRNA. Purified mRNA underwent first and second strand cDNA synthesis. cDNA was then adenylated, ligated to Illumina sequencing adapters, and amplified by PCR (using 10 cycles). The cDNA libraries were quantified using Fragment Analyzer (Agilent) and Spectramax M2 (Molecular Devices). Libraries were sequenced on an Illumina NovaSeq sequencer, using 2 x 100 bp cycles. FASTQ files were provided to the DAC, for further analysis.

##### Long-Read WGS Library Prep

High molecular weight (HMW) DNA samples were first sheared using the Megaruptor 3 (Hologic Diagenode, catalog number B06010003) to a target fragment size of 45 kB, following the manufacturer’s recommendations. Fragmented DNA quality was assessed using NanoDrop (ND-2000), Qubit Broad Range Assay (Q32850), and the Genomic TapeStation (G2964AA), all according to manufacturer instructions.

To improve 260/230 ratios, a 3X buffer exchange clean-up was performed using AMPure XP Beads for DNA Cleanup (A63882). The cleaned, fragmented HMW DNA was then prepared using the Oxford Nanopore Ligation Sequencing Kit V14 (SQK-LSK114), following the manufacturer’s protocol. Final libraries were quantified again using the Qubit Broad Range Assay (Q32850) and Genomic TapeStation (G2964AA).

##### Illumina Sequencing

Short read WGS libraries were sequenced on the Illumina NovaSeq X Plus, using 2 x 150 bp cycles. Bulk mRNA libraries were sequenced on the Illumina NovaSeq X Plus, using 2 x 100 bp cycles. FASTQ files were provided to the DAC, for further analysis.

##### Oxford Nanopore Sequencing

HMW libraries were sequenced on Promethion P24 (PRO-SEQ024) using R10.4.1 flow cells (FLO-PRO114M) according to manufacture instructions. FASTQ and unaligned BAM files were provided to the DAC, for further analysis.

#### Sequencing at Broad

*contributed by Shadi Zaheri*

##### Long-Read DNA (PacBio HiFi)

High-fidelity long-read WGS was performed using the PacBio Revio platform. DNA was extracted from tissues and cell lines using Qiagen’s MagAttract HMW DNA Kit. Libraries were prepared with the SMRTbell prep kit 3.0, size selected via Sage Science’s Pippin HT to ∼15kb, and sequenced on the PacBio Revio. The resulting raw data files were processed through circular consensus error correction. Sequencing was performed to a target depth of 24X (single-cell derived iPSC lines) or 96X (COLO829BLT50, tissues, HapMap) per sample, with average HiFi read lengths of 14.8 kb. The data were aligned to GRCh38 (no ALT contigs) using pbmm2 (v1.13.0).

##### Short-Read DNA (Illumina WGS)

Short-read whole genome sequencing was carried out using the Illumina NovaSeq X plus 10B platform. DNA was extracted from tissues and cell lines using the Qiagen AllPrep DNA/RNA/miRNA Universal Kit. Libraries were prepared with Kapa Biosciences HyperPrep library construction kit. Each sample was sequenced to an average depth of 80× across two libraries, achieving a combined coverage of 160× using Illumina 2×150 bp paired-end read chemistry. Reads were aligned to GRCh38 using DRAGEN aligner through the Sentinel pipeline.Short-Read RNA-seq (Illumina).

##### Short-Read RNA

RNA-seq was performed on total RNA extracted from tissues and cell lines using the Qiagen AllPrep DNA/RNA/miRNA Universal Kit. RNA integrity was assessed using the Caliper LabChip GX. Libraries were prepared using either Illumina’s TruSeq Stranded mRNA HT Kit or the Watchmaker RNA Library Prep Kit. TruSeq libraries were sequenced on the Illumina NovaSeq SP platform using 2×151 bp paired-end reads to a target depth of 75 million read pairs per sample. Watchmaker libraries were sequenced on the Illumina NovaSeq X Plus 25B platform using 2×146 bp paired-end reads to a target depth of 100 million read pairs per sample. Alignment was performed using the DRAGEN aligner (v07.031.677) against the GRCh38 reference genome.

#### Sequencing at Yale

*contributed by Livia Tomasini*

iPSC lines from LB donor have been previously sequenced by BGI to a coverage of 30-60× and the data has been made accessible at NDA (study no. 1057). DNA for the fibroblast sample was extracted using the DNeasy Blood & Tissue kit (Qiagen) following manufacturer’s instructions for cultured cells. Whole genome sequencing (WGS) libraries were prepared using the Elevate Mechanical Library Prep Kit (Element Biosciences) PCR-free. Genomic DNA was sheared using a Covaris LE220 sonicator. A double-sided SPRI (Solid Phase Reversible Immobilization) selection method was used to target a fragment size of 600 bp. DNA fragments were subsequently end-repaired, adenylated, and ligated using adaptors from the Elevate Long UDI Adapter Kit (Element Biosciences). Libraries were evaluated using qPCR. Four libraries were sequenced on an Element AVITI sequencing system using 2×150bp read length to an average depth of 220× genome coverage.

#### Generating MEI targeted sequencing data

*contributed by Eunjung Alice Lee and Weichen Zhou*

MEI-targeted sequencing methods, eHAT-seq on short reads and TEnCATS on long reads, were also benchmarked in HapMap mixture experiments. Somatic calls identified from WGS data were further orthogonally validated using these targeted sequencing approaches. Full experimental details, including sample preparation, sequencing data generation, and MEI analyses are provided in the SMaHT MEI benchmarking manuscript^21^.

For eHAT-seq, libraries were prepared from 500 ng genomic DNA using an optimized protocol based on the original HAT-seq method^87^, with major improvements to both experimental procedures and the analytical pipelines. Briefly, DNA was fragmented, end-repaired, A-tailed, and adapter-ligated, followed by L1 enrichment PCR to amplify the L1-genome junctions. To improve sequence diversity of the semi-amplicon library and sample labeling, L1HS-targeting primers with variable staggers and unique dual indexes were introduced during L1 enrichment and indexing PCR, respectively. The resulting stranded eHAT-seq libraries were size-selected, quantified, pooled, and sequenced on an Illumina NovaSeq platform (2 × 150 bp). Reads were trimmed, and read-pairs containing L1Hs sequences in Read 2 were retained. Corresponding Read 1 (representing 3’ genomic flanks) were aligned to hg38/GRCh38, and candidate insertion sites were identified by peak calling and annotation against reference MEI databases and genomic features.

The TEnCATS framework was previously described in McDonald et al.^76^ and further optimized here. Libraries were generated from HapMap mixture genomic DNA using an optimized protocol for the SQK-LSK114 kit and ONT R10.4.1 flow cells for targeting both L1Hs and Alu elements. Briefly, genomic DNA was dephosphorylated and subjected to Cas9-mediated target enrichment using ribonucleoprotein complexes assembled from in vitro transcribed guide RNAs, followed by cleavage, A-tailing, adapter ligation, bead purification, and long-fragment cleanup. Libraries were sequenced for 72h on MinION and PromethION2 Solo instruments and a wash and reload was completed at 24 hours for PromethION libraries. Reads were processed with NanoPal, aligned to the reference genome, and analyzed to identify non-reference MEI insertions. On-target reads for L1Hs and Alu elements were defined by BLASTn, clustered by locus, and filtered to exclude looping and chimeric artifacts by Minimera, with only calls supported by at least two reads retained.

Somatic calls identified from WGS data were further orthogonally validated using these targeted sequencing approaches. Full experimental details and benchmarking results from HapMap mixture experiments, including sample preparation, sequencing data generation, and MEI analyses are provided in the SMaHT MEI benchmarking manuscript^21^.

#### Generating duplex sequencing data

*contributed by Shadi Zaheri, Chenghang Zong, Harsha Doddapaneni, Nisrine T. Jabara, Gilad Evrony, Sangita Choudhury and Alexi Runnels*

##### Duplex sequencing experimental methods

Four donor tissue samples (ST001, ST002, ST003, and ST004), a COLO829 cell line, and a HapMap DNA mixture underwent duplex sequencing using the following six methods: CODEC, NanoSeq (HpyCH4V and MNB), CompDuplex-seq, HiDEF-seq, ppmSeq, and VISTA-seq. Detailed methods are covered in the companion paper^22^; and below is the brief description of the methods used for data generation.

##### CODEC

Libraries were sequenced on an Illumina NovaSeq S4 platform, targeting approximately 1× duplex coverage. CODEC library preparation and data processing followed previously described methods^80^. Two distinct end repair and dA-tailing strategies were implemented to suppress error propagation during library preparation. An enhanced version of this protocol (DRv2, manuscript in preparation) was also used to further minimize damage-related artifacts. Following ER/dA tailing, CODEC specific quadruplex adapters were ligated. Libraries were PCR-amplified and sequenced on the Illumina NovaSeq 6000 (S4 flow cells).

##### Compduplex and Nanoseq (HpyCH4V and MNB)

NanoSeq (HpyCH4V and Mung Bean) libraries for the benchmark materials were prepared using 50-100ng of DNA as described by Abascal et al.^88^ with minor changes as described in Chao et al. https://www.protocols.io/view/optimized-mung-bean-nuclease-nanoseq-library-prepa-d4h38t8n.html. Libraries were pooled and sequenced on Illumina NovaSeq sequencing platforms for 30× coverage. Analysis details are described in the Duplex sequencing companion manuscript. CompDuplex libraries were prepared using 20-50ngof gDNA as described by Niu and Zong (https://www.protocols.io/view/compduplex-accurate-detection-ofsomatic-mutations-kxygx3×4og8j/v1). Libraries were pooled and sequenced on Illumina NovaSeq sequencing platforms for 30× coverage. Analysis details are described in the Duplex sequencing companion manuscript.

##### HiDEF-seq

Genomic DNA was isolated from tissue and cultured cell samples using either the PacBio Nanobind PanDNA kit with Proteinase K and RNase A incubations at 37 °C or the Qiagen Puregene system with Proteinase K**/**RNase A incubations at room temperature. Libraries were prepared with an updated HiDEF-seq v3 workflow designed to streamline the procedure and provide genome-wide coverage via random fragmentation^79^. A detailed step-by-step protocol is available at: https://www.protocols.io/view/hidef-seq-kxygxy9mwl8j/v3. To reduce single-strand mismatch artifacts, we used a non-A-tailing strategy with blunt-end ligation to hairpin adapters. Short fragments were removed prior to fragmentation using the Short Read Eliminator (SRE) kit (PacBio), DNA was sheared to ∼4 kb and then end-repaired. Nicks were ligated with E. Coli DNA ligase, residual nicks were blocked with dideoxy-C/G/T nucleotides, hairpin adapters were ligated, and residual non-circularized molecules were depleted by exonuclease treatment. Final libraries were run on a PacBio Revio instrument with 30-hour movie times and with per-strand output settings.

##### VISTA-seq

VISTA-seq adapts the META-CS framework for duplex consensus sequencing to accommodate inputs ranging from single cells to bulk DNA containing many alleles. A detailed A detailed step-by-step protocol is available here https://www.protocols.io/view/tn5-duplex-sequencing-tn5-duplex-seq-for-low-input-6qpvr3nbzvmk/v1. After gentle lysis and cleanup, genomic DNA was tagmented with engineered Tn5 preloaded with strand-specific adapters, enabling simultaneous fragmentation and symmetric tagging of both strands. Strand-specific barcodes were then introduced through two rounds of indexed synthesis with high-fidelity polymerase, each followed by exonuclease cleanup to remove excess primers and reduce background. The resulting duplex-tagged libraries were PCR-amplified, purified, and size-selected to enrich ∼400–600 bp fragments. Sequencing was performed as 150 bp paired-end reads on the Illumina NovaSeq X Plus platform with a PhiX spike-in for run quality calibration.

##### ppmSeq (Ultima) sequencing

ppmSeq (Paired Plus Minus Sequencing) libraries were prepared using the NEBNext UltraShear module (NEB M7634L) and NEBNext Ultra II DNA PCR-free Library preparation Kit (NEB E7410L) in accordance with the manufacturer’s instructions. 250ng of DNA was sheared enzymatically and was subsequently end-repaired and adenylated. DNA fragments were ligated to Ultima Genomics Indexed Native Duplex Adapters and the libraries underwent bead-based size selection. Final libraries were quantified using the QuantStudio5 Real-Time PCR System (Applied Biosystems) and Fragment Analyzer (Agilent). The ppmSeq libraries were sequenced on Ultima Genomics Solaris (UG100) instrument according to manufacturer instructions.

#### Primary template-directed amplification (PTA)

*contributed by Joe Luquette, Abhiram Natu, and Milovan Suvakov*

Full experimental details sample preparation and data generation for PTA analyses are provided in a dedicated manuscript^23^. Single nuclei were isolated from frozen post-mortem lung and colon tissues and amplified by primary template-directed amplification (PTA) across four TTDs from Yale University, Boston Children’s Hospital (BCH), Weill Cornell College (Conell), and Mayo/Yonsei University. Sample input was frozen tissue cores at all sites except BCH, which used tissue homogenate.

The groups employed different nuclei preparation strategies. Tissue was dissociated mechanically using Dounce homogenizers or the Singulator 100 automated platform. Nuclei extraction was done in sucrose-based buffers or using Minute detergent-free nuclei isolation kit (Invent Biotechnologies) with NP40 buffer. FACS-based single-nucleus sorting was performed at all sites, but DNA staining varied and included DAPI, PI, AO/PI, SYTO9, or no staining. Whole-genome amplification was performed using the BioSkryb ResolveDNA PTA kit at all centers: V1 was used at Yale, BCH, and Cornell, while Yonsei used both V1 and V2. Amplification quality was assessed by a 4-locus multiplex PCR assay at Yale, BCH, and Yonsei, with only samples yielding all four amplicons (and DNA concentrations exceeding 100 ng by Qubit) retained for sequencing.

A total of 102 single nuclei (27 by Yale University, 20 by BCH, 7 Cornell, and 48 Yonsei University) were sequenced at Baylor College of Medicine, the Broad Institute, NYGC, WashU-VAI, and Theragen Bio. Sequencing library preparation varied across GCCs. All libraries were sequenced on Illumina NovaSeq X or NovaSeq X Plus platforms with 2×150 bp paired-end reads, targeting approximately 30X coverage, although final coverage varied per cell. Reads were uniformly processed at DAC as described in the following section.

### Data Processing

#### Data Processing at the Data Analysis Center (DAC)

*contributed by Michele Berselli and Elizabeth Chun*

##### Illumina paired-end whole genome sequencing (WGS) data

were processed using a standard pipeline based on the Genome Analysis Toolkit (GATK) Best Practices. Practices^89–91^. Reads containing artifactual poly-G repeats associated with Illumina two-color chemistry were initially detected and discarded using fastp^92^ (v0.23.2). The remaining reads were aligned to the GRCh38 reference genome (GCA_000001405.15), which excludes ALT contigs and decoy sequences from hs38d1, using BWA-MEM^93^ (v0.7.17) as implemented in the Sentieon software^94^ (v202308.01). This specific version of GRCh38 was chosen, because inclusion of ALT contigs, large variations with very long flanking sequences nearly identical to the primary human assembly, was deemed to reduce the sensitivity of variant calling and many other analyses due to most aligners giving MQ of 0 to reads mapped in the flanking sequences. Aligned reads were sorted by genomic coordinates, and read group information, including sample, flowcell, lane, and library, was assigned to each read. Duplicate reads were marked using Sentieon *LocusCollector* and *Dedup*, equivalent to *MarkDuplicates* in Picard (v2.9.0; https://broadinstitute.github.io/picard). To improve mapping accuracy, local realignment around indels was performed using Sentieon *Realigner*, equivalent to *RealignerTargetCreator* and *IndelRealigner* in GATK (v3.8). Base quality score recalibration (BQSR) and base alignment quality (BAQ) recalibration were performed using Sentieon *QualCal*, corresponding to *BaseRecalibrator* and *ApplyBQSR* in GATK (v4.1). Known sites used for BQSR included variants from dbSNP^95^ (v138) and gold-standard indels from Mills and 1000 Genomes Project^96^, as provided in the GATK resource bundle. Original base qualities were retained in the OQ tag, allowing recovery of the initial quality scores from the alignments to enable re-generation of the complete original FASTQs from the final BAM files. To reduce file size, we removed the indel-related base quality score tags BI and BD, as recommended by GATK Best Practices. While these tags may offer refined base quality estimates near indels, they are not currently used by GATK or other standard variant callers in common practice and take a lot of disk space to store.

##### Processing PacBio HiFi WGS data

Data were processed using a standardized pipeline as follows. Reads were aligned to the GRCh38 reference genome (GCA_000001405.15), which excludes ALT contigs and decoy sequences from hs38d1, using pbmm2 (v1.13.0; https://github.com/PacificBiosciences/pbmm2). By default, all tags from the original unaligned reads were annotated to the final alignments. To reduce the file size while retaining relevant alignment information, the --strip flag was used to remove non-essential kinetic tags (dq, dt, ip, iq, mq, pa, pc, pd, pe, pg, pm, pq, pt, pv, pw, px, sf, sq, st). This preserves methylation-related tags MM and ML, and assay-specific tags such as nucleosomes position in Fiber-seq data. Aligned reads were then sorted by genomic coordinates, and read group information, including sample and library, was assigned to each read.

##### Processing Oxford Nanopore Technologies (ONT) WGS data

Data were processed using a standardized pipeline as follows. Reads were aligned to the GRCh38 reference genome (GCA_000001405.15), which excludes ALT contigs and decoy sequences from hs38d1, using minimap2^59^ (v2.26). To ensure consistency with pbmm2, the following flags were applied: -Y to enable soft clipping for supplementary alignments, -L to produce long CIGAR strings (used in the CG tag), --eqx to represent matches and mismatches explicitly using = and X, and --secondary=no to suppress secondary alignments. Aligned reads were then sorted by genomic coordinates, and read group information, including sample and library, was assigned to each read. Methylation-related tags MM and ML were retained from the original unaligned reads and annotated to the final alignments using Methylink (v0.6.0; https://github.com/projectoriented/methylink).

##### Processing RNA-seq data

Data were processed using a pipeline based on the GTEx consortium analysis pipeline. Reads were aligned to the GRCh38 reference genome (GCA_000001405.15), which excludes ALT contigs and decoy sequences from hs38d1, using STAR (v2.7.10b) as implemented in the Sentieon software (v202308.01). The HLA region was also excluded from the reference genome to generate the STAR index files. Genome aligned reads were sorted by genomic coordinates, and read group information, including sample and library, was assigned to each read. Duplicate reads were marked using Sentieon LocusCollector and Dedup, equivalent to MarkDuplicates in Picard (v2.9.0). Transcriptome aligned reads were used for quantification.

To match the GTEx pipeline, quantification and quality control were performed using two independent software packages. RSEM (v1.3.3) was used to generate gene- and transcript-level quantification, while RNA-SeQC (v2.4.2) was used to generate gene- and exon-level quantification as well as quality control metrics. In GTEx, RNA-SeQC results are used for gene- and exon-level read counts and TPM values, and RSEM results are used for transcript quantification. Consistent with this approach, we release both sets of results, allowing users to select gene quantification from either RSEM or RNA-SeQC as needed. GENCODE v47 was used to generate the RSEM index files, and the collapsed gene model for RNA-SeQC.

##### Data quality assessment

Data quality was assessed using a multi-step pipeline at the Data Analysis Center (DAC) at the time of data processing and after alignment for both original raw (FASTQ) and aligned (BAM/CRAM) sequence data to ensure that the data generated by the Network is of acceptable quality for variant analyses The QC status is determined based on the results of select QC checks (**Table S5**), and the QC status of each BAM/CRAM file is indicated in the files of metadata (i.e., “manifest” files), which are visualized in and available to download from the SMaHT Data Portal (https://data.smaht.org). All datasets released from the Network is of either “PASSED” or “FLAGGED” QC status, and datasets with “FAILED” QC status are not released. For Illumina data generated on the NovaSeq platforms, raw reads were evaluated using FastQC (v0.12.0; http://www.bioinformatics.babraham.ac.uk/projects/fastqc) to assess base quality scores, GC content distribution, and the presence of overrepresented sequences including adapter or other known artifacts, such as poly-G runs. For PacBio and ONT data, unaligned reads were evaluated using NanoPlot^97^ (v1.44.1) to calculate read length distributions and other sequencing summary statistics. To assess potential microbial contamination of both bacterial and viral origin, all datasets were also processed using Kraken2^98^ (v2.1.3), applied to the raw reads. To optimize computational efficiency while achieving accurate bacterial detection, the Standard-16 database provided by the authors was used.

Standard quality metrics, including mapping rates and other alignment-based statistics, were calculated using Samtools^99^ (v1.17), Picard (v3.0.0), and a custom in-house Python script (https://github.com/smaht-dac/qc-pipelines/blob/main/dockerfiles/bamstats/bamStats.py). For all sequencing data, alignment-level statistics were summarized using the Samtools *stats* and *flagstat* commands. Sequencing coverage metrics, including mean and per-base coverage, were calculated using mosdepth^100^ (v0.3.9). Additional metrics were generated using Picard tools, including *CollectAlignmentSummaryMetrics*, *CollectBaseDistributionByCycle*, *CollectGcBiasMetrics*, and *MeanQualityByCycle*. For paired-end Illumina data, insert size distributions were calculated using *CollectInsertSizeMetrics*, and overall sequencing performance was evaluated using *CollectWgsMetrics*. These metrics include the mean and standard deviation of insert sizes based on read pair alignment, read pair orientation, the fraction of aligned bases excluded due to low mapping quality or low base quality, and the proportion of bases reaching predefined coverage thresholds. For RNA-seq data, additional metrics were generated using RNA-SeQC (v2.4.2), including the rate of rRNA sequences among mapped reads, 5’-3’ coverage ratios, estimates of library complexity, the exonic–intronic ratio, the number of genes detected, the intergenic rate, and the percentage of chimeric reads. We also implemented an in-house software that predicts tissue types using the RandomForest classification approach, to check correct tissue labeling based on gene-expression profiles.

Sample identity and integrity were evaluated using Somalier^101^ (v0.2.19), which calculates the degree of relatedness between samples based on alignments to detect potential sample swaps or mislabeling. Samples with relatedness scores of 90% or lower were flagged for further review at the Data Analysis Center and the sequencing center to determine whether a swap or mislabels had occurred. To assess contamination from other human samples, VerifyBamID2^102^ (v2.0.1) was used with a panel of known polymorphic loci in GRCh38, based on the 1000 Genomes Project and provided by the authors (https://github.com/Griffan/VerifyBamID/tree/master/resource). To increase computational efficiency while achieving accurate detection for PacBio and ONT data, VerifyBamID2 was run from pileups instead of the alignments directly. Intermediate pileups were generated using Samtools *mpileup* with the following options: -B to disable BAQ computation, -a to output all positions, -s to output base alignment quality, and -q for minimum alignment quality.

Datasets with low to moderate levels of human contamination (<3%) were considered still useful for detecting somatic variants and made available for download at the SMaHT Data Portal (https://data.smaht.org). Datasets identified as having sample swaps or mislabels were retracted or deleted. The Data Analysis Center keeps a record of retracted and replacement datasets in the metadata, and these are made available to view in the Data Portal.

For all output files released by the SMaHT Data Analysis Center, the file format and integrity were checked using MD5 checksum validation and file-format sanity checks developed in-house to confirm EOF and integrity of all generated alignment files. All bioinformatics pipelines at the DAC were executed in the Amazon Web Services (WGS) infrastructure, designed and implemented to support modular, distributed processing at scale.

### Analysis Methods

#### SNV and indel calling

*contributed by Yeongjun Jang, Stephanie Gardiner, Stephanie Georges, and Yoo-Jin Jiny Ha*

Full details of mutation calling and reference-set construction using the COLO829 cell lines are provided in a dedicated manuscript^15^. Briefly, to define a benchmarking set of somatic mutations unique to COLO829, we applied four variant callers—TNHaplotyper2 (Sentieon Genomics 202112.06), Strelka2 (v2.9.2), VarNet (v1.1.0), and RUFUS (v0.1)—using COLO829BL as the matched control. PacBio HiFi support was assessed using bcftools mpileup (v1.17). For SNVs, PacBio allele counts were parsed per candidate site, and calls were retained if supported by ≥2 tumor reads and absent in the matched blood sample. For indels, additional breakpoint-aware annotation was performed to quantify allele balance and distinguish the targeted indel from other alternative alleles with Illumina COLO829BL data. We also established a negative control set to avoid misclassifying true variants as false positives in the benchmark, ensuring a fair evaluation of variant caller performance and preventing penalization for detecting true mutations missed during reference set construction. The negative control set consists of homozygous reference sites in both COLO829T and COLO829BL samples. Germline variant sites were also provided as a separate negative control set.

Somatic mutation calling for the COLO829BLT50 mixture was performed using 12 available pipelines based on DRAGEN, Lancet2, GATK/Mutect2, TNHaplotyper2, RUFUS and MosaicForecast. Although post-calling filters varied across workflows, they typically removed calls in repetitive/duplicated regions and known germline variation, and applied VAF- and coverage-based thresholds and quality filters.

We evaluated performance across three genomic context strata. Easy regions were defined by the 1000 Genomes strict mask (high-confidence, short-read–callable loci). Difficult regions were sites within the PanMask strict (pm151b) accessibility set but outside the 1KG strict mask, and extreme regions were loci outside PanMask, which are typically highly repetitive or structurally complex; PanMask provides a pan-genome–derived, less reference-biased accessibility framework (detail and resources can be found on GitHub https://github.com/parklab/SMaHT_Regional_Categorization).,

To identify cell culture-induced mutations that may have arisen during continuous expansion and harvesting of COLO829BL lymphoblastic cells, we examined variants present in COLO829BLT50 but absent from the COLO829 reference set. RUFUS and Mutect2 were each run in paired control mode on the five COLO829BLT50 Illumina WGS datasets—generated across different sequencing centers—using COLO829BL as the matched normal. The RUFUS and Mutect2 call sets were merged using bcftools isec (v1.23), to generate five sequencing center-specific COLO829BLT50 call sets. The COLO829 reference set was subtracted from these sequencing center-specific sets, to retain the set of candidate cell line artifacts. We hypothesized that mutations identified across all five sequencing centers (set C(5,5)) were more likely to be true variants than those found in four (C(5,4)), three (C(5,3)), or fewer center-specific data sets, which are more likely to be sequencing or mutation calling tool artifacts.

We have used available, deep-coverage PacBio HiFi long-read data as correlative evidence to determine if any given candidate is truly a culture-induced mutation or an artifact. We called a candidate a true culture-induced mutation if the following criteria were fulfilled: (1) the presence of ≥2 supporting PacBio reads in COLO829BLT50 (confirming the mutation in the cell-line mix); (2) no PacBio or Illumina supporting reads in COLO829BL (confirming the absence of the mutation in the original blood cell line); and (3) no PacBio or Illumina supporting reads in COLO892 (confirming the absence of the mutation in the tumor cell line). Next, we checked the presence of the mutation in a deeply sequenced sample from an unrelated individual, donor ST001, and accepted the site if the mutant allele was present < 2 PacBio reads (eliminating recurrent sequencing error at the candidate mutation site). Finally, we removed any variants defined in the negative control set (described above). This procedure yielded a total of 5,372 mutations across C(5,5), …, C(5,1) sets; as expected C(5,5) contributing the largest fraction of sites.

We carried out a similar procedure for INDEL mutations. Here, the fraction of sites confirmed by the filtering procedure described above was so low in C(5,3), C(5,2), and C(5,1) that we only kept confirmed sites from C(5,5) and C(5,4). The procedure yielded a total of 196 INDEL sites.

Full details of mutation calling in fibroblasts and post-mortem tissues are provided in a dedicated manuscript^16^. To identify benchmarking somatic SNVs and indels unique to fibroblasts we performed an all-to-all exhaustive comparison of genomes of 25 clonal iPSC lines. These initial calls were then processed using the All2 filtering strategy^45^ and calls with VAF > 25% were retained. We designated the mutations discovered in five LA2 iPSC lines as the benchmarking set. Mutation from select iPSC lines that originated from different biopsies formed the negative control mutation set.

Somatic mosaic mutation calling in the fibroblast sample was conducted using RUFUS tools and Mutect2 (GATK version 4.2.6) in three different modes to ensure comprehensive discovery: (1) Tumor with Matched Normal, (2) Tumor-Only with default parameters, and (3) Tumor-Only with high sensitivity options “--initial-tumor-lod 0 --tumor-lod-to-emit 0 --af-of-alleles-not-in-resource 5e-8 --adaptive-pruning-initial-error-rate 0.0001 --pruning-lod-threshold -4”. In the Tumor with Matched Normal mode, the five iPSC lines comprising the negative control set were used collectively as the matched “normal” samples against which the fibroblast was compared. For the Tumor-Only mode, we used a Panel of Normals (PON), which we downloaded from the GATK resource bundle. Calls were filtered using a slightly modified version of the BSMN filtering workflow^11,44^. Called mutations were confirmed using data from different sequencing platforms, specifically PacBio and AVITI. A minimum quality cutoff of 20 was applied for both mapping quality and base quality. A mutation call was considered confirmed in fibroblast sample if at least one read supporting the alternative allele met these quality criteria. When analyzing post-mortem samples from the ST002 sample the same workflow as for the fibroblasts was utilized but for additional stringency, we applied two additional filters to generate the final set of mutations: MFRL (median fragment length) >300 and MMQ (median mapping quality) >56.

#### Methods for MEI analyses

*contributed by Eunjung Alice Lee and Weichen Zhou*

Full details of MEI calling analysis can be found in the MEI benchmarking manuscript^21^.

We first established a benchmarking set in the HapMap mixture experiment. Computationally, germline MEIs were defined per HapMap donor by combining assembly-based callsets (i.e., DipCall v0.20 and Minigraph v0.34 on haplotype-resolved assemblies) with read-based calls (i.e., Sniffles2 v2.2.2). Candidate insertions were then systematically annotated and filtered to focus on young, active subfamilies (L1HS/L1PA2, AluY/AluYb8/AluYa5, SVA_E/F). A custom TPRT-signature pipeline assigned mechanistic evidence by detecting TSDs (via synthetic pseudo-reads and bwa-mem CIGAR inspection) and polyA/T tails, enabling confidence tiering (Tier 1: TSD+polyA/T; Tier 2: polyA-only; Tier 3: remaining young-subfamily insertions). Zygosity was inferred from VAF distributions using a Gaussian mixture model. Somatic truth in the mixture was simulated by labeling HG005-present insertions as “germline” and insertions private to minor donors as “somatic,” with expected VAF computed as mixture ratio × zygosity.

We then conducted HapMap mixture benchmarks for computational sMEI detection and evaluation, emphasizing read-signal-driven callers and harmonized scoring. Single-sample-capable WGS methods (short-read: xTea_mosaic v0.1.9, RetroSom v2, MELT v2.2.2; long-read: PALMER v2.0.1; plus tuned germline SV callers xTea_long v0.1.0, cuteSV v2.1.2, Sniffles2 v2.6.3 with reduced support thresholds) were run with repeat-based annotation (RepeatMasker) and young-subfamily filters to prioritize biologically plausible events. Performance was quantified against the tiered truth sets using breakpoint matching (≤100 bp for WGS; ≤200 bp for targeted) to define TP/FP, with recall/precision/F1 computed globally and stratified by mappability (151-mer unique vs non-unique), repeat context (ME vs ME-free), expected/caller VAF bins, and insertion length bins. Two computational refinements were layered on top: (i) cross-platform integration combining callset overlap and raw clipped-read evidence between short- and long-read data, with caller-specific FP filters (e.g., exclude non-unique regions for SR-only; require young-subfamily BLAST support and remove supplementary-only evidence for LR-only), and (ii) local haplotype phasing around each insertion (high-quality hetSNPs within ±40 kb; long reads spanning SNP+insertion) to separate likely germline from low-read, haplotype-linked mosaic events. Finally, the impact of DSA alignment on false positives in segmental duplications was tested using a binomial Generalized linear model comparing FP composition in SD vs outside SD while adjusting for caller and mixture effects.

For benchmarking donors (tissue homogenates), we called sMEIs per sample using xTea_mosaic on Illumina WGS and PALMER on long-read WGS, then applied additional computational filters (BLAST-based subfamily annotation with removal of older subfamilies and supplementary-alignment-only calls). Germline candidates were reduced by haplotype phasing, cross-sample consistency checks (sites classified as germline in any sample were treated as germline across samples), and exclusion of known polymorphic MEI loci curated from prior studies. We then applied the same short-/long-read integration framework across all platform pairs per donor, followed by structured manual review using TPRT evidence (TSD/polyA), breakpoint concordance, and repeat identity to finalize high-confidence sMEIs and estimate VAF from supporting reads normalized by local long-read depth.

#### Methods for SV analyses

*contributed by Fritz Sedlazeck*

Human Pangenome Reference Consortium assemblies for the six HapMap samples were procured from https://github.com/human-pangenomics/HPP_Year1_Assemblies. Each of the twelve haplotypes were aligned to GRCh38 and variants called with minimap and paftools, and post processed with BCFtools and Truvari to consolidate SV across haplotypes and collapse highly similar, distinct SVs.

Eight callers that met the following inclusion criteria were tested: (1) the default mode supports calling SV at VAF <10%; (2) the caller accepts single-sample input (i.e., requires no paired matched control). We tested GRIDSS2 (v2.13.2), SvABA (v1.2.0) and Delly (v1.2.6) for short read data; Severus (v0.1.1), Sniffles (v2.4), and Delly (v1.2.6) for long read data; Sawfish (v0.12.4), pbsv (v2.9.0), and SVDSS (v2.0.0-alpha.1) for HiFi data only. All callers were executed with default parameters. Uniform VCF normalization was conducted for each call set, and we only evaluated duplication, insertions and deletions (over 50 bp in length) with PASS flag.

Truvari was used to benchmark each call set against the set of benchmark SVs. Calls were considered as germline if present in HG005; somatic calls were defined as those with the reported <25% VAF. Somatic SVs from the benchmark were partitioned into six VAF bins: <0.5%, 0.5-1%, 1-4%, 4-5%, 5-10%, and 10-20%. In the difficulty-stratified analysis, each benchmark and caller-reported SV was annotated with the following features: proximity to other SV; breakpoint overlapping a tandem repeat; low VAF; small event; and insertions. Calls with none of the features were considered as “not-challenging”. To compare platforms at matched depth, replicate BAMs were merged per platform and then randomly down-sampled to 60X, 90X, 120X, 150X, and 180X.

#### Methods for assembly

*contributed by Min-Hwan Sohn, Youngjun Kwon and Andrew B. Stergachis*

UW-SCRI Genome Characterization Center constructed the COLO829BL diploid DSA^19^ using COLO829BL-derived PacBio HiFi/Fiber-seq (60×) and the combination of ONT Ultra-long WGS and ONT Standard WGS (60x) coupled with Hi-C sequencing data (30×) with Verkko v2.1^36^. To maximize N50 of the assembly, ONT reads were downsampled by selecting the longest reads until a targeted 60x coverage was achieved. Specifically, based on the index file (.fai) of the ONT FASTQ, read IDs were extracted in descending order of read length until reaching 60× coverage based on 3.1 Gbp as the genome size. Reads corresponding to these IDs were then stored separately to generate the downsampled ONT FASTQ file. Phased assembly was generated using Verkko v2.1^36^. For the benchmarking donors (ST001, ST002, ST003 and ST004) and LB-LA2 Fibroblast DSA, HiFi-only diploid assemblies were generated using hifiasm^58^ with default parameters, including the --dual-scaf and --telo-m CCCTAA options. PacBio HiFi reads from available tissue data for ST001 and ST002 were downsampled to 60× coverage by selecting the longest reads, consistent with the approach described above, while ST003 (49.76×) and ST004 (37.70×) were assembled without downsampling. SMHTLBLA2 was assembled using 110× coverage.

After the initial assembly, we implemented additional filtering steps to ensure that contaminated sequences are removed using NCBI FCS41 v0.4.0 beta and BLAST42 v2.15. In short, contigs shorter than 10 bp, which can cause errors in FCS, were excluded, followed by the detection of foreign DNA and adapter sequences. Subsequently, mitochondrial, Epstein–Barr virus (EBV) and ribosomal DNA (rDNA) sequences were identified using a BLAST-based approach. The detected contaminant and adapter regions were trimmed, and rDNA contigs were separated from the assembly. The command used to detect foreign DNA is as follows: python3 fcs.py --image FCS_IMG screen genome --fasta HAPLOID_ASSEMBLY.fasta --out-dir OUTDIR --gx-db GXDB_LOC --tax-id 9606. The command used to detect adapter sequence is as follows: av_screen_x -o OUTDIR --euk HAPLOID_ASSEMBLY.fasta. The command used to detect mitochondrial, EBV, and rDNA sequences is as follows: blastn -query HAPLOID_ASSEMBLY.fasta -db TARGET_DB -outfmt 6 –evalue 1e-30 > OUTPUT. Finally, for each haplotype, a decontaminated assembly FASTA file was produced with foreign DNA (including EBV), adapter, mitochondrial sequences, and rDNA contigs removed.

#### Methods for duplex

*contributed by Yang Zhang, Vinayak V. Viswanadham, and Michail Andreopoulos*

Duplex sequencing data were processed using platform-specific analysis pipelines optimized for each library architecture and benchmarked against low-mutation reference samples. Across all methods, reads were aligned to the hg38/GRCh38 reference genome without decoy sequences when applicable, duplex molecules were reconstructed according to platform-specific designs, and somatic variants were retained only after stringent filtering to suppress sequencing, amplification, and single-strand artefacts. Technology-specific thresholds and post-processing steps were calibrated separately for each platform rather than imposed uniformly across assays. Detailed implementation, parameter settings, and benchmarking procedures are described in the Duplex sequencing companion manuscript^22^.

For ppmSeq, aligned CRAMs were processed using the manufacturer’s preprocessing workflow together with the downstream SRSNV framework and an additional custom SNV-calling procedure that estimates sensitivity-corrected mutational spectra and mutation burden under a predefined false discovery rate. Analyses were restricted to the Ultima high-confidence genome regions.

For CODEC, data were processed from raw sequencing output to variant calls using the standard CODECsuite workflow implemented in a Terra/WDL-based cloud pipeline.

For CompDuplex-seq and NanoSeq, variant calling and filtering were performed using a unified CompDuplex-NanoSeq workflow that includes read preprocessing, alignment, consensus molecule generation, duplex validation, and stringent filtering based on sequence quality, mapping quality, and duplex support.

For HiDEF-seq, data were processed using the HiDEF-seq v3.1 workflow, which is compatible with PacBio Revio data and preserves the core read-processing and filtering logic of the previously published HiDEF-seq pipeline (https://github.com/evronylab/HiDEF-seq) while using a refactored Nextflow-based implementation. For VISTA-seq, somatic SNVs were called using a customized workflow derived from pre-pe and lianti, with modifications for barcode extraction, pooled read handling, and additional filtering to reduce recurrent false positives in multi-allelic and pooled-input settings.

#### Methods for PTA analyses

*contributed by Joe Luquette and Milovan Suvakov*

Full details of computational analyses of PTA data are provided in a dedicated manuscript^23^. Small somatic mutation detection and analysis with SCAN2 was performed to call single nucleotide variants (SNVs), insertions and deletions <50 bp (indels), and dinucleotide variants (DNVs). SCAN2 is a somatic mutation caller specifically developed for analysis of PTA-amplified single cells. The pipeline utilized the GRCh38 reference genome, dbSNP v151 for common variant annotation, and Sentieon’s implementation of GATK HaplotypeCaller. To enable sensitive somatic indel calling, a cross-sample panel was constructed from 100 kb bins spanning chromosomes 1-22, X and Y (excluding pseudo-autosomal regions and regions with anomalously high sequencing depth), using single cells and matched bulks from this study combined with previously published PTA single-cell and matched bulk data downloaded from dbGaP. Mutations were called separately for colon and lung tissues to allow single cells to be matched with bulks from the opposite tissue, thereby reducing the likelihood that SCAN2 would incorrectly discard clonal mutations due to read support in the matched bulk. To retain maximum read support during variant calling, all samples were analyzed simultaneously rather than employing a standard GVCF approach, which loses information by omitting low read support. Eagle v2.4.1 was used for phasing with the 1000 Genomes reference panel. Mutation rates were automatically estimated by SCAN2, and false discovery rates were estimated using previously established false positive rates for PTA-amplified genomes. Sequencing data quality control evaluated cells across five metrics generated by SCAN2: MAPD, allele balance (distribution of VAFs at heterozygous germline SNPs), read depth distribution, GC bias (assessed by fitting a loess curve to read counts in non-overlapping 1 kb bins as a function of GC content), and genomic read depth profiles. Cells exhibiting outlier status in two or more of these metrics were excluded from all small mutation analyses (SNVs, indels, and DNVs).

Single-cell lineage reconstruction was performed using Sequoia. To avoid discarding early somatic mutations that define cell phylogenies near the zygote, SCAN2’s filters requiring no read evidence of a somatic mutation in the matched bulk sample were relaxed by modifying the maximum bulk alternate read count, allele frequency, and binomial probability parameters. Relaxed calls were extracted and a list of unique mutations, identified by chromosome, position, reference and mutated sequence, was compiled across cells. The numbers of reference- and mutation-supporting reads were retrieved for each cell and site to create site-by-cell matrices of reference and mutation read counts. These matrices formed the input for Sequoia, which further filtered the relaxed mutation calls and inferred the phylogenetic tree. To account for artificial overdispersion introduced by research site-specific variation, beta-binomial overdispersion-based filtering was expanded to calculate the maximum likelihood overdispersion parameter (rho) across all cells and within cells from each center, requiring rho > 0.3 for shared mutation calls. For comparison, somatic mutations independently identified in high-depth, multi-tissue bulk whole-genome sequencing were interrogated in the single cells by generating per-cell pileups, producing variant-by-cell read count matrices that were similarly processed by Sequoia. Topologies of the bulk-and single-cell-derived phylogenetic trees were compared using a normalized generalized Robinson-Foulds distance computed via the JaccardRobinsonFoulds metric from the R package TreeDist.

Mutational signature extraction was conducted using the HDP (hierarchical Dirichlet process) R package. Counts spanning 257 distinct categories were analyzed: 96 SNVs with trinucleotide contexts, 83 indels classified by length and repeat/microhomology characteristics, and 78 distinct types of DNVs. To prevent double counting of mutations, variants were assigned to individual phylogenetic tree branches as dependent nodes, with sequencing centers constituting the parent nodes in the hierarchical structure.

Genome-wide copy number alteration (CNA) detection was conducted using CNVpytor and HiScanner. CNVpytor utilized a two-dimensional approach integrating both read depth coverage and B-allele frequency (BAF) signals during segmentation, with germline SNPs phased using shapeit4. BAF was calculated at heterozygous SNPs within the 1000 Genomes Project strict mask in 100 kb bins. High-confidence events were identified by filtering for integer copy number changes expected in a single cell: deletions (CN < 1.5, |delta_BAF| > 0.4), duplications (CN > 2.5, |delta_BAF| 0.08 to 0.24), and copy-neutral loss of heterozygosity (CN-LOH; 1.6 < CN < 2.4, |delta_BAF| > 0.4), with all events required to span at least 10 bins of heterozygous SNPs and a minimum size of 10 Mb. HiScanner integrated read depth with BAF and phasing information from Eagle-phased germline SNPs created by SCAN2. Autosomal diploid segments were examined for CN-LOH by segmenting into approximately 5 Mb intervals and identifying those with mean BAF < 0.25. To smooth out small CNAs potentially representing noise, segments spanning 10 Mb or less had their copy number set to that of the nearest larger flanking segment, and adjacent segments with equal copy number were merged. HiScanner was also executed in read depth-only mode to quantify sex chromosome aneuploidies. Aneuploidies on the Y chromosome were further analyzed using majority-voting phasing of pseudoautosomal region (PAR1) SNPs from cells with chromosome Y loss, which allowed high-confidence definition of the Y-chromosome haplotype.

Somatic structural variants (SVs) were detected by running Manta with default parameters, comparing each single-cell sample against the matched bulk sample. Manta’s deletion and duplication calls larger than 100 kb were treated as candidates and subsequently validated using CNVpytor by requiring a corresponding integer-level copy number change and significant BAF shift. Of the confident calls, the majority were located in T-cell receptor loci, consistent with V(D)J recombination. For complex rearrangements, Manta’s break-end (BND) calls were used to provide orthogonal, breakpoint-level support for large sub-chromosomal CNAs identified by CNVpytor.

#### Comparing benchmarking mutations set for SVs and MEIs

*contributed by Nahyun Kong and Zitian Tang*

To evaluate the agreement of two complementary somatic benchmarking strategies in HapMap cell line mixtures, we compared SV and MEI benchmark sets generated using pangenome-based and hybrid assembly-based approaches. This comparison was designed to determine the extent to which the two methods produce concordant results, identify variant sets unique to each approach, and clarify the factors underlying these differences. We focused specifically on SVs and MEIs because these variant classes are often underrepresented in published somatic benchmarking resources.

Prior to comparison, shared benchmark regions were defined by intersecting the pangenome-based^17^ and hybrid assembly-based^20,21^ benchmark regions using bedtools (v2.27.1) intersect, and all subsequent variant comparisons were restricted to these shared regions. Since the pangenome-based approach targets pseudo-somatic variants exclusively, with HG005 germline variants filtered out, non-HG005 SVs and non-HG005 MEIs were selected from the assembly-based benchmark set or MEI working group set before comparison. For SVs, the SV working group easy set and the pangenome-based SV benchmark set were evaluated using Truvari v4.0.0 with the -pick multi flag. For MEIs, concordance between the MEI working group’s tier1 benchmark set and the pangenome-based MEI benchmark set was assessed by comparing insertion start positions within the shared benchmark regions using in-house scripts.

When comparing SV and MEI benchmark sets generated using a pangenome-based approach with those from hybrid, assembly-based approaches, we found substantial overlap. The primary differences laid in variant representation: the pangenome-based method preserved exact genotypes and haplotype structures, separating overlapping variants into distinct records, whereas the assembly-based approaches merged comparable variants to facilitate benchmarking. Each approach therefore offers distinct advantages, depending on the goals of benchmarking and downstream analyses. For MEI, counts were also different depending on whether they were classified based on repeat annotation (higher count) or mechanistic signature of target primed reverse transcription (lower counts).

#### Evaluating pangenome-based somatic variant detection approach

*contributed by Qichen Fu and Benpeng Miao*

Full details of pangenome-based somatic variant detection and benchmark can be found in the pangenome somatic variant benchmark manuscript^18^.

The assemblies from HPRC release 1, together with GRCh38 and CHM13^33^, were used to construct pangenome graphs using Minigraph-Cactus^62^. This generated both allele-frequency (AF)-filtered and unfiltered pangenome graphs. In this study, only the clipped graphs were used.

The pangenome-inferred diploid assembly was generated from the AF-unfiltered pangenome graph. First, HG005 whole-genome sequencing (WGS) data were used to generate k-mer profiles with KMC using the options -k29 -m128 -okff^103^. Haplotype sampling was then performed by matching k-mer profiles derived from the WGS data to haplotypes in the pangenome graph using vg (v1.64.1) haplotypes^104^. This procedure imputes a subgraph from the clipped, unfiltered HPRC Release 1 pangenome graph^104^, using GRCh38 as the reference backbone and incorporating two recombinant assemblies (hap1 and hap2). These recombinant assemblies closely approximate the HG005 diploid assembly^18^ and together constitute the pangenome-inferred diploid assembly used in this study.

To generate HapMap mixture call sets across GRCh38, the pangenome, and the pangenome-inferred diploid assemblies, we used 500× Illumina short-read WGS data from the WashU-VAI GCC. Reads were aligned to GRCh38 using BWA-MEM (v0.7.17)^105^. The same dataset was also aligned either to the clipped, AF-filtered pangenome graph or to personalized pangenome graphs containing GRCh38 and the pangenome-inferred diploid assembly using vg giraffe (v1.64.1)^106^. Alignments were subsequently surjected to GRCh38 for the AF-filtered pangenome graph, or to the pangenome-inferred diploid assemblies for the personalized pangenome graph, using vg surject^33^. All alignments were processed with GATK (v4.1.0.0)^90^, including duplicate marking, indel realignment, and base quality score recalibration (BQSR) before variant calling.

SNVs and indels were called from 500× short-read data using MuTect2 (v4.5.0.0)^50^ in tumor-only mode. Variants were subsequently filtered with FilterMutectCalls, and only calls annotated as “PASS”, “haplotype” or “clustered_events” were retained. Variants were then decomposed and normalized with bcftools norm (v1.20-24-g5d48dbc3)^99^ using the options -a --atom-overlaps ‘.’ --check-ref w -N, and finally separated into SNVs and indels using Picard SplitVcfs (v2.9.0).

For benchmarking, we used HapMap mixture benchmarking variant sets based on GRCh38 and the pangenome-inferred diploid assembly generated using a genome-graph approach^17^ (https://zenodo.org/records/17544617). The benchmarking was performed within the confident regions on chromosomes 1-22, X, and Y. GRCh38 confident regions were further stratified into easy, difficult, and extreme regions as defined by the SMaHT network (https://data.smaht.org). These stratifications were lifted over to pangenome-inferred diploid assembly using UCSC liftover and the corresponding chain files generated by nf-LO^107^.

The somatic SNVs were benchmarked using RTG vcfeval (v3.12.1)^108^ using options --squash-ploidy and --all-records. SNVs were partitioned into nine variant allele fraction (VAF) bins: 0-0.5%, 0.5-1%, 1-1.5%, 1.5-2%, 2-3%, 3-4%, 4-5%, 5-10%, and 10-16.5%. For recall assessment, the benchmarking variant set was stratified by VAF, and each VAF bin was compared against the full call set. For precision assessment, the call set was stratified by VAF, and each bin was compared against the complete benchmarking variant set. Precision and recall were calculated as follows: Precision = TP / (TP + FP); Recall = TP / (TP + FN).

#### Evaluating DSA-based somatic variant detection approach

*contributed by Min-Hwan Sohn and Andrew B. Stergachis*

We established a comprehensive benchmarking framework using *in silico* tumor-normal mixtures at three dilution ratios (1:4, 1:9, and 1:19 tumor:normal) corresponding to approximately 20%, 10%, and 5% variant allele fractions. These dilution ratios were designed to simulate the detection of low-fraction mosaic somatic variants in donor tissues within a variable range that depends on the timing of mutation acquisition during development and the proportion of affected cells in the sampled tissues. *In silico* mixtures were generated by independently subsampling pure normal (COLO829BL) and tumor (COLO829) Fiber-seq data aligned onto COLO829BL DSA (samtools *view* --subsample) to achieve target coverage ratios and then merging the subsampled files to create composite alignment datasets at 20× total haplotype coverage. For comparative analysis against GRCh38, the aforementioned DSA-mapped alignment mixtures were first converted to unaligned BAM format, then realigned to the GRCh38 reference genome, maintaining identical read composition but with different genomic assembly coordinates.

The method for obtaining a high-confidence DSA-based somatic SNV reference set for COLO829 is described in detail in Sohn et al.^19^. Briefly, we performed SNV calling using DeepVariant v1.6.1^64^ on COLO829BL Fiber-seq and two COLO829 Fiber-seq biological replicates independently. Somatic variants were identified by retaining only those variants present in both COLO829 replicates but absent from the COLO829BL with additional quality control and masking steps to ensure high quality variant set. This involves masking genomic positions containing variants that are either unique to single tumor replicates or present only in the normal sample from evaluation to prevent mis-categorization of variants into confusion matrix; the former representing potentially true but somatic events with lack of strong evidence, and the latter representing potential culture artifacts or assembly artifacts since these variants should not appear as somatic relative to an assembly derived from the same source. For GRCh38-based evaluation, DSA-based somatic SNV coordinates were moved to GRCh38 coordinates using a donor-specific graph (DSG)^19^ constructed by minigraph-cactus^62^ from Telomere-to-Telomere assembly of the CHM13 cell line^109^ (T2T-CHM13v2.0), GRCh38 and COLO829BL DSA haplotypes 1 and 2, with SNV positions from COLO829BL DSA space injected into the graph and surjected onto GRCh38 using variation graph (vg) toolkit^63^ to generate the corresponding reference set. The same approach was applied to generate a T2T-CHM13-based reference set by surjecting the injected SNV positions onto T2T-CHM13v2.0 coordinates. Genomic loci unique to the COLO829BL DSA with respect to each reference were defined in a similar fashion using DSG, retaining only regions that could not be surjected to GRCh38 or T2T-CHM13v2.0, respectively, and are ≥1kb in length.

Finally, we ran DeepSomatic v1.9.0^40^ in tumor-only mode at different dilution ratios on DSA, GRCh38 and T2T-CHM13, respectively. Somatic SNVs were further filtered to include only PASS variants and exclude variants in masked genomic position. Furthermore, we restricted the evaluation only to pre-computed callable genomic regions in the DSA defined by accounting for the misassembled proportion of the genome and also large-scale deletions. GRCh38-based variants and T2T-CHM13-based variants were also filtered against a panel of normals (PoN) catalogues packaged within DeepSomatic to reduce germline contamination^40^. The PoN records in GRCh38 coordinates were lifted over to T2T-CHM13v2.0 using the Picard LiftoverVcf (v.2.23.3) with default parameter following the exact method described in the link: https://s3-us-west-2.amazonaws.com/human-pangenomics/T2T/CHM13/assemblies/annotation/liftover/dbSNP.html.

## KEY RESOURCES TABLE

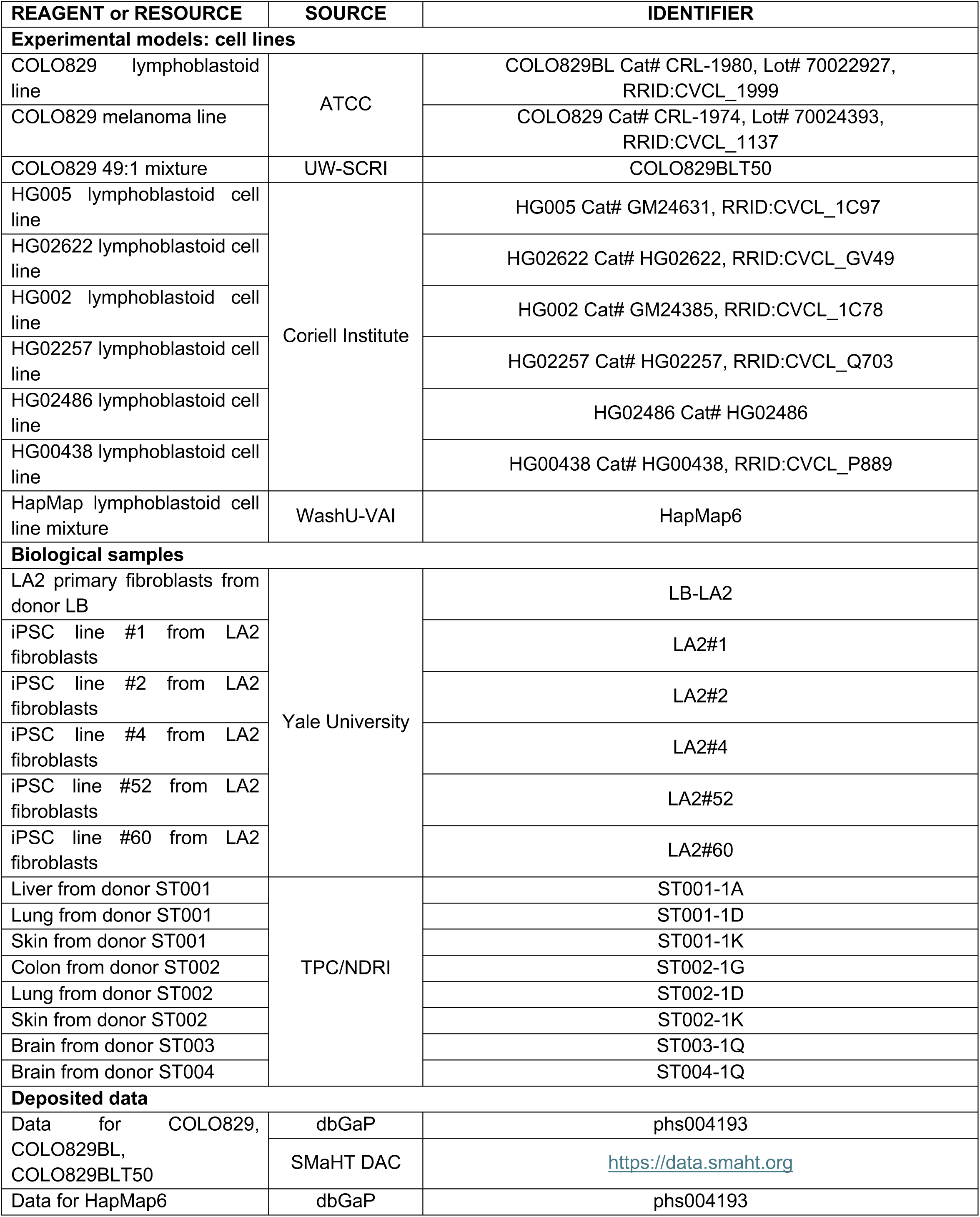
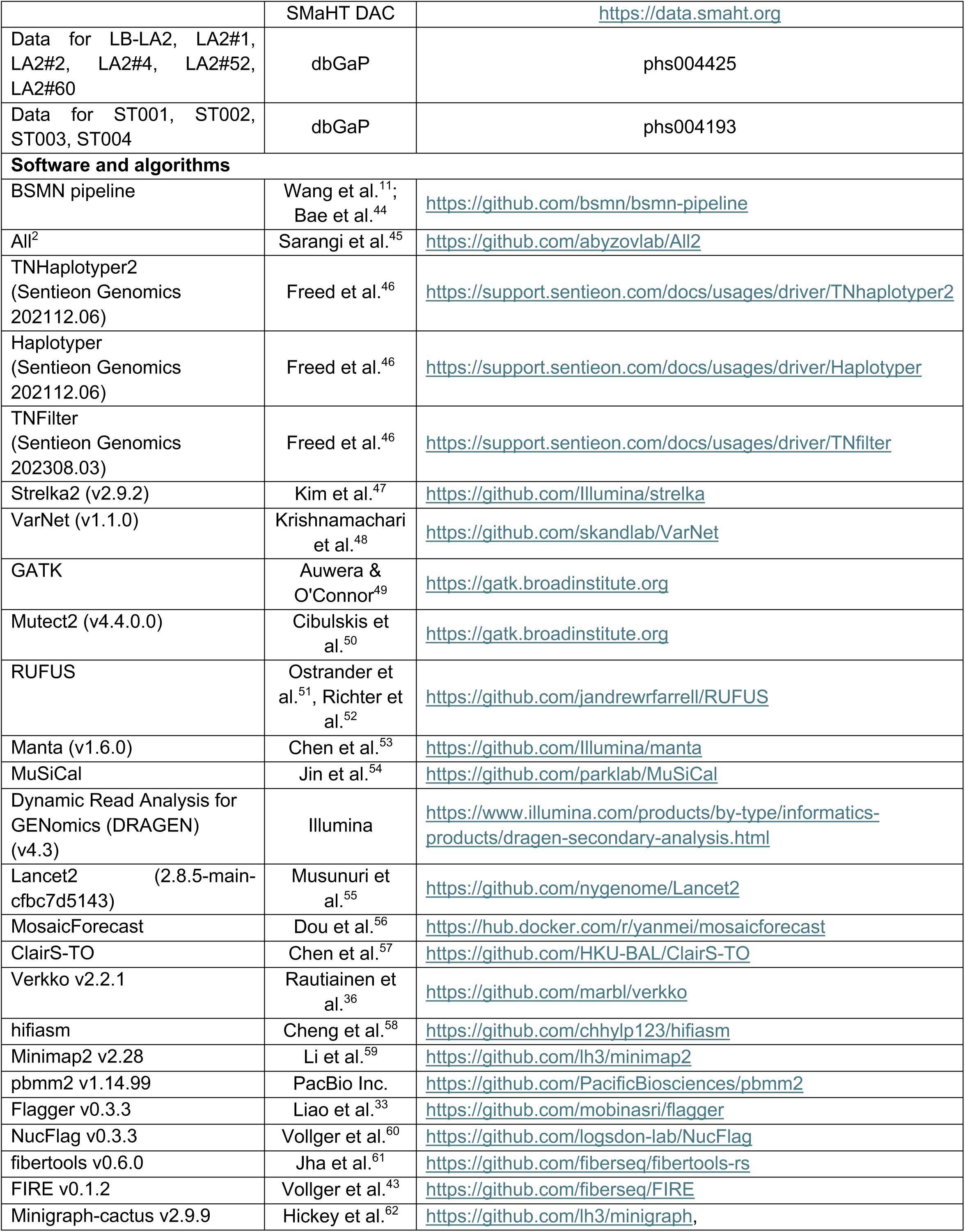
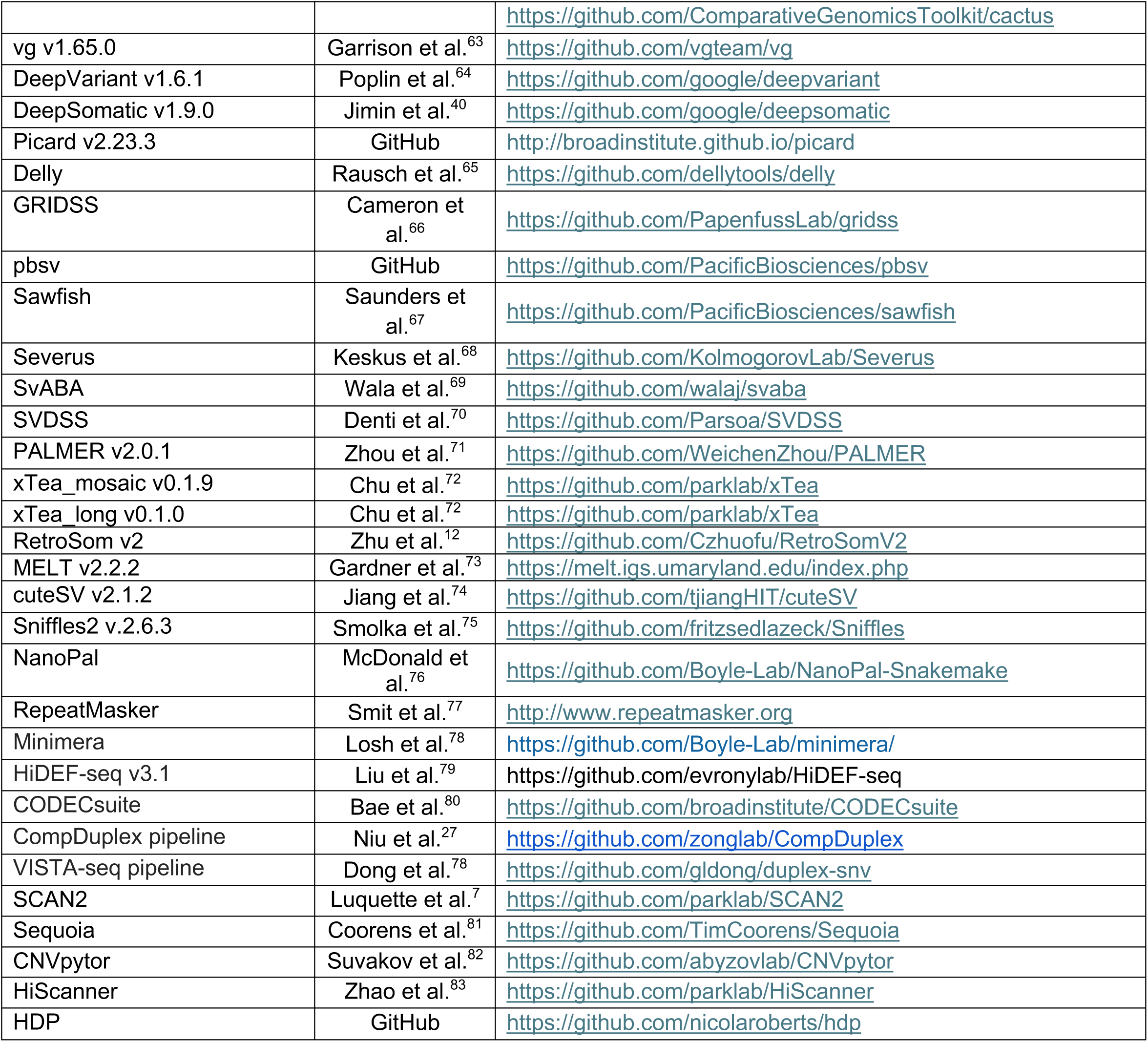

