## Supplementary figures and tables for "Comprehensive benchmarking of somatic mutation detection by the SMaHT Network"

### Supplemental information

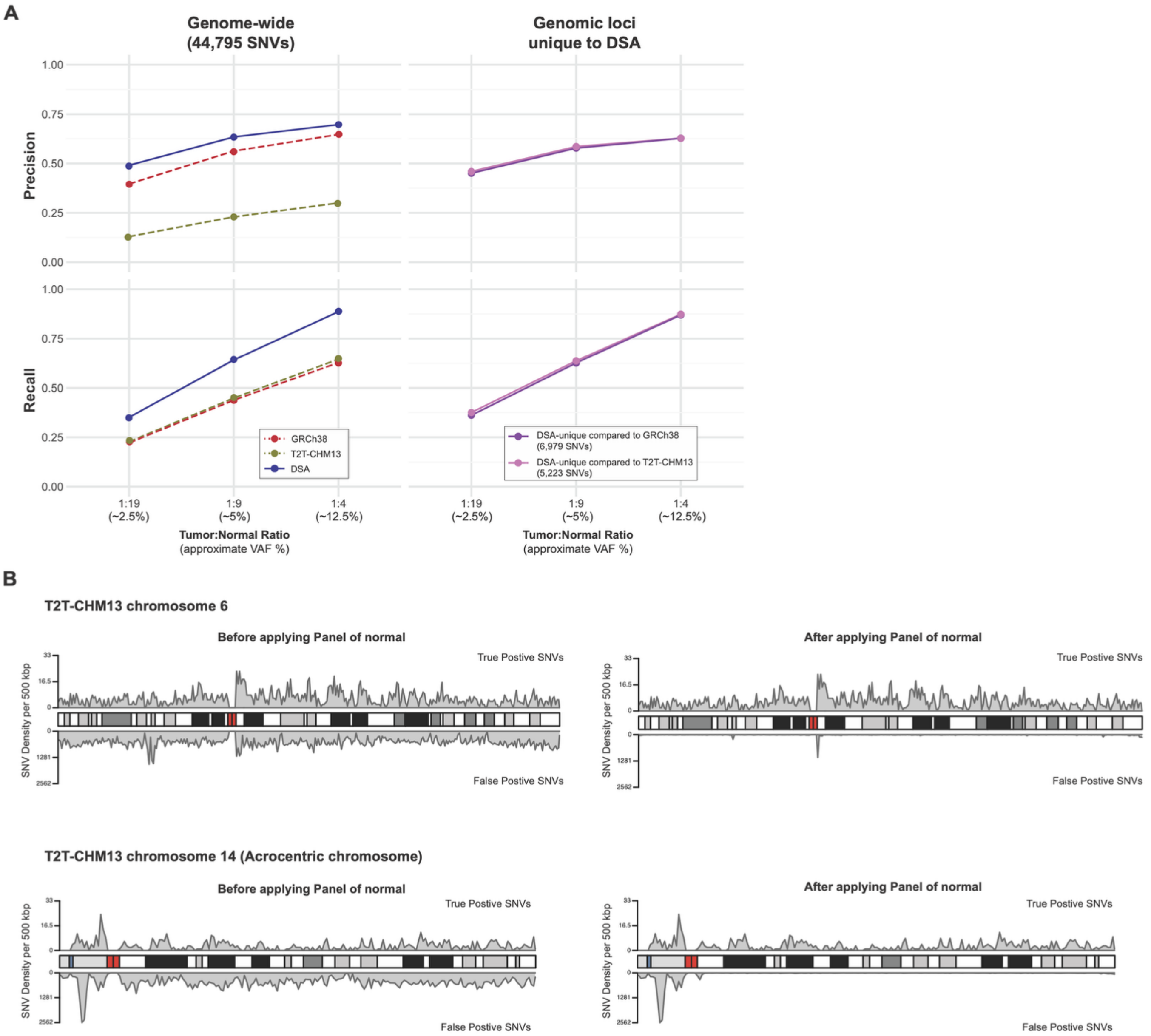


**Figure S1. Benchmarking different assemblies (DSA, GRCh38 and T2T-CHM13) for somatic SNV discovery. A)** Precision and recall for somatic SNV discovery using 40× PacBio HiFi long-read sequencing data of in silico mixture of the COLO829BL and COLO829 cell lines for GRCh38-, T2T-CHM13- and DSA-based approaches. Findings are separated into genome-wide, as well as genomic loci that are unique to the COLO829BL DSA compared to GRCh38 and T2T-CHM13, respectively. **B)** Ideogram examples showing the effect of a panel of normals (PON) on the somatic SNVs discovered using the T2T-CHM13 reference genome with false positives shown on the bottom of each panel. Most of the false positive somatic SNVs are eliminated by PON filtering, with the exception of regions not well represented in germline databases catalogues based on GRCh38, such as those surrounding centromeres or the short arms of acrocentric chromosomes.

**Table S1. Benchmark Data Generators.** The following institutions generated the SMaHT benchmark sequencing data and metadata that were submitted to and analyzed at the Data Analysis Center.

| **Institution (PI)** | **Abbreviation used in this study** | **Role in the SMaHT Network** |
|---|---|---|
| Baylor College of Medicine (Gibbs) | BCM | GCC (Genome Characterization Center) |
| Broad Institute (Ardlie) | Broad | GCC |
| New York Genome Center (Germer) | NYGC | GCC |
| University of Washington and Seatle Children Research Institute (Bennett) | UW-SCRI | GCC |
| Washington University at St. Louis and Van Andell Institute (Wang) | WashU-VAI | GCC |
| Boston Children's Hospital (Walsh) | BCH1 | TTD (Technology and Tool Development) |
| Boston Children's Hospital (Choudhury) | BCH2 | TTD |
| Baylor College of Medicine (Zong) | BCM1 | TTD |
| Dana Farber Cancer Institute and Boston Children’s Hospital (Burns) | DFCI-BCH | TTD |
| Mayo Clinic and Yonsei University (Abyzov) | Mayo | TTD |
| Stanford University and Yale University (Urban) | Yale | TTD |
| New York University (Evrony) | NYU | TTD |
| University of Massachusetts Medical School (Fazzio) | UMass | TTD |
| University of Michigan (Mills) | UMich | TTD |
| Weill Cornell Medical College (Landau) | Cornell | TTD |
| Harvard Medical School (Park) | DAC | DAC (Data Analysis Center) |
| National Disease Research Interchange (Bell) | TPC | TPC (Tissue Procurement Center) |

**Table S2. Comparison of biological and technical advantages (+) and disadvantages (-) of reference cell systems used for mutation benchmarking.**

| | Feature | HapMap lines | COLO829 | Fibroblasts / iPSC |
|---|---|---|---|---|
| **Scientific** | Mutation types | Broad spectrum across all variant classes (+) | High burden of SNVs, indels, SV/CNVs (+); low burden of MEI (-) | Medium burden (+); rare SVs and MEIs (-) |
| | Mutation origin | Germline variants (-) | Tumor-derived somatic mutations (-) | Physiological somatic mutations (+) |
| | Haplotype structure | Multiple unrelated haplotypes (-) | Matched diploid haplotypes (+) | Matched diploid haplotypes (+) |
| | Control of mutation frequencies | Defined by line proportions (+) | Defined by line proportions (+) | Not experimentally controlled (-) |
| **Technical** | Cell growth | Immortal, fast-growing (+) | Immortal, fast-growing (+) | Finite lifespan (-) |
| | Reference assembly | Complete T2T diploid assembly available (+) | Needed to be done (-) | Needed to be done (-) |
| | Existing sequencing data | Extensively characterized with short- and long-read data (+) | Extensively characterized with short- and long-read data (+) | Requires long-read characterization (-) |
| | Sample composition | Requires mixing of multiple lines (-) | Requires mixing for three times (-) | Natural mix of clones (+) |

**Table S3. Overview SMaHT Benchmarking Data generated by Centers (additional file).**

**Table S4. Benchmarking WGS and RNA-Seq Data Overview (additional file).**

**Table S5. Various QC checks performed on whole genome sequencing data at the SMaHT Data Analysis Center.** The QC statuses are included in the File Manifest file, that are downloaded upon the files are selected (usually for downloads) in the SMaHT Data Portal (https://data.smaht.org). For more detailed descriptions of the QC checks and tools used, see Methods.

| **WGS file checked** | **QC check** | **Tool used** | **QC status when check failed** |
|---|---|---|---|
| FASTQ | Estimated % of microbial and viral contaminations.<br>PASS when <0.1% bacterial, <0.3% viral, and >95% human. | Kraken2 | FLAGGED |
| BAM/CRAM | Sample relatedness to check for duplicate sample or sample swap | Somalier | FAILED |
| | Estimated average coverage checked against target coverage | Mosdepth | FLAGGED |
| | Estimated % contamination from another human sample.<br>PASS when is <1%. | VerifyBamID2 | FLAGGED |
| | Mean insert size. PASS when >250 bp. | Picard | FLAGGED |
| | % of properly paired reads | Samtools | FLAGGED |
| | % of mapped reads | Samtools | FLAGGED |
| | % of duplicate reads | Samtools | FLAGGED |

**Table S6. Data access levels of the SMaHT benchmarking sequence data and derived data, including germline and somatic mutations, gene expression, and epigenetic profiles.** Authorizations for *post-mortem* donors within the main production phase of SMaHT allow for open access of most data, with the clinical metadata and germline sequence to remain under controlled access.

| | **COLO829, COLO829BL, COLO829BLT50** | **HapMap mixture** | **LB-LA2 fibroblasts and iPSCs** | **Tissues from *post-mortem* donors** |
|---|---|---|---|---|
| **Sequence data** (FASTQ, BAM/CRAM) | Open | Open | Protected | Protected |
| **Germline mutations** | Open | Open | Protected | Protected |
| **Somatic mutations:**<br>All mutation types from a cell line or individual donors as well as aggregated results from multiple donors | Open | Open | Protected | Protected |
| **Gene expression profile** | Open | Open | Protected | Protected |
| **Epigenetic profile** | Open | Open | Protected | Protected |

**Table S7. Advantages and disadvantages of DSA and pangenome-based somatic mutation detection.**

| Advantages | Disadvantages |
| --- | --- |
| Align reads accurately and improve somatic variant detection accuracy, especially in difficult and extreme regions | More complex analysis requires specialized knowledge and different data processing starting from alignment |
| Align reads and detect somatic variants in non-reference sequences (sequences missed in GRCh38) | Existing graph pangenomes clip highly repetitive sequences, limiting the overall genomic space represented. |
| Reduce germline contamination in alignments and somatic variant call sets | More computationally intensive and resource demanding (memory, storage) |
| | Pangenome is only an approximation of an individual's actual germline genome |

**Table S8. Remaining challenges in discovering somatic mutations.**

| Discovery mode | Challenge | Current state-of-the-art | Solution outlook |
|---|---|---|---|
| SNVs using bulk WGS based on the reference genome | Discovering in STR | Existing machine learning approaches (like MosaicForecast) perform well for coverages up to 300-400X | Novel machine learning and deep learning approaches trained for high coverage data |
| Indels using bulk WGS based on the reference genome | Discovering in homopolymers | Discovering indels with >3% VAF using assembly-based tools (like Lancet and Mutect2) | Using high fidelity sequencing (like provided by Elements) and incorporate platform specific error model into calling algorithms |
| MEIs using bulk WGS based on the reference genome | Accurate estimation of VAF at low levels (<3%) | MEI-targeted methods combined with ddPCR | Building models from higher-depth sequencing data |
| | MEI discovery in near-centromeric regions | Assembly-based alignment and tools | Leveraging T2T-level donor-specific assemblies |
| SVs from bulk WGS based on the reference genome | Discovering SV at VAF < 10% | Using long read sequencing | Increased coverage and improved analytics |
| | Adjacent SVs | Using long read sequencing and tools like Delly or Severus | Improved analytics |
| All mutation types using DSAs/graph genomes | Nascent computational tooling accounting for diploid nature of human genome | Somatic mutation discovery largely limited to linear reference genomes | Graph-genome and diploid-genome-based somatic mutation discovery methods |
| | Limited panel of germline variants | Pangenome graphs made from hundreds of diverse diploid assemblies | Larger diverse pangenomes with germline variant calls across large diverse human populations |
| SVs, CNVs, and MEIs using duplex sequencing approaches | False-positive chimeric reads and read mapping to repetitive genomic regions | Unresolved | New duplex sequencing approaches, including long read duplex approaches together with leveraging DSA |
| SNVs and Indels using duplex approaches | Distinguishing artifacts reaching single-strand consensus from true single-strand lesions | HiDEF-seq (SNVs); unresolved for indels | New approaches to model error rates based on sequence and coverage; model per-sample and per-technology errors and biases in sequence capture |
| | Accurately calling mutations beyond the traditional a4s2 requirement | DupCaller (Cheng et al., DOI: 10.1101/2025.07.13.664565.) | |
| | Variable input mass requirements due to different inefficiencies in library construction | Picogram-scale inputs; single-cell duplex sequencing | Approaches to experimentally pre-empt artifacts while preserving input DNA mass; filter out mismatches in data generated from low-input but error-prone solutions |

| | | | |
|---|---|---|---|
| All mutation types using PTA-amplified single cells (WGS) | Throughput and QC | Amplification via 96-well plate;<br>manual 4-loci PCR for the initial QC | Automated approaches for single cell/nuclei isolation and amplification;<br>SNP-based library-free amplicon sequencing QC |
| | Cost of sequencing | Sequence to at least 15× | Progressive reduction in sequencing cost |
| | Application to any post-mortem tissue | Application has been demonstrated for fresh cells and nuclei from post-mortem brain, heart, lung, and colon | Development of experimental protocols tailored for tissues and conditions (like cell density, disease, post-mortem interval) |

**Table S9. Details of the variants confirmed in the COLO829BLT50.**

| Variant | COSMIC ID | Bio-Rad Assay | VAF Tumor (Seattle) | VAF Tumor (BCM) | Expected VAF COLO829BLT 50 | Observed ddPCR VAF (Seattle) | Observed ddPCR VAF (BCM) |
|---|---|---|---|---|---|---|---|
| SCN11A p.T340T | COSM36715 | dHsaMDS432990804 | 49.38% | 50.1% | 1.00% | 1.44% | 1.41% |
| ZNF217 p.P651S | COSM25218 | dHsaMDS581392264 | 50.41% | 50.2% | 1.00% | 1.33% | 1.33% |
| MADD p.S1620F | COSM26934 | dHsaMDS851278352 | 99.91% | 100% | 2.00% | 1.79% | 1.64% |
| MAP4K1 p.P298P | COSM21033 | dHsaMDS546407091 | 32.70% | 33.2% | 0.83% | 0.76% | 0.71% |
| RELN p.E331K | COSM36731 | dHsaMDS550923980 | 49% | 48.9% | 1.64% | 1.53% | 1.40% |

**Table S10. Details of the variants confirmed in the HapMap sample mixture.**

| Sample ID | Expected AF | dbSNP ID | Chr. | Position (hg38) | Change | Observed AF (BCM) |
|---|---|---|---|---|---|---|
| **HG02622** | 10% | rs73825128 | chr3 | 28,336,081 | G>C | 15.18% |
| **HG002** | 2% | rs747651768 | chr6 | 157,128,737 | A>G | 1.8% |
| **HG02257** | 2% | rs199853723 | chr20 | 10,640,763 | T>G | 2.6% |
| **HG02486** | 2% | rs16961993 | chr17 | 17,768,088 | G>A | 4.5% |
| **HG00438** | 0.5% | rs149078332 | chr1 | 63,703,537 | G>A | 0.6% |
| **HG005** | 83.5% | rs2269756 | chr19 | 35,308,468 | C>A | 79% |

**Table S11**. **Details of the primers and probe designed to validate the variants in the HapMap sample mix.** Listed below are the sense (forward), anti-sense (reverse) as well as the wild-type (FAM) and variant (HEX) fluorescent TaqMan probes. For the probe designs the capital letters denote locked nucleic acid (LNA) bases.

| Primers/Probe | HG02622 | HG002 | HG02257 | HG02486 | HG00438 | HG005 |
|---|---|---|---|---|---|---|
| **Forward** | CACTGTCATGATGTCTGA | AGAGCGTGACCAATAGAA | GATGGCTTTATTGAATAGTATAATG | AGTGCTCAGTAAAGAGGA | ACAGCATTCTCAAACTTG | CCTTGTAGATGAAAGACTTTG |
| **Reverse** | CCTGAAGTCAACCACTAA | GCTGTCTTTACTTCCTTTTC | GCAGAAGTAAGAGTTCAGA | CCTCATATTACACCTGTGAC | GGCTTTTGTTCCTTAGTTA | CAGGGTCTTGCTATGTTC |
| **Wild Sense Dual-Labeled Probe** | ccaagCttGttAgaAcct | aagaGgcAaaCgaTata | tcacAcaAacTagTccca | ctacCctGagGatTgctg | tacaTtcTggGgtTttgt | tggcTtgCgcCttAtaa |
| **Mutant Sense Dual-Labeled Probe** | ccaagCttCttAgaAcct | aagaGgcGaaCgaTata | tcacAcaAacGagTccca | atggcaActAgcCaaTctc | tacaTtcTggAgtTttgt | tggcTtgAgcCttAtaa |
